# Transfer learning of an *in vivo-*derived senescence signature identifies conserved and tissue-specific senescence across species and diverse pathologies

**DOI:** 10.1101/2022.03.22.485297

**Authors:** Christopher Cherry, James I Andorko, Kavita Krishnan, Joscelyn C Mejias, Helen Hieu Nguyen, Katlin B Stivers, Elise F Gray-Gaillard, Anna Ruta, Naomi Hamada, Masakazu Hamada, Ines Sturmlechner, Shawn Trewartha, John H Michel, Locke Davenport Huyer, Matthew T Wolf, Ada Tam, Alexis N Peña, Claude Jordan Le Saux, Elana J Fertig, Darren J Baker, Franck Housseau, Jan M van Deursen, Drew M Pardoll, Jennifer H Elisseeff

## Abstract

Senescent cells (SnCs) contribute to normal tissue development and repair but accumulate with aging where they are implicated in a number of pathologies and diseases. Despite their pathological role and therapeutic interest, SnC phenotype and function *in vivo* remains unclear due to the challenges in identifying and isolating these rare cells. Here, we developed an *in vivo*-derived senescence gene expression signature using a model of the foreign body response (FBR) fibrosis in a *p16^Ink4a^-*reporter mouse, a cell cycle inhibitor commonly used to identify SnCs. We identified stromal cells (CD45^-^CD31^-^ CD29^+^) as the primary *p16^Ink4a^* expressing cell type in the FBR and collected the cells to produce a SnC transcriptomic signature with bulk RNA sequencing. To computationally identify SnCs in bulk and single-cell data sets across species and tissues, we used this signature with transfer learning to generate a SnC signature score (SenSig). We found senescent pericyte and cartilage-like fibroblasts in newly collected single cell RNAseq (scRNASeq) data sets of murine and human FBR suggesting populations associated with angiogenesis and secretion of fibrotic extracellular matrix, respectively. Application of the senescence signature to human scRNAseq data sets from idiopathic pulmonary fibrosis (IPF) and the basal cell carcinoma microenvironment identified both conserved and tissue-specific SnC phenotypes, including epithelial-derived basaloid and endothelial cells. In a wound healing model, ligand-receptor signaling prediction identified putative interactions between SnC SASP and myeloid cells that were validated by immunofluorescent staining and *in vitro* coculture of SnCs and macrophages. Collectively, we have found that our SenSig transfer learning strategy from an *in vivo* signature outperforms *in vitro*-derived signatures and identifies conserved and tissue-specific SnCs and their SASP, independent of *p16^Ink4a^* expression, and may be broadly applied to elucidate SnC identity and function *in vivo*.

---

SnCs are implicated in multiple age-related pathologies including heart disease, diabetes, neurodegeneration, cancer, and arthritis (Campisi, 2013; Childs, Li, & Van Deursen, 2018; Faust et al., 2020; Howcroft et al., 2013; Jeon et al., 2017; Jeon et al., 2019; Minamino et al., 2009). Despite their possible central role in these conditions, and as potential targets for therapeutic intervention, the phenotype and function of SnCs *in vivo* remains elusive due to challenges with identification and isolation of these cells. Moreover, the presence of SnCs during embryological development and their beneficial contributions to wound healing suggest heterogeneity in phenotype and function, further amplifying the need for understanding the diverse context-dependent senescence phenotypes that define beneficial versus pathologic senescence (Li et al., 2018; Muñoz-Espín et al., 2013; Storer et al., 2013). Developing an in depth understanding of SnCs in health and disease will enable more efficient therapeutical targeting of these cells and their products.

SnCs are non-proliferative and identified by markers including the cell cycle gate keeper p16^INK4A^ and the expression of a senescence associated secretory phenotype (SASP). SnCs contribute to tissue physiology through the SASP which includes many immunologically-relevant molecules such as Interleukin(IL)-6, IL1-β, and CCL2 (Coppé et al., 2008; Tchkonia, Zhu, Van Deursen, Campisi, & Kirkland, 2013; Wan, Gray-Gaillard, & Elisseeff, 2021). To date, phenotypical characterization of SnCs primarily relied on methods to induce senescence *in vitro* such as high dose gamma irradiation, oncogene induction, drug-induced oxidative damage and proliferative senescence (Bai, Gao, Hoyle, Cheng, & Wang, 2017; Dumont et al., 2000; Hooten & Evans, 2017). While SnCs and their impact on tissue physiology can be identified *in vivo* by immunofluorescence and using transgenic animals, there are no widely applicable markers to effectively and comprehensively isolate the cells for further identification and phenotypic analysis including SASP expression (Amor et al., 2020; Kim et al., 2017; Poblocka et al., 2021). The emerging wealth of single-cell atlases across healthy tissues and disease provide a plethora of data in which to potentially evaluate the role of SnCs across different tissue contexts, however an accurate SnC signature is required to identify the cells and leverage these resources.

Single cell RNA Sequencing (scRNASeq) and other single cell “omics” technologies are enabling the development of cell atlases across tissues and pathologies. The technology also allows for discovery of intercellular interactions, a feature which is appealing for the study of a cell population which is canonically active largely through secretion of signaling proteins. For example, single cell analyses illuminated heterogeneity in fibroblasts, a cell population frequently used to study senescence *in vitro* and suggested to be senescent *in vivo* (Chung et al., 2020; Demaria et al., 2014). Notably, Buechler at al., recently created an atlas of fibroblasts across multiple tissues and identified common and pathology-specific fibroblasts (Buechler et al., 2021). In the case of arthritis, Wei et al., identified multiple fibroblast populations that contribute to disease pathology and were activated by NOTCH3 signaling (Wei et al., 2020). In cancer, Elyada et al., characterized two subtypes of cancer associated fibroblasts (CAFs) using scRNAseq, inflammatory- and myo-fibroblasts, and their respective secreted signaling molecules contributing to tumor behavior and therapeutic response (Elyada et al., 2019).

The advances in single cell -omics technologies have not yet led to the same breakthroughs in the senescence field. SnCs, a relatively rare cell population, cannot be identified with canonical markers from scRNAseq data sets. This is likely because high-throughput, microfluidic based scRNAseq methods only capture ∼10% of the transcripts in a given cell, leading to dropout where genes with lower expression levels cannot be identified (Hwang, Lee, & Bang, 2018). This limits the ability of scRNAseq to identify SnCs using the common marker *Cdkn2a,* the gene that encodes p16^INK4A^, due to its low levels of expression in a small number of cells. Transfer learning analyses, which can leverage small expression changes across hundreds of genes, may provide an opportunity to identify SnC in scRNAseq data sets despite dropout and have been shown to robustly identify preserved biological processes across technologies, tissues, diseases, and species (Stein-O’Brien et al., 2019; Taroni et al., 2019). These techniques provide the potential to identify SnCs based upon gene signatures in place of isolated gene or protein markers to query their phenotype and function in atlas data sets from diverse biomedical contexts.

Here, we present a new approach for identifying physiologically-relevant SnCs *in vivo* across tissues, conditions, and species. We first used a novel *p16^ink4a^*-marked tdTomato (tdTom) reporter mouse to collect and characterize SnCs that develop during the foreign body response (FBR) fibrosis to develop a comprehensive *in vivo* gene expression signature. We then used a transfer learning technique to score mouse and human scRNAseq data sets for concordance with the expression signature. The technique, which we term SenSig, also allows for identification of genes responsible for increased SenSig in a given population of cells. Application of the SenSig method to mouse and human FBR single cell data sets defined two populations of senescent stromal cells: pericytes and a cartilage-like fibroblast involved in fibrosis. The SenSig method also functioned in public human-derived data sets of idiopathic pulmonary fibrosis (IPF) and basal cell carcinoma, confirming broad applicability and elements of a conserved signature within the murine FBR SnC expression signature. Subsequent intercellular signaling analysis found specific signaling patterns through which SnCs communicate with myeloid cells during wound healing, findings which were validated by *in vitro* coculture experiments. Overall, the SenSig identifies a conserved transcriptional profile of SnCs that may be broadly applied to identify and understand senescence across a range of diseases. The ability to identify SnCs in the multitude of singe cell and even bulk data sets that are publicly available will enable understanding of senescence phenotype, conserved and tissue-specific SASPs, and subsequent interactions with other cell types such as those from the immune system to define their contribution to tissue homeostasis and pathology.

## Generation of a physiological senescence signature from the foreign body response

Our first goal towards creating a senescence signature was phenotypic identification and characterization of SnCs *in vivo*. We previously demonstrated that SnCs develop during the FBR to a biomaterial implant in both murine models and clinical samples (Chung et al., 2020). Similarly, we implanted polymeric particles composed of poly(caprolactone), PCL, to induce fibrosis in a muscle wound to enrich the number of SnCs (Fig. 1A and B). *p16^ink4a^* expression, a classical marker for senescence, increased significantly in the muscle tissue one week after injury with or without an implant confirming wound-induced senescence (Fig. 1C). After 6 weeks, when fibrosis around the implant matures, *Cdkn2a* expression significantly increased in the whole tissue containing the PCL implant compared to the wound alone (Fig. 1C). Sorted stromal cells (CD45^-^CD31^-^CD29^+^) from the muscle tissue increased *Cdkn2a* expression over 10-fold at both 1 and 6 weeks, suggesting the stromal compartment represents a significant source of the SnCs in the healing muscle tissue and FBR fibrosis (Fig. 1D).

**Figure 1.**
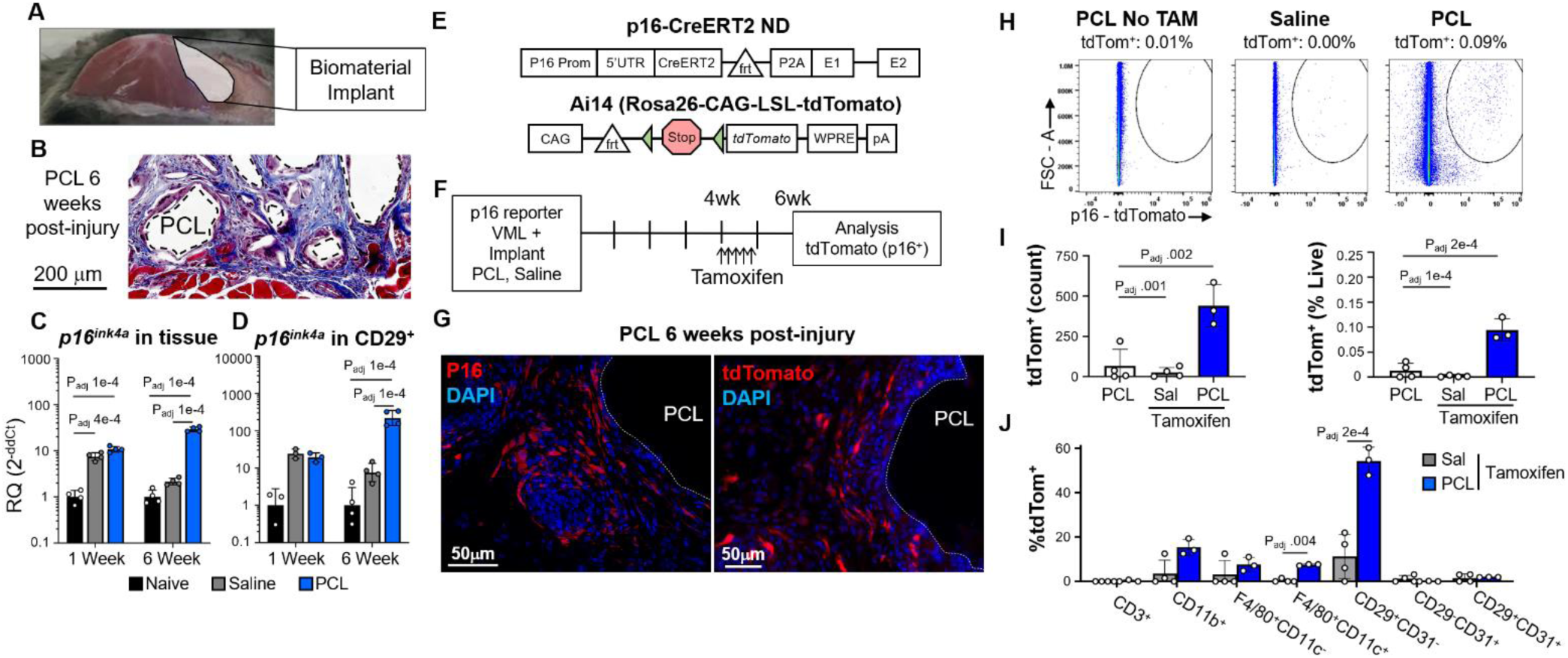
Cre-transgenic mouse identifies p16+ SnCs in fibrotic tissue from the foreign body response of biomaterial implant. (A) Diagram of the volumetric muscle loss (VML) surgery. After surgical excision of a portion of quadriceps, a biomaterial implant, (poly)caprolactone (PCL), or an equivalent volume of saline are implanted into the tissue space. (B) Representative Masson’s trichrome staining of fibrosis around PCL implant particles after 6 weeks of implantation, scale bar = 200 µm. (C) qPCR for *p16^ink4a^*, encoding the protein p16, of whole muscle tissue implanted with PCL and control tissue treated with saline and naïve/no surgery collected 1- or 6-weeks post-surgery. Statistics shown are from ANOVA followed by multiple t-test with Benjamini-Hochberg correction for multiple testing. Adjusted P-values for all statistically significant (P_adj_ < .05) comparisons are shown. (D) qPCR for *Cdkn2a*in CD29+CD45-CD31-cells sorted by fluorescence-activated cell sorting (FACS). Statistics shown are from ANOVA followed by multiple t-test with Benjamini-Hochberg correction for multiple testing. Adjusted P-values for all statistically significant (P_adj_ < .05) comparisons are shown. (E) Schema for transgenic p16 reporter mice*. CreER^T2^* is inserted inside the *p16-*specific exon 1 of the *Cdkn2a* gene. After Tamoxifen administration, Cre recombines loxP sites (triangles) of the *Ai14* reporter construct resulting in excision of the stop cassette and subsequent functional expression of *tdTom*. (F) Treatment schematic for induction of tdTom fluorescent signal in p16+ cells in *p16-CreER^T2^;Ai14* mice. Mice were treated with tamoxifen injections daily for 5 days starting 4 weeks post surgery to induce permanent tdTom expression at the 6-week harvest. (G) Immunostaining for p16 in WT animals and native tdTom expression in *p16-CreER^T2^;Ai14* transgenic mice treated with PCL implants six weeks post-surgery, scale bar = 50 µm. (H-J) Flow cytometric measurement of tdTom+ cells from animals with VML treated with saline or PCL with or without tamoxifen. (H) Representative scatter plots for live, tdTom+ cells, (I) quantification of cell frequency and percent of total live cells, and (J) numbers of tdTom+ cells in various cellular subsets in animals given VML with saline or PCL with tamoxifen are shown. Statistics shown are from ANOVA followed by multiple t-test with Benjamini-Hochberg correction for multiple testing. Adjusted P-values for all statistically significant (P_adj_ < .05) comparisons are shown.

To further investigate which cell types became senescent in the fibrotic capsule, we utilized an inducible *p16*-dependent reporter mouse model (Fig.1E) (Hamada et al., bioRxiv submission pending). In this model, the endogenous *p16* promoter drives the expression of a tamoxifen-activated form of Cre recombinase (CreER^T2^), but without exogenously added elements that enhance transcriptional activity (Omori et al., 2020). These mice were bred to a reporter strain that harbors an Ai14 cassette inserted in the *Rosa26* locus with an upstream loxP flanked stop cassette that does not allow constitutive tdTom expression (Fig. 1E). With tamoxifen exposure, Cre becomes active allowing for deletion of the stop cassette and subsequent transcription of tdTom. In this way, only cells expressing *p16* within a specified timeframe, defined by the duration of tamoxifen exposure, will be positive for tdTom. This system also allows detection and collection of viable SnCs. Homozygote *p16-CreER^T2^;Ai14* mice still expressed small amounts of p16 protein due to the presence of a viral 2A sequence, and underwent senescence as measured by growth arrest after high dose γ-irradiation and expression of SASP factors *Il6*, *Mmp3*, and *Ccl2* (Supplementary Fig. 1 and 2).

To identify SnCs associated specifically with fibrosis around the biomaterial, we delivered tamoxifen 4 weeks post implantation after the primary trauma-related senescence resolved. At 6 weeks, tdTom fluorescence was visible around the PCL implant with a similar distribution as p16 protein immunofluorescence in wildtype animals (Fig 1G). Flow cytometric analysis confirmed that the presence of a PCL implant significantly increased tdTom^+^ cells with few to no cells present in animals not receiving tamoxifen or in saline treated mice without a biomaterial (Fig. 1H and I). Using a pan immune-stromal flow cytometry panel we could further determine the cell type that expressed tdTom (Fig. 1J and Supplementary Fig. 3). While less than 0.1% (+/- 0.01%) of cells in the tissue were tdTom^+^, 54% (+/- 3.6%) of these tdTom^+^ cells were CD45^-^CD31^-^CD29^+^ stromal cells. Meanwhile, 15% (+/-2.1%) of the tdTom^+^ cells were CD45^+^CD11b^+^ myeloid cells. Of the myeloid cells, F4/80^+^CD11c^+^ scaffold-associated macrophages and F4/80^+^CD11c^-^ macrophages accounted for approximately half of the tdTom^+^ cells (7.5% +/- 0.2% and 7.6% +/- 1.8% respectively). A small number of myeloid cells expressed tdTom at significantly increased levels, however we previously demonstrated that sorted CD11b^+^F4/80^+^ macrophages in this model did not express *p16* (Chung et al., 2020) suggesting that the tdTom signal may be due in part to macrophage engulfment of SnCs or SnC-derived exosomes (Chung et al., 2020; Sturmlechner et al., 2021). Since the majority of the tdTom^+^ cells in the FBR fibrosis were CD45^-^CD31^-^CD29^+^ stromal cells, we used this population to generate a senescence gene expression signature.

## Angiogenesis and cartilage pathways are activated in tdTom+ senescent stromal cells

The significant enrichment of SnCs in the biomaterial-induced fibrosis model enables isolation of tdTom-expressing cells. To identify the phenotype of the stromal tdTom^+^ cells in the FBR, we sorted CD45^-^CD31^-^CD29^+^ tdTom^+^ cells and the tdTom^-^ counterpart and performed bulk RNA sequencing. Expression of both the *tdTom* transgene and *Cdkn2a* were elevated in the tdTom/p16^+^ cell population compared to the tdTom^-^/p16^-^ cells confirming that we isolated stromal SnCs (Fig. 2A). Furthermore, the tdTom^+^ cells expressed higher levels of multiple genes associated with senescence including *Mmp13*, *Mmp3*, and *Serpinb2* (Childs et al., 2016; Wiley et al., 2019). Of the genes classically related to SnC that we investigated, only *Il6* was not increased in the sorted tdTom^+^ stromal cells. Beyond the recognized SnC genes, broader differential expression identified 1803 genes with increased or decreased expression characteristics in the tdTom^+^ versus tdTom^-^ sorted cell populations (2B and C and Supplementary Table 1, FDR < .05).

**Figure 2.**
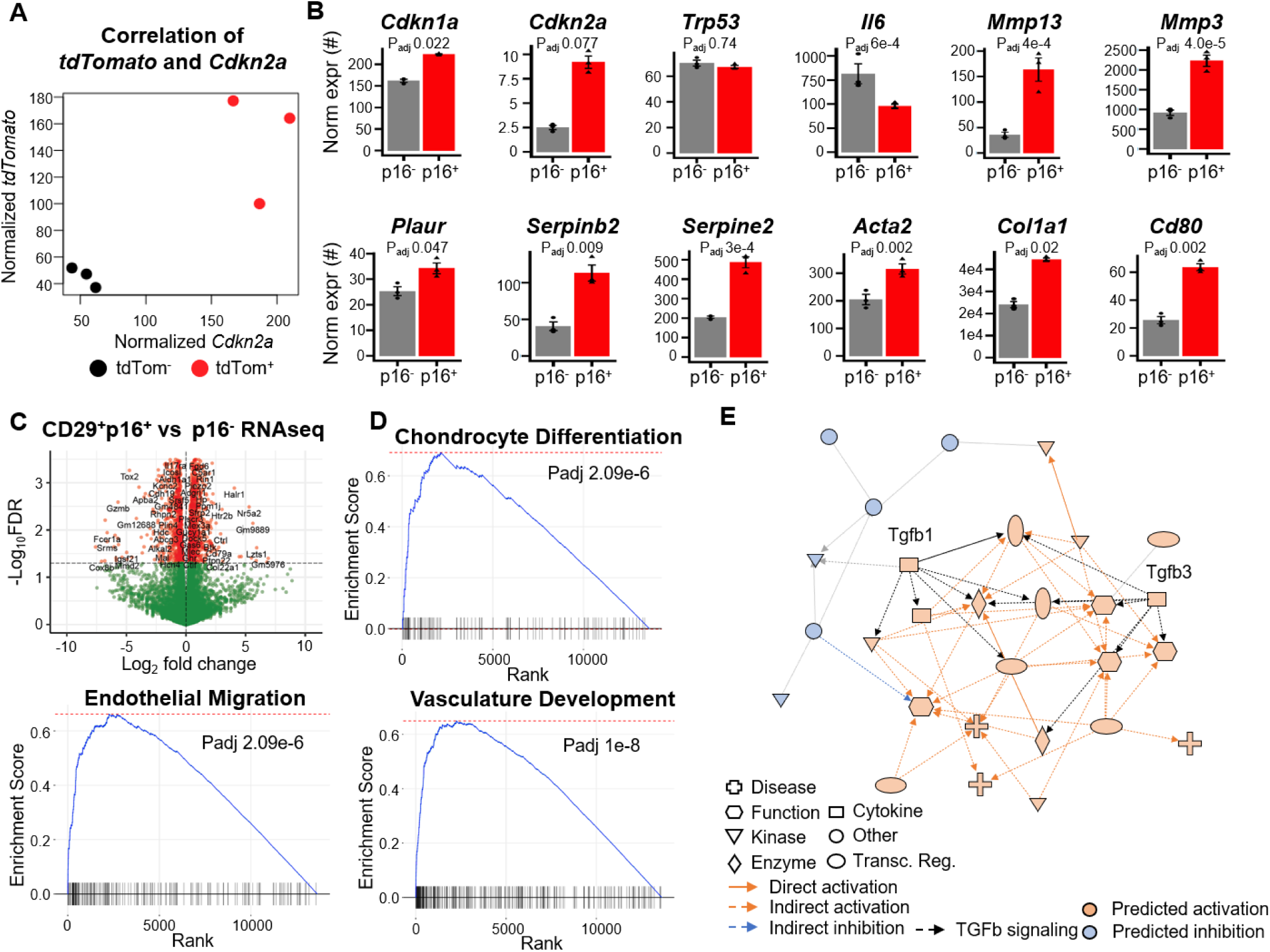
RNA sequencing identifies a senescence signature from stromal cells in the foreign body response. (A and B) Gene expression from bulk RNA sequencing comparing sorted p16+CD29+CD45-CD31- and p16-CD29+CD45-CD31-populations. Correlation of normalized *tdTom* and *Cdkn2a* by sample (A) and normalized gene expression for common senescence associated genes (B) are shown. Statistics shown are derived from a negative-binomial test using edgeR followed by FDR correction for multiple testing. (C) Volcano plot visualizing differentially expressed genes between p16+ and p16-sorted populations. Statistics shown are from a negative binomial test under edgeR with FDR correction for multiple testing. A threshold of 0.05 is used to select significant genes. Positive fold-change indicates increased expression in p16+ cells. (D) Gene set enrichment analysis (GSEA) for gene ontology biological process annotations. Positive enrichment scores indicate increased expression in the gene set in the p16+ cells with increased scores indicating increasingly unlikely enrichment under the assumption of random gene order. Statistics reported are from GSEA with FDR multiple testing correction. (E) Ingenuity pathways analysis upstream regulatory motifs from the bulk RNA sequencing data. The network indicates upstream regulators predicted to be potential modulators of the gene expression changes detected from bulk RNA sequencing between the p16+ and p16-cells. Tgfb1 and Tgfb3 transcription factors are noted and their outgoing regulatory connections indicated in black.

To characterize the nature of expression differences in the senescent versus non-senescent stromal cells, we used gene set enrichment analysis with gene ontology annotations (Fig. 2D and Supplementary Tables 2). Gene set enrichment analysis (GSEA) identified a number of gene sets associated with either vasculature development or bone/cartilage formation upregulated in the p16^+^ stromal populations. Upregulation of cartilage and bone related gene sets were driven by extracellular matrix proteins *Ccn3*, *Ccn2*, *Col11a1*, and *Col20a1* as well as growth factors *Bmp2/4/6*, *Fgf18*, and *Tgfb1*. In contrast, the vasculature associated gene sets were driven by genes *Tek*, *Anxa3*, *Sparc*, *Vegfa*, *Vegfc*, and *Myh9*, all genes associated with promotion of vascularization (Supplementary Fig. 4 and Supplementary Table 3). These distinct signatures may represent two senescent stromal cell populations in the FBR fibrosis, similar to those previously observed in skin wound healing (Demaria et al., 2014). Lastly, we used Ingenuity Pathways Analysis (IPA) to probe potential regulatory signatures involved in the senescence phenotype (Fig. 2E). IPA identified transforming growth factor beta (TGFβ) as the primary upstream regulator of the differentially expressed genes in the senescent stromal cells. TGFβ has been widely implicated in fibrosis in multiple pathologies (Frangogiannis, 2020; Rice et al., 2015; Sanderson et al., 1995; Sime, Xing, Graham, Csaky, & Gauldie, 1997; Sonnylal et al., 2007).

## Single cell RNA sequencing of stromal cells identifies clusters of the FBR

Our next goal was to broadly identify the composition and heterogeneity of the stroma in the FBR fibrosis model for subseqent correlation with the SnC phenotypes identified in the *p16* reporter model. We performed Drop-Seq (Macosko et al., 2015) on CD45^-^CD31^-^CD29^+^ cells isolated from C57BL/6 (wild type) mice with biomaterial implants, control wounds (without an implant), as well as native muscle tissue at one- and six-weeks post-surgery (Fig. 3A). The early (one week) time point and the tissue injury without biomaterial data sets provides information on normal wound healing compared to the later biomaterial implant condition where fibrosis, inflammation, and senescence is enriched. After quality control and thresholding on total UMI count, total feature count, and contribution of mitochondrial genes to total UMI count, the samples were integrated using Harmony. Unsupervised clustering on the resulting combined data set identified 16 distinct stromal clusters characterized by unique gene expression signatures (Fig. 3B, Supplementary Fig. 5A, and Supplementary Table 4).

**Figure 3.**
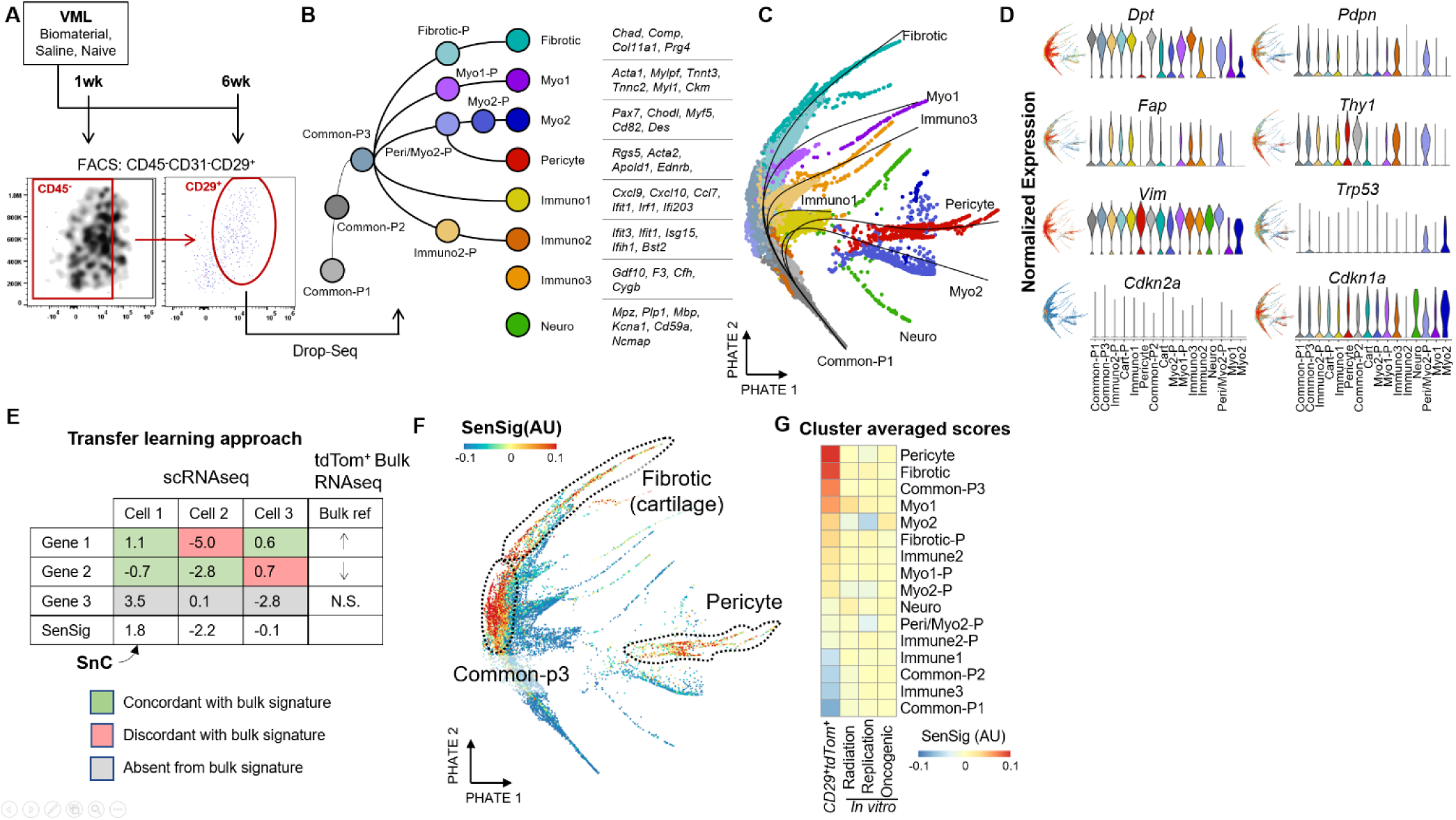
Transfer learning of senescence signature identifies SnCs in single cell sequencing of the murine synthetic implant microenvironment. (A) Experimental schema and FACS sorting for single cell RNA sequencing. Animals were given VML and treated with biomaterial implants, saline, or were naive to surgery. CD29^+^CD45^-^CD31^-^ cells were sorted one- and six- weeks post-surgery before input into the Drop-seq scRNAseq platform. (B) scRNAseq data set overview. Clusters are labeled and ordered according to their predicted pseudotime trajectory. Characteristic genes for terminal clusters are shown. (C) PHATE dimensional reduction visualization of clusters with psuedotime overlays with terminal clusters labeled. (D) Expression of fibroblast marker genes (top) and senescent markers *Cdkn1a* and *Cdkn2a* (bottom). Both feature plots showing localization of log-normalized gene expression on the PHATE dimensionality reduction (left) and violin plots (right) are shown. (E) Transfer learning to identify putative SnC in the scRNAseq data set. A scoring methodology based on z-scored gene expression was used to calculate senescent signatures (SenSig) based on the set of genes differentially expressed in the bulk RNA sequencing comparison of P16^+^ and P16^-^CD29^+^CD45^-^ CD31^-^ cells. (F) Calculated SenSig by cells arranged by PHATE. The three clusters with highest average SenSig are highlighted. (G) Cluster averaged SenSig derived from the p16 reporter mice as well as from three publicly available senescence signatures from *in vitro* bulk RNA sequencing.

We first investigated differentiation pathways in the clusters obtained from the scRNASeq analysis. Embedding of cells into a two-dimensional space using PHATE (Moon et al., 2019), an algorithm based on diffusion mapping which preserves global relationships between cells better than uniform manifold approximation and projection (UMAP) (Becht et al., 2019), showed multiple branching arms. These arms indicate increasingly differentiated expression states when compared to the other stromal cells, possibly as a result of cellular differentiation (Fig. 3C). CytoTRACE (Gulati et al., 2020), an algorithm which scores scRNAseq data for stemness, identified stem-like clusters which we used in combination with RNA velocity (La Manno et al., 2018) and pseudotime (Street et al., 2018) analysis to identify six separate stromal cell differentiation trajectories corresponding to the arms from the PHATE projection (Supplementary Fig. 5B and C). In combination with two cell populations which did not connect via pseudotime with the other subsets, 8 total terminally differentiated cell populations were identified. The remaining 8 clusters of cells were classified as intermediate or progenitor populations along these trajectories. Three of the terminal populations were primarily characterized by expression of inflammatory or signaling molecules (Immuno1, Immuno2, and Immuno3). The other five terminally differentiated cell clusters were characterized by genes associated with tissue formation (Fibrotic/Cartilage, Myo1, Myo2, Pericyte, Neuro).

Commonly used fibroblast markers did not appear to resolve the heterogeneity of clusters in our single cell data set (Fig. 3D). The majority of common fibroblast surface markers like *Thy1*, *Fap*, *Dpt, Vim*, and *Pdpn* (Buechler et al., 2021; Cheng et al., 2016; Kahounová et al., 2018; Muhl et al., 2020; Schmidt et al., 2015) were highly expressed across multiple phenotypically distinct populations, suggesting that traditional fibroblast surface markers are not adequate to resolve the phenotypic diversity seen in the FBR scRNAseq data set. Additionally, we mapped previously collated single cell atlases (Buechler et al., 2021) of fibroblasts from naïve or perturbed mice to our data set (Supplementary Fig. 6). While most of our clusters mapped to the atlases, many of the distinct phenotypes were pooled more generally. For example, the myogenic, neuro, and pericyte clusters all mapped to stem-like progenitor populations. The majority of the other FBR fibroblast populations mapped to a broad fibroblast cluster define d as adventitia.

The fibrotic cluster from the FBR data set mapped exclusively to an atlas cluster labelled tendon/ligament. To validate cartilage-like fibroblasts in the FBR scRNASeq data set were not tendon or ligament from the muscle tissue, we performed fluorescent *in situ* hybridization (FISH) examining expression *Cdkn2a*, *Cd29* (*Itgb1*), and the cartilage-related marker *Fmod* in the fibrosis surrounding the PCL implants (6 weeks post-surgery, Supplementary Fig. 7 and 8). We found expression of *Fmod* exclusively in the fibrosis area surrounding PCL implants and not in healthy tissue, confirming that contaminating tendon or ligament was not responsible for the cartilage-like population.

Despite the known enrichment of p16^+^ cells in the wound and fibrosis environment determined by qPCR and flow cytometry (Fig. 1), none of the scRNASeq clusters showed distinct expression of senescence markers (Fig. 3D). Expression of *Cdkn2a* was sparse, likely due to dropout, a phenomenon in scRNAseq where low-expression transcripts are lost. In contrast, expression of *Cdkn1a*, another senescence marker gene, was found across most of the stromal populations in the data set.

## Transfer learning with the SenSig identifies putative SnC clusters in stromal scRNASeq

To identify SnCs in the scRNAseq, we developed a transfer learning approach that maps the SnC signature from bulk RNAseq of the transgenic p16^+^ reporter mice to the scRNAseq data set (Fig. 3E). This approach uses a broad set of genes to identify SnC instead of a small number of marker genes, allowing for identification of SnC even if detection of marker genes (like *p16^ink4a^*) is not found. We first select the differentially expressed genes from bulk RNAseq of the tdTom^+^ vs tdTom^-^ cells (FDR < .05) in the scRNASeq data set, then we identify overlapping genes present in the scRNAseq data set, and lastly sum the z-scored expression data by cell that is weighted by concordance with the direction of the fold-change from bulk RNAseq. Genes expressed in a concordant direction increased score while those expressed in a discordant direction decreased the score. The resulting score that we term SenSig (senescence signature) identifies cells with similar expression patterns to the sorted p16^+^ cells from the transgenic reporter model. Cells with a high SenSig represent putative SnCs and clusters enriched with high SenSig cells identifies the cellular phenotypes likely enriched for SnCs.

We found the *in vivo* derived SenSig enriched in cells from three clusters in the FBR stromal scRNAseq data set; pericytes, fibrotic fibroblasts, and a common progenitor to both (Fig. 3F). The pericyte cluster was defined primarily by common pericyte marker genes *Rgs5*, *Myh11*, and *Acta2* (Brandt et al., 2019; Kumar et al., 2017; Mitchell, Bradley, Robinson, Shima, & Ng, 2008). The fibrotic (cartilage-like) fibroblast cluster was defined by secreted extracellular matrix proteins like *Fmod, Chad, Comp,* and *Col11a1,* many of which are also expressed by chondrocytes. Three *in vitro-*derived senescence signatures from the literature (Hernandez-Segura et al., 2017) (irradiation, replicative, and oncogene induced senescence) did not map strongly to the scRNAseq cells when compared to the *in vivo* derived signature (Fig. 3G), suggesting that physiological senescence differs significantly from *in vitro* models.

The contribution of genes driving similarity to senescence and a high SenSig can be ranked by cluster (Supplementary Fig. 9). The top SenSig driver genes for the pericyte population were largely associated with vascular function like *Myh11*, *Acta2*, and *Apold1*. Interestingly, *Notch3* was also elevated in a pericyte cluster that was implicated in spatial and transcriptional control of fibroblasts in the synovium of inflammatory arthritis (Wei et al., 2020). In contrast, the genes driving senescence similarity score in the fibrotic population were generally extracellular matrix proteins or glycoproteins like *Col11a1*, *Col12a1*, *Thbs4*, and *Ccn2*, the latter of which has been implicated in induction of senescence (Joon-II Jun & Lau, 2017). This reinforces the association of a senescent, cartilage-like fibroblast population that secretes ECM proteins, supported by correlation with tendon and ligament in the fibroblast atlas, and these molecules contribute to pathological fibrosis.

To determine if the transfer learning could identify senescence associated with normal wound healing versus pathological fibrosis, we investigated how SenSig scores varied by sample condition and timepoint with the two pericyte and fibrotic clusters (Supplementary Fig. 10). SnC involved in the normal wound healing process peak around 1 week post injury in both saline and PCL treated mice as demonstrated by whole tissue expression of *p16* (Fig. 1C). In contrast, fibrosis associated SnCs increases at 6 weeks post injury in only PCL treated animals when extensive fibrosis is observed. SenSig high cells were distributed across all conditions and timepoints indicating that SnCs contributing to both normal wound healing and pathological fibrosis can be identified with the bulk-derived senescence signature. Finally, we sought to investigate whether *Il6*, a well-recognized SASP component that was not significantly upregulated in the bulk RNAseq, differentially regulated in our data set. While *Il6* was lower in the fibrotic fibroblast cluster, it was higher in the pericyte cluster (Supplementary Fig. 11A and B), indicating the importance of resolving the cellular phenotypes to accurately capture phenotype, understand SnC heterogeneity and specific SnC contributions to the local tissue environment.

## Cross-species SnC identification from single cell RNA sequencing with transfer learning

To determine if similar senescent populations were present in human models of the FBR, we collected tissue surrounding surgically excised synthetic breast implants that reproducibly induce fibrosis and demonstrate high levels of SnCs (Fig. 4A). We processed a single cell suspension enriched for CD45^+^ immune cells with the Drop-seq scRNAseq platform. After clustering, we computationally isolated stromal cells based on expression of *CD29* (*ITGB1*) and absence of *CD45* (*PTPRC*) and *CD31* (*PECAM1*) (Fig. 4B and C). We re-processed the stromal cells to generate 6 stromal clusters and then calculated a SenSig by translating the murine gene signature to human equivalent genes based on Ensembl orthologue mapping(Howe et al., 2021) (Fig. 4D and E).

**Figure 4.**
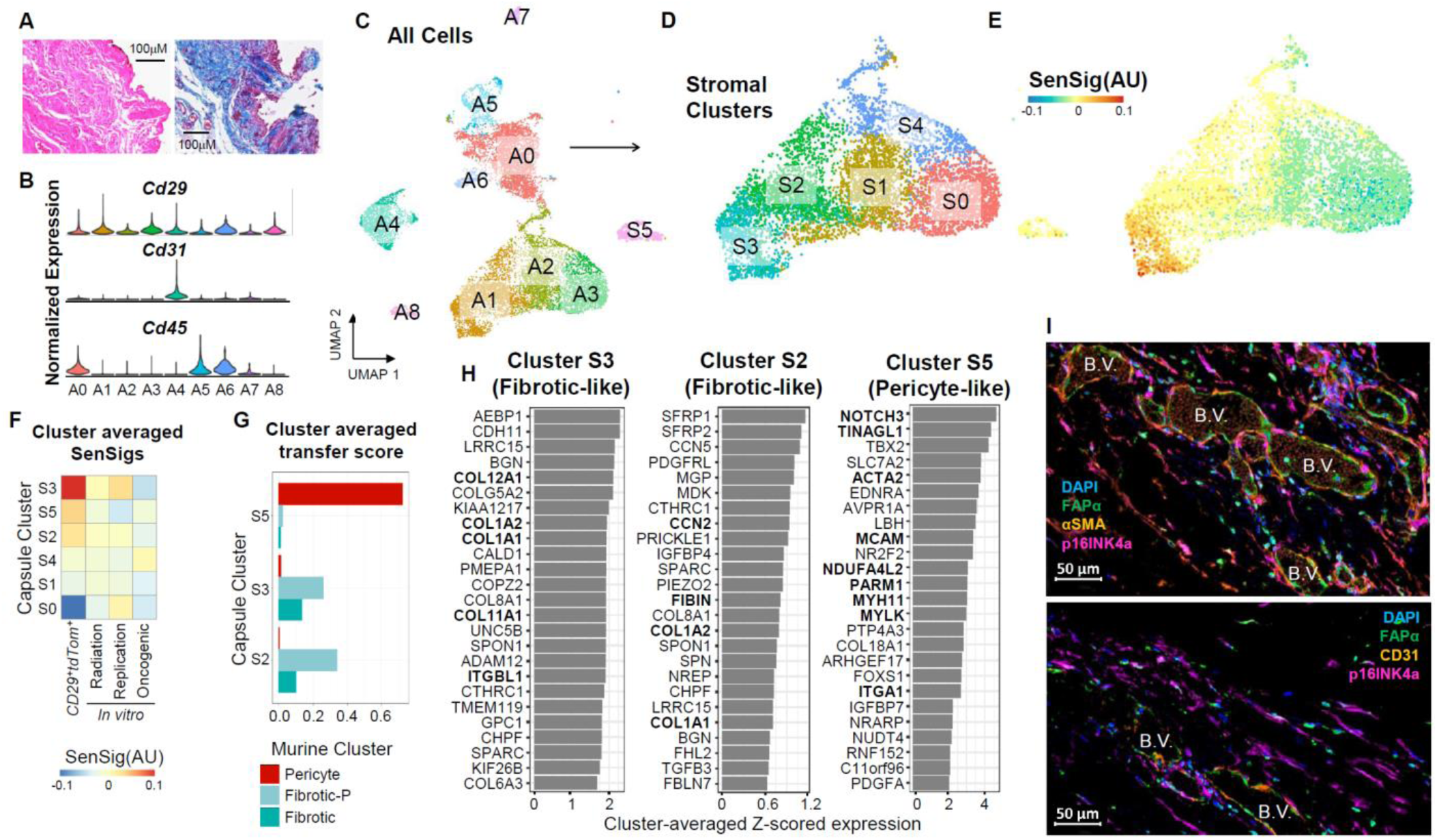
Mouse to human transfer learning of the senescent signature in single cell sequencing data set from the fibrotic capsule of a human breast implant. (A) Histological staining of surgically removed fibrotic capsules surrounding synthetic breast implants. Hematoxylin and Eosin (left) and Masson’s trichrome (right) staining are shown demonstrating fibrosis of the breast capsule tissue. (B) Expression of stromal cell markers by cluster from a scRNAseq data set collected from surgically removed synthetic breast implants. CD45+ immune cells were enriched to 50% of the total population prior to scRNAseq library generation using the Drop-seq protocol. (C) Visualization of the scRNAseq clusters on UMAP before subsetting to stromal populations. (D) Stromal cells from the scRNAseq data set after computational isolate of CD45-CD31-CD29+ cells and subsequent re-clustering of the stromal cells. (E) Feature plot visualization of SenSig in stromal cells derived as described in Figure 3E. (F) SenSig averaged by stromal cluster using our *in vivo* derived SnC gene set as well as three publicly available *in vitro* derived senescent gene sets. (G) Cluster level similarity scores to the murine putative SnC clusters using singleCellNet. Higher similarity score indicates similar gene expression patterns to the indicated murine scRNAseq clusters. (H) Genes driving the largest increase in SenSig in the three human stromal clusters with highest average SenSig. Values shown are average z-scored expression across cells in the target cluster. Genes that were also in the top 25 genes driving senescent signature in the murine fibrotic or pericyte clusters are shown in bold. (I) Fluorescent staining for fibroblasts (FAPα), endothelial cells (CD31), smooth muscle actin (αSMA), and SnC (p16INK4a) in fibrotic tissue capsules surrounding surgically removed synthetic breast implants.

Three of the six stromal clusters from the human capsule stromal populations had moderate or elevated SenSig (Fig. 4F). After comparing broad transcriptional signatures of the SenSig high human clusters to the murine stromal clusters, two of these clusters shared transcriptional similarities with the murine fibrotic fibroblast cluster and the third was similar to the murine pericyte cluster (Fig. 4G and Supplementary Fig. 12). Investigation of the genes most responsible for the elevated SenSig by cluster showed distinct modules of genes by cluster (Fig. 4H). The fibrotic, fibroblast-like human clusters were strongly elevated for matricellular proteins *CCN5* and *CCN2*, collagens *COL81A* and *COL1A1*, and *TGFB3*, although interestingly each fibrotic fibroblast-like capsule cluster appeared to have a distinct expression program driving their SenSig respective to each other. The genes driving high SenSig in the human pericyte-like cluster overlapped with numerous genes driving SenSig in the murine pericyte cluster, including *NOTCH3*, *MYH11,* and *MYLK*. Additionally, both the fibrotic fibroblast- and pericyte-like clusters shared a number of SenSig driver genes with their respective murine clusters. Most similar driver genes between the human and murine fibrotic fibroblast-like clusters were fibrotic extracellular matrix proteins like *COL1A1, COL1A2, COL11A1,* and *CCN2* while similar drive genes in the pericyte-like cluster were smooth muscle extracellular matrix proteins like *ACTA2*, *MYH11*, and *MYLK,* although *NOTCH3* was also a top SenSig driver gene.

Finally, to validate the presence of SnCs in the fibrotic breast capsule, we performed immunofluorescent staining for p16 in conjunction with stromal (FAPα, αSMA) and endothelial (CD31) cell markers (Fig. 4I). We observed co-staining of αSMA and p16 both adjacent to and distanced from CD31^+^ blood vessels. Importantly, we did not observe costaining of FAPα and p16, indicating concordance with the scRNAseq data that the senescent populations are not FAPα^+^. These findings suggests that although the SenSig is derived from murine bulk RNA sequencing data, it is sufficient to identify putative SnCs in the human fibrotic environment.

## SenSig identifies distinct senescent phenotypes in human idiopathic pulmonary fibrosis and cancer

We then sought to determine if our transfer learning approach would identify similar senescent populations in different human pathologies where senescence is thought to play a detrimental role. We used two publicly available data sets of idiopathic pulmonary fibrosis (IPF), a disease involving scarring and fibrosis of the lungs independent of any wound or synthetic implant (Adams et al., 2020; Habermann et al., 2020). Starting from the previously reported clustering from each data set, we calculated SenSigs for each data set and determined putative senescent clusters (Fig. 5A and B and Supplementary Fig. 13A and B). In both data sets, we identified similar clusters with high SenSig and distinct phenotypes, including a KRT5-/KRT17+ cluster, which had previously been proposed to be senescent based on expression of SASP factors and growth arrest (Habermann et al., 2020).

**Figure 5.**
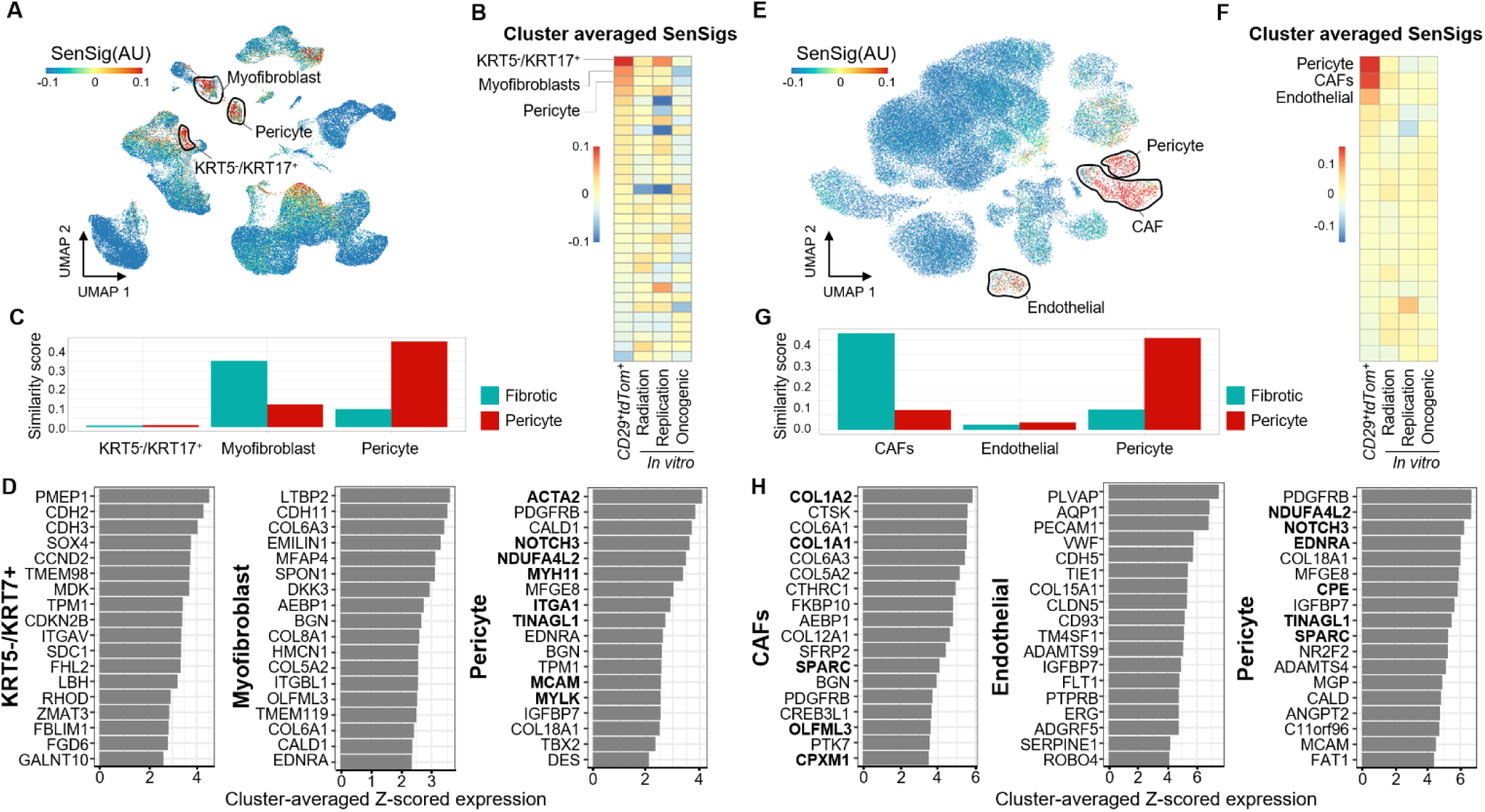
Transfer learning of senescence signature in alternate microenvironments. Putative senescent characteristics of two publicly available scRNAseq data sets from human samples of idiopathic pulmonary fibrosis (left), and basal cell carcinoma (right). (A and E) Feature plots of SenSig calculated using our *in vivo* p16+ reporter mouse as described in Figure 3E. (B and F) Cluster averages of SenSig using our senescence signature as well as three *in vitro* derived publicly available gene sets. Clustering included in the publicly available data were used and are described in detail in the source publications. (C and G) Similarity scores using our murine, VML derived scRNAseq clusters as references. Higher similarity score indicates similar gene expression patterns to murine scRNAseq stromal clusters. (D and H) Genes driving the largest increase in SenSig in the three human stromal clusters with highest average SenSig. Values shown are average z-scored expression across cells in the target cluster. Genes that were also in the top 25 genes driving senescent signature in the murine fibrotic or pericyte clusters are shown in bold.

We next compared the IPF clusters with our murine clusters and found that two of the higher SenSig clusters mapped to our fibrotic fibroblast and pericyte clusters in both data sets (Fig. 5C and Supplementary Fig. 13C). The KRT5-/KRT17+ cluster did not map to our murine data set effectively, indicating that the KRT5-/KRT17+ population did not have a corresponding cell type in our murine data set. Despite this, SenSig was able to identify this cell population as senescent. This indicates that a cell type specific compositional bias is not driving elevated SenSig scores but rather that a true conserved SnC phenotype is present and is sufficient to identify senescence in new cell types that were not a part of the bulk senescence sequencing.

Finally, we investigate the genes driving the elevated SenSig in the top 3 clusters from each data set (Fig. 5D and Supplementary Fig. 13D). The pericyte-like clusters shared SenSig driver genes with their respective murine cluster. However, the myofibroblast clusters and the two top-scoring SenSig clusters had distinct SenSig driver genes which did not overlap with murine clusters. The absence of overlap in SenSig driver genes between the IPF myofibroblast and murine fibrotic clusters suggests that although they share some similar transcriptional signatures, there may be distinct features of senescence between these populations. Further, the identification of the aberrant basaloid and KRT5-/KRT7+ clusters as senescent, neither of which mapped to any murine clusters, suggest that although our gene signature is derived from foreign body response to a synthetic implant, it can be successful in identifying SnC in broader contexts and cell types beyond a pericyte and fibroblast.

We then applied the same analysis technique to a publicly available data set of basal cell carcinoma (Yost et al., 2019). Using the authors cluster annotations, we identified three clusters with elevated SenSig: pericytes, cancer associated fibroblasts (CAFs), and endothelial cells (Fig. 5E and F). The CAFs and pericytes mapped to our fibrotic and pericyte murine clusters respectively and, unsurprisingly, the endothelial cell population was not similar to either (Fig. 5G). Finally, examination of the driver genes within each cluster identified a number of fibrotic genes driving SenSig in the CAFs and a number of genes associated with smooth muscle formation in the pericytes as well as *NOTCH3* (Fig. 5H). While the IPF derived myofibroblast population did not have any overlap in driver genes with our murine fibrotic population, the cancer derived CAFs shared a handful of common driver genes (*COL1A1, COL1A2, SPARC, OLFML3,* and *CPXM1*). The CAFs also had a higher similarity score to our fibrotic population than the myofibroblasts. Together, these findings suggest that our fibrotic population is more similar to CAFs than the myofibroblasts derived from the IPF environment. The endothelial cell population had a number of traditional endothelial cell markers as driver genes (*VWF*, *PECAM1*, *TIE1*) none of which overlapped with either the fibrotic or pericyte SenSig driver genes. The identification of a SnC phenotype distinct from the reference data suggests that the transfer learning technique is generalizable to both new microenvironments and at least some new cellular phenotypes.

## Senescent pericytes activate myeloid cells through IL34 signaling in wound healing

To determine intercellular signaling patterns between SnC and other cells in the microenvironment, we collected a scRNAseq data set in VML treated mice 1 week post-injury with enrichment of CD45^+^ cells to ∼50% of the total cell population (Fig. 6A). After computationally identifying immune cell populations as well as the CD29^+^CD31^-^CD45^-^ stromal populations we previously identified as senescent, we used singleCellNet to label stromal cells in the CD45^+^ enriched data set with our previous cluster labels (Fig. 6B). The resulting data set contains both immune clusters as well as stromal clusters corresponding to what we described above. We then used Domino (Cherry et al., 2021) to identify any potential signaling between the SnC clusters and immune clusters. Domino is an algorithm which attempts to reconstruct intercellular signaling patterns by first identifying activated transcription factors within each cell and then finding receptors with correlated expression to the activated transcription factors. Finally, if corresponding ligand expression is captured by scRNAseq, the cluster expression is used to identify inter-cluster signaling. The results are a global signaling network connecting transcription factors activated in the data set with their correlated receptors and, if present in the data set, ligands which could potentially activate the receptors. By selecting transcription factors enriched in a target cell population, it is possible to identify signaling directed at that population, for example targeting the myeloid or fibrotic clusters (Fig. 6C and Supplementary Fig. 14).

**Figure 6.**
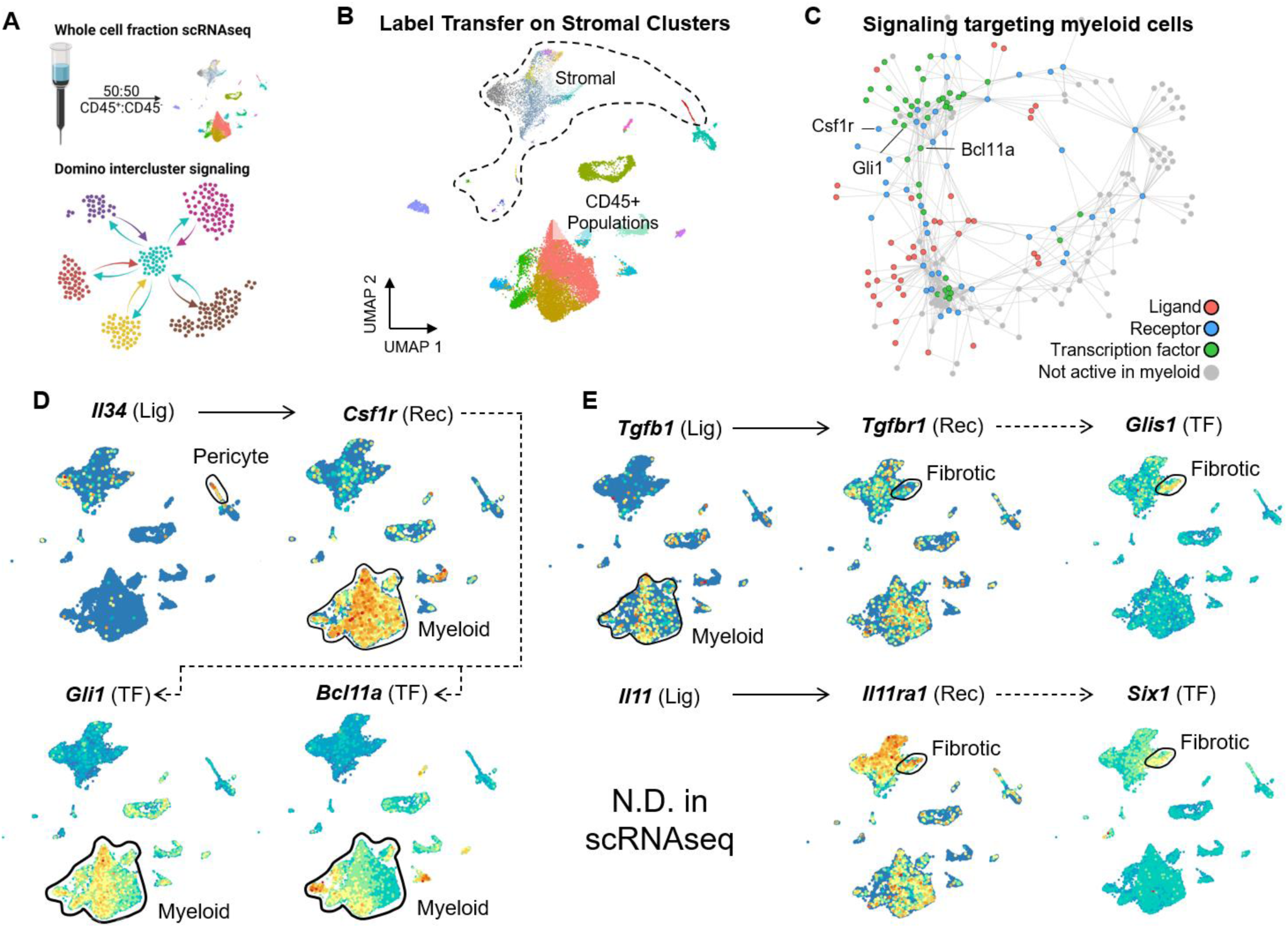
Intercellular signaling patterns involving SnC. (A) Summary of methodology to obtain intercluster signaling patterns. A whole cell fraction scRNAseq data set from mice treated with VML with saline, biomaterial implants, or naïve animals was obtained with enrichment of CD45^+^ cells using MACS beads. After mapping stromal clusters to the data set, Domino was used to calculate intercluster signaling patterns. (B) Visualization of stromal and non-stromal clusters within the whole cell fraction scRNAseq data set. Stromal clusters are colored as in the stromal scRNAseq data set shown in Figure 3. (C) A signaling pathway predicted to be active targeting myeloid cells by perictyes. *Il34*, expressed by pericytes in our data set, targets *Csf1r*, expressed by myeloid cells in our data set, and is predicted to activate *Gli1* and *Bcl11a* in the myeloid populations. (D and E) Feature plots of components of signaling pathways predicted to be active from pericytes targeting myeloid cells (D) and from myeloid cells targeting fibrotic fibroblasts (E). Solid lines represent ligands capable of activating receptors in the CellphoneDB2 database and dotted lines represent correlation between transcription factors and receptors.

Domino identified two major roles of senescent stromal cells in signaling to immune populations. First, it predicted IL34 from senescent pericytes would activate myeloid populations through Csf1r and ultimately activation of transcription factors Bcl11a and Gli1 (Fig. 6D). Importantly, *Il34* was one of the genes detected both in the bulk RNA sequencing senescent signature as well as upregulated in the pericyte scRNAseq cluster. It also predicted that Tgfb1 and IL11 would target the fibrotic fibroblast population of SnCs and ultimately lead to fibrosis through activation of transcription factors Glis1 and Six1 respectively, both of which have been implicated in fibrosis (Jetten, 2018; Liu et al., 2021) (Fig. 6E). The myeloid populations expressed the highest levels of *Tgfb,* although *Il11* was not detected in our single cell data set. Together, these findings suggest signaling from SnC pericytes influences differentiation of myeloid populations which, in turn, secrete molecules affecting the fibrosis by fibrotic populations (Fig. 7A).

**Figure 7.**
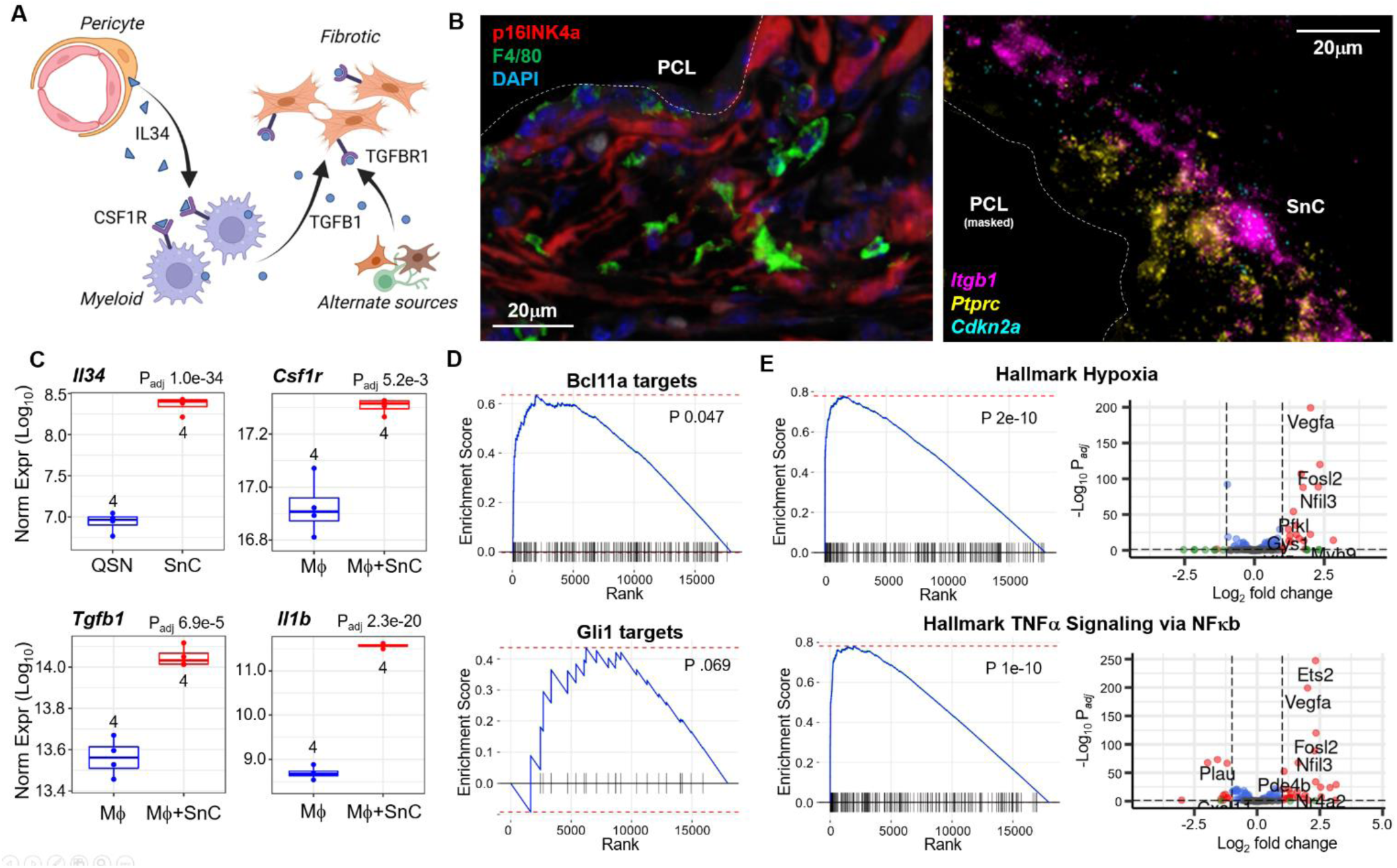
Validation of Domino predicted intercellular signaling patterns. (A) A graphical representation of signaling patterns involving the pericyte and fibrotic senescent populations predicted by Domino, an algorithm which predicts intercellular signaling patterns from scRNAseq data. In the PCL microenvironment, pericytes express IL34 binding to and activating CSF1R on myeloid cells. The myeloid cells further express TGFB1 which induces fibrosis in fibrotic SnCs through TGFBR1. (B) Immunofluorescent staining of P16 and macrophage marker F4/80 near the PCL implant (left). FISH staining for immune cell marker *Cd45* and stromal cell marker *Cd29* with *p16*near the PCL implant (right). Both images are representative of animals treated with PCL implants 6 weeks post-surgery. (C) Gene expression of select transcripts predicted to be involved in signaling between senescent pericytes and macrophages after coculture of senescent stromal cells with macrophages. Gene expression of *Il34* is compared between quiescent (QSN) and SnC stromal cells while other genes are shown in macrophages cultured alone or with SnC stromal cells in transwell plates. (D) Gene set enrichment for the Domino predicted Bcl11a and Gli1 target genes in macrophages cultured with SnC compared to cultured alone. Positive enrichment indicates overexpression of the module of genes predicted to be targeted by each transcription factor in macrophages cocultured with SnC. (E) Gene set enrichment for Hallmark Hypoxia and TNFa signaling gene sets in macrophages cultured with SnC compared to cultured alone (left). Positive enrichment indicates overexpression of the module of genes predicted to be targeted by each transcription factor in macrophages cocultured with SnC. Volcano plots showing fold-change and statistical significance of each gene in the gene sets are also shown (right).

To probe possible SnC communication, we first sought to determine whether senescent and myeloid cells were adjacent to each other in the fibrotic microenvironment *in vivo*. Both FISH staining for *Cd45* and *Cdkn2a* and immunofluorescent staining for P16 and F4/80, a macrophage marker, showed colocalization of immune/myeloid cells and SnC in mice treated with PCL implants (Fig. 7B). Immunofluorescent staining for P16, endothelial cell marker CD31, and macrophage marker CD68 in human fibrotic breast implant samples further demonstrated close proximity of senescent pericytes and fibrotic cells adjacent to macrophages, suggesting possible interactions and communication (Supplementary Fig. 15).

To further validate the computationally predicted interactions, we ran bulk RNA sequencing on senescent and quiescent stromal cells induced by irradiation and low serum media as well as macrophages alone or co-cultured with senescent stromal cells *in vitro*. We found elevated expression of *Il34* in senescent stromal cells compared to quiescent stromal cells as well as increased expression of *Csf1r*, the receptor for *Il34*, *Tgfb1*, and type 1 inflammatory cytokine *Il1b* in the macrophages co-cultured with SnCs (Fig. 7C). These results suggest that senescent stromal cells have elevated *Il34* and that upon co-culture with SnCs, macrophages increase expression of *Csf1r*, begin to secrete elevated levels of *Tgf1b*, and obtain an inflammatory phenotype as marked by increased expression of *Il1b*. Gene set enrichment analysis identified statistically significant over-representation of the Bcl11a and Gli1 targets predicted by Domino, suggesting that co-culture of macrophage with *Il34*-expressing SnC led to the changes in gene expression predicted by Domino (Fig. 7D). Finally, gene set enrichment analysis with the Hallmark gene sets identified overrepresentation of genes associated with hypoxia and TNFα signaling in macrophages co-cultured with SnC, confirming an inflammatory response in the co-cultured macrophages (Fig. 7E).

## Discussion

The field of senescence has grown significantly over recent years as the association of senescence and multiple age-related pathologies has been elucidated, however, the question of senescent cell identity, heterogeneity, and ultimately function remains. New transgenic models are enabling better visualization and identification of the rare cells expressing *p16*, however the phenotypic understanding of SnCs is limited due to the challenges in isolating these cells. Model systems that can enrich expressing cells enable the isolation of adequate cell numbers for in-depth phenotypic analysis. Fibrosis induced by the biomaterial foreign body response provides a reproducible system for SnC enrichment where p16 expressing cells can be readily identified, their phenotype characterized and specific senescent cell populations can be isolated. Bulk RNA sequencing of CD45^-^CD31^-^CD29^+^ SnCs captured 17,638 RNA transcripts after quality control filtering compared to the 6,751 transcripts expressed in more than 5% of cells in our single cell RNA sequencing data set. The relatively low transcript abundance in scRNAseq makes it difficult to identify SnC with transcripts like *Cdkn2a*, the transcript encoding p16, too low to detect by microfluidic-based scRNAseq techniques. The resulting differential expression profile represents the first direct, physiologically relevant, *in vivo* senescent gene expression signature.

Our SenSig transfer learning method provides a robust method to identify SnCs across tissues and species. Transfer learning in the context of scRNAseq refers to techniques which leverage existing or higher-fidelity data sets to identify similar cell populations across data sets (Stein-O’Brien et al., 2019). Importantly, these techniques can leverage gene expression patterns across hundreds or thousands of genes to overcome the inherently low transcript capture rates in scRNAseq. Here, we showed how the murine FBR-induced fibrosis senescence signature could be used to identify SnCs in the human foreign body response, idiopathic pulmonary fibrosis, and the tumor microenvironment. The SenSig approach identifies conserved properties of senescence while at the same time providing tissue-specific and pathology-specific context allowing for examination of specific genes driving the elevated signature within each cluster or even within each cell. Identification of senescent epithelial-derived basaloid cells in IPF from the stromal senescence signature highlights the power of the transfer learning to capture conserved elements of senescence within tissue specific cellular phenotypes not present in the reference data set. Moreover, publicly available senescence signatures derived from *in vitro* models did not lead to identification of senescent populations in any of the *in vivo* single cell data sets where there are known SnCs. This suggests that the *in vitro* models of senescence, ie, oxidative, oncogenic and proliferative, do not necessarily recapitulate physiologically relevant senescence in many pathologies. Other mechanisms of senescence, particularly immunologically-induced senescence (Faust et al., 2020), may provide new insights into physiologically relevant function that can be modelled *in vitro*.

One of the paradoxes of the senescence field is the beneficial contribution of SnCs during development and wound healing and the pathological influences in age-related diseases. Possible explanations suggest the existence of “good” versus “bad” SnCs with different phenotypes or kinetics of senescence that contribute to tissue development or pathology. Gene set enrichment analysis of the SnC expression profile identified two distinct pathways related to angiogenesis/vascularization and cartilage. The angiogenesis connection of SnCs was first identified by Demaria et al., in skin wounds where PDGF-BB could reconstitute SnC functions in tissue healing (Demaria et al., 2014). The presence of p16 expressing cells surrounding blood vessels visualized by immunofluorescence and the pericyte single cell cluster expressing the strongest SenSig similarity score further support the angiogenic contributions of SnCs and suggest that this population could be considered a beneficial senescent cell at least in the context of wound healing.

The annotation of the cluster with the second highest similarity with the SenSig was first classified as cartilage due to the large number of associated extracellular matrix molecules. FISH staining for the cluster associated gene *Fmod* confirmed the cells were in the regions of fibrosis and not contaminating tendon or ligament cells from the surrounding muscle. Fibrosis is however characterized by a thick, often aligned, extracellular matrix (ECM) similar to cartilage and tendon tissues. Many ECM proteins that are expressed by cells in the fibrotic region of the FBR are also found in cartilage including *Acan*, *Cilp*, *Chad*, and *Comp* (Vuga et al., 2013). While tissue-specific properties of fibrosis occur, it appears that fibrosis recapitulates aspects of native tendon and cartilage tissue structure to create a barrier tissue with dense, avascular, organized and frequently aligned extracellular matrix. Particularly with the correlation of senescence with dysregulated tissue function and pathological fibrosis, this unique senescent population provides a potentially new therapeutic target.

Interestingly, *CCN2* and *CCN5* were elevated in the senescent fibrotic fibroblast clusters as well as the in the sorted tdTom^+^ SnC. The CCN matricellular proteins have been implicated in induction of senescence in a variety of contexts (Joon-Il Jun & Lau, 2011; Joon-II Jun & Lau, 2017; Valentijn et al., 2021). They are also regulatory targets of the YAP/TAZ pathway, a mechanosensing pathway recently highlighted for its potential role in fibrosis (Dwivedi et al., 2020; Mascharak et al., 2022; Zhou et al., 2022). Activation of YAP was associated with increased fibrosis, increased expression of CSF1 leading to myeloid-derived inflammation, and increased expression of CCN family members. Taken together with our findings, we hypothesize that induction of YAP associated with tissue damage-associated mechanical changes in damaged tissues may induce senescence through secretion of matricellular CCN family members that subsequently activate myeloid cells, a process that is important for wound healing but that can also result in fibrosis.

While p16 is one of the primary markers used to identify SnCs and multiple transgenic mouse models have been developed to specifically monitor and clear *p16* expressing cells to extrapolate function, it has long been questioned as a definitive marker of senescence. For example, Dieckman et al, showed that p16 expression correlated with senescence but it was not a driver of senescence since inducing p16 expression did not lead to production of a SASP (Diekman et al., 2018). The senescent expression signature we present here was generated from p16 expressing cells but since it creates a signature of many genes that are increased and decreased it can be used to identify cells with similar expression changes regardless of p16 expression. This is demonstrated by the identification of senescent clusters in the single cell data sets where p16 expression is not captured and p21 expression, another marker of senescence, is expressed across multiple subsets. Cell intrinsic or extrinsic environmental factors may be responsible for the variable expression and need for p16 expression to arrest cell growth. Terminally differentiated cells such as those expressing many ECM molecules have limited replicative capacity and the dense fibrotic ECM in which the cells are embedded may reduce the need for cyclin dependent kinase inhibitors for growth arrest compared to cells such as pericytes active in angiogenesis. However, the fibrotic nature of the cartilage-like SnCs suggests these cells may be more associated with pathology rather than wound healing or potential beneficial effects of the angiogenic pericytes.

While fibroblasts were previously considered a supporting cell type, the complexity of fibroblast populations and their significant roles in multiple diseases and aging is now being elucidated. Single cell technologies opened the door to characterizing fibroblast heterogeneity and function while also providing markers for classification. Because p16 expression is not captured in scRNAseq data sets as mentioned previously, SnCs are not identified in the fibroblast or stromal single cell clusters in most data sets even though multiple studies identified fibroblasts as a cell type that becomes senescent (Chung et al., 2020; Demaria et al., 2014). Moreover, expression patterns and immunofluorescence of well accepted fibroblast markers, including Thy1.1, FAP, PDPN, and αSMA, do not distinguish senescent fibroblasts. The SenSig transfer learning approach contributes to senescence and fibroblast advances by identifying which subsets of fibroblasts exhibit high senescence signatures and distinguishing senescence heterogeneity with elucidation of distinct senescent fibroblast subsets.

Senescent cells may be a key player in immune-stromal interactions that are critical for tissue homeostasis and repair. Inflammatory fibroblasts and SnCs expressing a SASP produce immunologically relevant cytokines that support robust stromal communication with the immune system. We previously demonstrated a connection between SnCs and the immune environment in the foreign body response and osteoarthritis (Chung et al., 2020). In these models, there was a feedforward reinforcement between type 3 immune immunity characterized by Il17 production and senescence with an inflammatory-induced senescence signature identified *in vitro*. Of note, this is not immunosenescence but the impact of SnCs on the migration, activation and regulation of immune cell phenotype. T cells are intimately connected to the innate immune system and often drive myeloid skewing and macrophage phenotype and thus are likely connected, directly or indirectly with SnC behavior. Applying cell-cell communication analysis to a single cell data set where the SnCs are identified by transfer learning, we were able to predict the unique myeloid communication with pericyte and fibrotic SnCs that contributes to the balance of tissue vascularity and matrix production. The first prediction of senescent pericytes activating transcription factor programs in myeloid cells suggests coordinated stromal-immune activity in angiogenesis. Pericytes are well recognized as progenitor cells that can contribute to angiogenesis but myeloid cells also promote vascular development through production of VEGF as demonstrated in the FBR (Dondossola et al., 2016; Stockmann, Kirmse, Helfrich, Weidemann, & Takeda, 2011; Willenborg et al., 2012). A different set of myeloid cells expressed ligands that activated TFs in the fibrotic SnCs suggesting that the immune system contributes to pathological SnC activation and fibrosis.

Overall, we developed an *in vivo* senescence signature and SenSig transfer learning algorithm that identifies SnCs across species and multiple tissue pathologies. We found SnC populations associated with angiogenesis and fibrosis with unique myeloid communication profiles. Conserved senescence signatures and genes driving tissue-specific senescence discovered through transfer learning will enable detailed mapping of senescence and their impact on tissue physiology including stromal-immune regulation.

### Limitations of the Study

While our SenSig method appears to accurately identify SnC in a variety of contexts, the reference signature generated from the bulk RNA sequencing was created using implanted biomaterials in young mice. The methodology still appears to have successfully identified the disease-causing SnC in the IPF environment despite the IPF population not appearing transcriptionally similar to any cells in the biomaterial implant model, but nonetheless we cannot be certain that we are not missing SnC from differing contexts with differing expression signatures. Further, while we validate the SenSig method in our own murine model, we do not validate the findings after application to public data sets from differing pathologies. The SenSig score also does not identify true molecular senescence directly but only cells with similar expression patterns to the senescence.

The specific molecular characteristics of the SnC we identify are not fully explored. We explore intercellular signaling with Domino and validate both the SenSig and intercellular signaling with orthogonal biological experiments but do not investigate specific mechanisms of signaling. Extensive work remains to completely understand the dynamics of senescence formation in the fibrotic environment, how signaling is involved in skewing of cells to various SnC phenotypes, and how these processes may be accessible to pharmaceutical intervention.

## Supporting information

High Resolution Figures

## Acknowledgements

Department of Defense (W81XWH-17-1-0627 and W81XWH-14-1-0285), National Institutes of Health Pioneer Award DP1AR076959 (J.H.E.), Bloomberg∼Kimmel Institute (J.H.E., D.M.P.), Morton Goldberg Professorship (J.H.E.), Bristol Myers Squib (J.H.E., D.M.P.), National Science Foundation Graduate Research Fellowship Program DGE-1746891 (A.R. and A.N.P.), NCI U01CA253403 (E.J.F.), National Institutes of Health R01 AG057493 (J.M.v.D.). NIH T32 Training Grants 1T32AG058527-01 and 5T32CA153952-08 (J.I.A.). Biorender was used to create some of the figures presented in this manuscript. We would like to thank David R. Maestas Jr. for experimental insight and advice.

## Author contributions

C.C., J.I.A., and J.H.E. conceptualized and drafted figures and manuscript. C.C., K.K., J.H.E, and E.J.F. formulated, performed, and interpreted computational analysis of bulk and single cell RNA sequencing data sets. C.C., J.C.M., and K.K. performed Drop-seq single cell RNA sequencing. J.I.A. and L.D.H. performed the volumetric muscle loss surgeries. J.I.A., J.C.M., K.B.S., F.H., and D.M.P. performed and analyzed flow cytometry. H.H.N., A.N.P., and M.T.W. performed, imaged, and analyzed immunofluorescent staining and imaging including sectioning and sample processing. E.F.G.-G. performed and analyzed *in vitro* coculture experiments. A.R. performed, imaged, and analyzed fluorescence *in situ* hybridization staining and imaging. H.M. and J.M.v.D designed and generated the p16-EF/CreERT2 strain, and N.H., I.S., S.T., and D.J.B. contributed to various validation experiments for this model. J.I.A. and J.H.M. performed cryosectioning and imaging of native fluorescence of the fluorescent reporter. A.T. and J.I.A. performed fluorescence-activated cytometric sorting. C.J.L.S. performed and analyzed Ingenuity Pathway Analysis. All authors participated in construction of the manuscript and figures.

## Methods

### Animal Welfare Statement

All animal procedures were performed in adherence to approved JHU IACUC protocols.

### Volumetric muscle loss surgery and biomaterial implantation

All volumetric muscle loss surgeries were performed when animals were 10 weeks of age. Bilateral muscle resections of the quadriceps were performed as previously described. Defects were filled with 30 mg of (poly)caprolactone (particulate, Mn=50,000 g/mol, mean particle size < 600um, Polysciences), decellularized extracellular matrix (Matristem, Acell), or 50 uL of phosphate buffered saline as a control. All materials were UV sterilized prior to implantation and all animals given subcutaneous carprofen (Rimadyl Zoetis) at 5 mg/kg. Samples were harvested 1- or 6- weeks post-surgery. During harvest, the complete quadriceps together with any remaining implant material was harvested and processed for downstream experiments as described in the following sections.

### Clinical samples

Deidentified surgical discards from patients undergoing breast implant exchange or replacement surgeries were collected under Johns Hopkins University Institutional Review Board exemption IRB00088842. After collection, the samples were processed for downstream experiments as described in the following sections.

### Transgenic Mouse

The Ai14 reporter strain was purchased from Jackson Laboratories (strain #007908). The p16-CreER^T2^ strain was *de novo* created by inserting a gene cassette encoding CreER^T2^ at the exon 1-ATG start codon of the endogenous p16^Ink4a^ locus via homologous recombination in embryonic stem (ES) cells using standard procedures. The neomycin phosphotransferase (neo) gene sequence downstream of CreER^T2^ that was used to select targeted ES cell clones has been shown to serve as a transcriptional enhancer and was removed by FLP recombination to ensure exclusive control of CreER^T2^ expression by the endogenous p16^Ink4a^ gene promoter. Details on the generation and validation of the p16-CreER^T2^ strain will be provided elsewhere (Hamada et al., bioRxiv submission pending). Mice used for experiments were on a C57BL/6 x 129SvE mixed genetic background. Two weeks prior to collection of tissue from reporter mice, 5 intraperitoneal injections of tamoxifen solubilized in corn oil (14 mg/mL) and were administered daily at 70 mg/kg/day.

### MEF isolation and culture

Mouse embryonic fibroblasts were generated from embryonic day E13.5 embryos as described previously. *P16* knockout cells were described previously (Baker et al., 2008). Cells were expanded in DMEM (Gibco, #11960) supplemented with 10% heat-inactivated fetal bovine serum, L-glutamine, non-essential amino-acids, sodium pyruvate, gentamicin and β-mercaptoethanol. At cell passage 3, cells were harvested “d0 non-senescent” or subconfluent cell layers (60-80% confluence) were subjected to 10 Gy γ-radiation (^137^Caesium source) to induce senescence “d10 senescent”. Cells were harvested and subjected to downstream applications as described previously (Sturmlechner et al., 2021). Senescence-associated β-Galactosidase assay (Cell Signaling, #9860S) was performed and quantified as previously described. At least 100 cells were counted. For the EdU incorporation assay, cells were allowed to incorporate 1 μM EdU (5-ethynyl-2′-deoxyuridine, stock in DMSO) for 48 hours. EdU detection was performed according to the manufacturer’s protocol (Thermo Scientific, ClickiT Plus EdU Alexa Fluor 488 Imaging Kit, #C10637). At least 100 cells were quantified.

### Western Blot

Immunoblot analyses were performed as described previously (Kasper et al., 1999). Immunoblots were incubated with rabbit, anti-p16 (Santa Cruz, sc-1207; 1:1,000) antibody overnight and after washings, with HRP-conjugated goat, anti-rabbit antibody (Jackson Immunoresearch;1:10,000) for 3 hours. Ponceau S (PonS) (0.2% w/v in 5% glacial acetic acid, Sigma-Aldrich, #P3504) served as a loading control (Sturmlechner et al., 2021).

### Fibroblast isolation and culture

Dermal fibroblasts were isolated from 5-week-old C56Bl/6 female mice. Mice were euthanized, shaved, and a 1×2 cm section of dermal skin was removed and cut into small pieces before digestion. The dermal sections were digested for 1 hour at 37°C in 0.5 mg/ml Liberase TM (Roche) in serum free RPMI (Gibco + 2mM glutamine) shaking at ∼150 rpm. The digestion was neutralized with complete RPMI (10% fetal bovine serum [FBS, Gibco], 1% penicillin/streptomycin [Gibco], 1x sodium pyruvate [100x Gibco]), and the digested skin was pelleted via centrifugation for 5 minutes at 300g. The digested skin was resuspended in complete RPMI, divided into 3 T-175 flasks, and incubated at 37°C, 5% CO_2_ undisturbed for 4 days. On day 4, the cells were washed with PBS and the culture media was changed to complete MEM (10% fetal bovine serum [FBS, Gibco], 1% penicillin/streptomycin [Gibco]) to promote only fibroblast survival. Fibroblasts were then cultured in complete MEM until use.

### In vitro induction of senescence in stromal cells for coculture

Fibroblasts (passage 3-5), once 60-80% confluent, were placed in a minimal amount of media and an Xstrahl CIXD X-ray irradiator was used to deliver a dose of 10 Gy to the cells. The irradiated cells were washed with PBS and fresh complete MEM was added. The irradiated cells were incubated at 37°C 5% CO_2_ for 10 days to allow for the senescent phenotype to develop. Control quiescent fibroblasts were obtained by culturing fibroblasts in low-serum MEM (0.5% fetal bovine serum [FBS, Gibco], 1% penicillin/streptomycin [Gibco]). The senescent phenotype of the irradiation-induced senescent fibroblasts was verified with β-galactosidase staining (following the manufacturers protocol [Cell BioLabs Inc.]) and compared to the control quiescent fibroblasts.

### Macrophage isolation and culture

Bone marrow cells were isolated from 8-week-old C57Bl/6 female mice. Mice were euthanized and the bone marrow from both femurs was isolated. The red blood cells were lysed [BD Pharm Lyse], and the cells were passed through a 40μm strainer. The isolated cells were washed and plated on non-tissue culture treated plates in complete RPMI (10% fetal bovine serum [FBS, Gibco], 1% penicillin/streptomycin [Gibco]) supplemented with 20ng/mL of M-CSF [Miltenyi Biotech]). On day 3 and 5, 50% of the media was removed and replenished with fresh complete RPMI supplemented with 20ng/mL of M-CSF. On day 7, the adherent cells (differentiated macrophages) were detached using 0.25% trypsin (Gibco) and used for experiments.

### Fibroblast-macrophage co-culture

Macrophages (500,000 cells/well) were plated in 12-well plates and allowed to adhere before co-culture. Fibroblasts, either quiescent control or senescent, were seeded (100,000 cells/well) onto transwell membranes (Corning) and allowed to adhere before co-culture. Upon co-culture, macrophages were either cultured alone, with quiescent fibroblasts, or with senescent fibroblasts in complete RPMI for 6 hours (n=4 for each condition). After 6 hours, the macrophages and the fibroblasts (QSN or SnC), were individually lysed and the respective RNA was isolated using the RNeasy micro kit (Qiagen) for bulk RNA-sequencing.

### Isolation of RNA

After collection of tissue samples, samples were submerged in TRIzol reagent, and finely minced and grinded. For *in vitro* cultures, media was aspirated from tissue culture plates and TRIzol added according to the manufacturers recommended volume. The TRIzol was repeated pipetted across the dish. RNA isolation for both tissue and culture samples was then performed using Qiagen RNeasy Mini Kits following TRIzol extraction.

### Quantitative PCR

cDNA synthesis was performed using SuperScript VILO and quantitative PCR performed using TaqMan probes and reagents with the StepOne Plus Real-Time PCR System (Thermo Fisher Scientific) with 50ng of product per reaction. Analysis of data was performed with the delta-delta ct method using *B2m* and *Actb* as housekeeping genes. GraphPad Prism was used to perform statistical analysis using an ANOVA followed by multiple t-tests with Benjamini-Hochberg multiple test correction. For RNA isolation of MEF cultures, cDNA was generated with SuperScript III First-Strand Synthesis kit (Qiagen, #18080051) and SYBR Green Real-Time PCR Master Mix (Applied Biosystem, #4309155) was used to perform qPCR analyses. *Tbp* was used as housekeeping gene. Primers were:

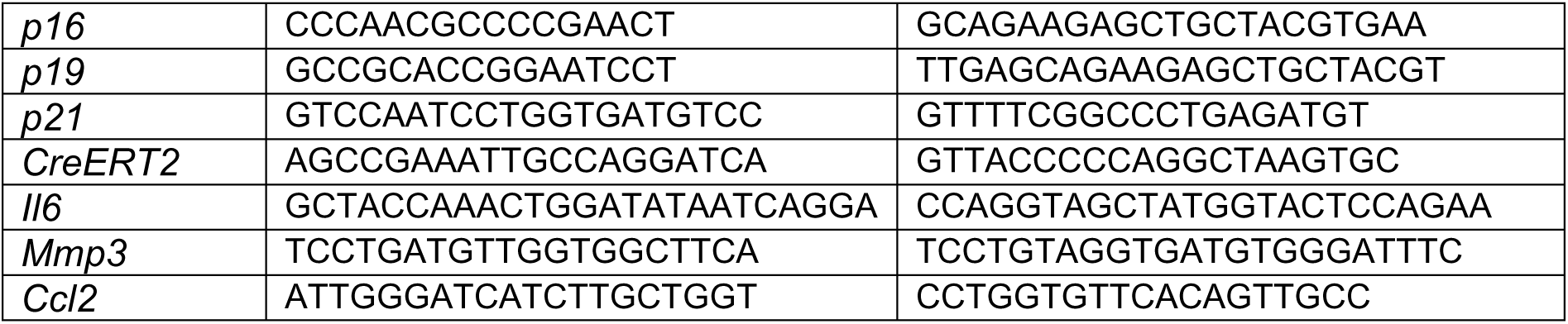

### Bulk RNA library preparation and sequencing

After isolation of RNA as described previously, cDNA synthesis and library preparation were performed using the TruSeq RNA Library Prep Kit V2. The Agilent BioAnalyzer system was used to perform quality control on samples before and during library preparation. RNA samples with RIN scores below 7 post-cDNA synthesis were removed from analysis. Libraries were pooled and sequenced with paired-end sequencing to a depth of 50bp for each end at a targeted depth of 30M unique reads per sample.

### Computational analysis of bulk RNA sequencing data

Reads were aligned with STAR (Dobin et al., 2013) against the GENCODE (Frankish et al., 2021) release M23 GRCm38.p6 genome and accompanying annotations. Differential expression was calculated using a negative binomial model using edgeR (Robinson, McCarthy, & Smyth, 2010). EnhancedVolcano (Blighe, Rana, & Lewis, 2020) was used to generate volcano plots. The differential expression results were evaluated for gene set enrichment using fgsea (Korotkevich et al., 2021) with a total of 1 billion permutations were used with gene set enrichment, leading to a minimum possible p-value of 1e-8 for each gene set. For the tdTom sorted bulk RNA sequencing samples both the hallmark and gene ontology gene sets were used. For the coculture of macrophages with senescent stromal cells the hallmark gene sets and the predicted transcriptional targets of transcription factors of interest from Domino were used. Finally, upstream regulators of differentially expressed genes were evaluated using Ingenuity Pathways Analysis with standard settings using the differential expression test results from edgeR.

### FFPE sectioning

After tissue collection, tissue was fixed in 10% NBF for 48 hours. Next, tissue was rinsed in water, then rinsed twice in 1X D-PBS, each time for 15 minutes. Tissue was dehydrated using 70% ethanol, 80% ethanol, 95% ethanol, and 100% ethanol at room temperature, changing the solution at hourly intervals. Tissue was cleared using xylene for 1 hour, followed by overnight incubation in melted paraffin wax in the oven at 55°C. The next morning, tissue was embedded in metal molds filled with paraffin. The molds were left on the cooling plate for a few hours until paraffin solidified. The formalin-fixed paraffin-embedded blocks were removed from the mold and stored at room temperature until sectioning. Human breast capsule and mouse quadricep tissue were sectioned at 5 µm and 7 µm, respectively, using a microtome.

### Hematoxylin and Eosin staining

Slides were deparaffinized in xylene and rehydrated using decreasing concentrations of ethanol and water. Slides were incubated in Harris Hematoxylin solution (Sigma-Aldrich, product number HHS32) for 3 minutes and then rinsed with deionized water. Slides were washed with tap water for 5 minutes and immersed in 0.33% hydrogen chloride – ethanol solution until the tissue background became clear. Subsequently, slides were rinsed in Scott’s tap water (Sigma-Aldrich, product number S5134) and immersed in Eosin Y solution (Sigma-Aldrich, product number HT110216) for 10 seconds. Lastly, slides were dehydrated in ethanol and xylene and finally mounted using Permount mounting medium (Fisher Scientific, product number SP15-100).

### Masson’s Trichrome staining

Slides were deparaffinized in xylene and rehydrated using decreasing concentrations of ethanol and water. Slides were stained in heated Bouin’s solution (Sigma-Aldrich, product number HT10132) at 56°C for 30 minutes and then cooled down for 3 minutes inside a chemical hood. Slides were washed with tap water for 3 minutes to remove the yellow color from the sections. Slides were stained in Weigert’s iron hematoxylin solution (Sigma-Aldrich, product number HT1079) for 5 minutes and washed in running tap water for 5 minutes. After that, slides were stained in Biebrich Scarlet-Acid Fucshin (Sigma-Aldrich, product number HT15-1) for 5 minutes and rinsed again in tap water until the samples had the desired color shade (light red). Subsequently, slides were stained in Phosphomolybdic Acid / Phosphotungstic Acid mix (Sigma-Aldrich, product number HT15-3 and HT15-2) for 7 minutes and in Aniline Blue Solution (Sigma-Aldrich, product number HT15) for 5 minutes. After that, slides were immersed twice in 1% acetic acid for a minute each time. Slides were rinsed in deionized water for 2 minutes until the water became clear and de-hydrated quickly in increasing concentrations of ethanol and xylene. Slides were mounted using Permount mounting medium (Fisher Scientific, product number SP15-100).

### Immunofluorescent staining

Slides were deparaffinized in xylene and rehydrated using decreasing concentrations of ethanol. Heat-induced epitope retrieval was performed in sodium citrate buffer, pH 6 for mouse tissue, and Tris-EDTA, pH 9 for human tissue, at 95°C for 15 minutes. Endogenous peroxidase activity was quenched using 3% H2O2 in PBS for 15 min. Slides were blocked in 10% BSA and 0.05% Tween 20 in PBS for 30 minutes before application of one of the following antibodies: mouse anti-human p16^INK4a^ (CINTEC-Roche, cat# 705-4793), rabbit anti-human alpha-smooth muscle actin or aSMA (Abcam, ab124964), mouse anti-human CD68 (Abcam, ab955), rabbit anti-human fibroblast activation protein or FAP (Abcam, ab218164), rabbit anti-human CD31 (Abcam, ab76533), rabbit anti-mouse p16^INK4a^ (Abcam, ab211542), rabbit anti-mouse CD31 (Abcam, ab182981), rabbit anti-mouse Fibromodulin or Fmod (Bioss, 12362R), rat anti-mouse F480 (eBioscience, 14-4801-82) for 30 minutes at room temperature. Subsequently, slides were incubated with horseradish peroxidase (HRP) polymer-conjugated secondary antibody for 20 min and reacted with one of the tyramide signal amplification reagents including Opal 520 (Akoya, FP1487001KT), Opal 570 (Akoya, FP1488001KT), Opal 650 (Akoya, FP1496001KT) for 10 min. Subsequently, antibodies were stripped by steaming in citrate buffer at 95°C for 15 minutes to allow the introduction of the next primary antibody with an Opal dye that is different from the first one. Cell nuclei were then counterstained with 4′,6-diamidino-2-phenylindole (DAPI) for 5 min before being mounted using DAKO mounting medium (Agilent, catalog no. S302380–2). Imaging of the histological samples was performed on the Microscope Axio Imager.A2 (Carl Zeiss Microscopy, LLC, model #: 490022-0009-000) and Zeiss ZEN 3.4 (blue edition) software.

| Target Antigen | Target Species | Host Species | Host Isotype | Clone | Manufacturer | Catalog # |
| --- | --- | --- | --- | --- | --- | --- |
| p16<sup>INK4a</sup> | Human | Mouse | IgG<sub>2a</sub> | E6H4 | CINTEC-Roche | 705-4793 |
| aSMA | Human | Rabbit | IgG | EPR5368 | Abcam | 124964 |
| CD68 | Human | Mouse | IgG1 | KP1 | Abcam | 955 |
| FAP | Human | Rabbit | IgG | polyclonal | Abcam | 218164 |
| CD31 | Human | Rabbit | IgG | EPR3094 | Abcam | 76533 |
| p16<sup>INK4a</sup> | Mouse | Rabbit | IgG | EPR20418 | Abcam | 211542 |
| CD31 | Mouse | Rabbit | IgG | EPR17259 | Abcam | 182981 |
| Fmod | Mouse | Rabbit | IgG | polyclonal | Bioss | 12362R |
| F480 | Mouse | Rat | IgG<sub>2a</sub> | BM8 | eBioscience | 14-4801-82 |

### RNAscope HiPlex Staining

RNA fluorescence in situ (FISH) staining was performed to visualize spatial distribution of stromal and immune populations in injured murine quadricep muscles with PCL implants. RNA probe targets were selected based on differentially expressed genes identifying distinct fibroblast clusters from single-cell RNAseq analysis of sorted stromal cells from the muscle injury model. All samples were processed using ACD RNAscope HiPlex Assay protocol (Wang et al., 2012) and reagents for fixed frozen samples (ACD 324100-UM, Cat 324100, 324140). In short, the injured muscles (and surrounding material implants) were harvested, fixed (10% formalin) and frozen in optimal cutting temperature (OCT) embedding media following a sucrose gradient. Tissue cryosections (14um, transverse orientation) were baked, post-fixed and dehydrated prior to antigen retrieval and protease treatment. Custom HiPlex RNA probes (12 probes per assay, T1-T12) were hybridized (2hr, 40°C) and HiPlex amplified, then probes T1-T3 were fluorophore conjugated (alternate display module: AF488, Atto550, Atto675N). Cryosections were counterstained with DAPI, mounted, and imaged with a Zeiss Axio Imager A2 (20x objective, Zeiss ZEN 3.4 software). For subsequent rounds of probe staining (T4-T6, T7-T9, and T10-12), coverslips were removed (4x SSC buffer), previous fluorophores cleaved (10% Cleaving Stock for 15min, PSBT-0.5%Tween washes), new tagged-fluorophores conjugated, and cryosections imaged. RNAscope HiPlex Image Registration Software (ACD Cat. 300065, v1) was used to register the multiple rounds of imaging based on DAPI. Image visualization was performed using Zeiss ZEN 3.4 (blue edition), Aperio ImageScope (Leica, 12.4.3.5008) and Fiji software (Schindelin et al., 2012). Autofluorescence from remaining PCL particles was manually marked and masked. Overall image brightness was increased 20-40% to improve visualization of stained RNA probes.

### Imaging of native tdTom expression

Muscles (and surrounding material implants) were harvested, fixed (10% paraformaldehyde) and frozen in optimal cutting temperature (OCT) embedding media following a sucrose gradient. Tissues were sectioned at 14 μm. Cryosections were counterstained with DAPI, mounted, and imaged with a Zeiss Axio Imager A2 (20x objective, Zeiss ZEN 3.4 software).

### Isolation of single cell suspensions

All single cell suspensions isolated from tissue samples for flow cytometry and scRNAseq were processed identically. Tissue samples were finely diced and digested in RPMI 1640 media (Gibco) with 1.6 Wunsh U/mL Liberase TL (Roche Diagnostics) and 0.2 mg/mL deoxyribonuclease I (Roche Diagnostics) for 45 minutes at 37°C. Digested tissues were ground through a 100 μm then 70 μm strainer (Thermo Fisher Scientific) with excess RPMI and then washed twice with 1x PBS.

### Flow cytometry and fluorescence-activated cell sorting

After isolation of single cell suspensions, cells were washed once with DPBS and stained with LIVE/DEAD™ Fixable Aqua Dead Cell viability dye (Thermo Fisher Scientific) or eBioscience Fixable Viability Dye eFluor780 (Thermo Fisher Scientific) for 30 minutes on ice. Following incubation, the cells were washed twice with FACS buffer 1x DPBS + 1% w/v BSA (Millipore Sigma) + 1mM EDTA and stained for surface antigens with fluorophore-conjugated antibodies. After incubating in the dark for 45 minutes on ice, the cells were washed as before with FACS buffer. Samples and controls were then resuspended in FACS buffer. Data was acquired on a 4 laser Attune Nxt Flow Cytometer (Thermo Fisher Scientific) and analyzed with FlowJo software (Tree Star). Compensation beads (BD Biosciences) were used at the time of data acquisition, but compensation was manually adjusted during analysis to account for the autofluorescent nature of muscle stromal cells. GraphPad Prism was used to perform statistical analysis using an ANOVA followed by multiple t-tests with Benjamini-Hochberg multiple test correction.

For sorting experiments, tissues were passed through 100 μm then 70 μm cell strainers (Thermo Scientific) with RPMI media then washed with FACS Buffer. Cells were then stained with LIVE/DEAD Fixable Aqua Dead Cell viability dye and the appropriate fluorophore-conjugated antibodies as above. After the final wash, cells were resuspended in FACS buffer and sorted using a BD FACSAria Fusion Sorter (BD Biosciences). Cells were sorted directly into either RLT Plus Buffer (Qiagen) or 1x DPBS containing 0.01% BSA for RNA extraction or scRNAseq sequencing, respectfully.

| **Antibody** | **Manufacturer** | **Catalog #** | **Clone** |
| --- | --- | --- | --- |
| Brilliant Violet 421 anti-mouse CD31 | BioLegend | 102424 | 390 |
| Brilliant Violet 605 anti-mouse CD45 | BioLegend | 103140 | 30-F11 |
| Brilliant Violet 711 anti-mouse CD11b | BioLegend | 101241 | M1/70 |
| Alexa Fluor 488 anti-mouse CD3 | BioLegend | 100210 | 17A2 |
| PE/Cyanine7 anti-mouse F4/80 | BioLegend | 123113 | BM8 |
| APC anti-mouse CD11c | BioLegend | 117310 | N418 |
| Alexa Fluor 700 anti-mouse/rat CD29 | BioLegend | 102218 | HMβ1-1 |
| Alexa Fluor 700 anti-mouse CD90.2 | BioLegend | 105320 | 30-H12 |
| APC-Cyanine 7 anti-mouse/rat CD29 | BioLegend | 102226 | HMβ1-1 |
| TruStain FcX (anti-mouse CD16/32) | BioLegend | 101320 | 93 |

### Magnetic-activated cell sorting

For scRNAseq samples where CD45^+^ cells were enriched to 50% of the total cell population, the cell suspension was run through a MACS column using the dead cell removal bead kit (Miltenyi Biotec) following manufacturer’s protocol. CD45 MicroBeads (Miltenyi Biotec) were then used to select the CD45 positive cells. These were then mixed 1:1 with the CD45 negative flow through for the final CD45 enriched mixture.

### Drop-seq single cell RNA sequencing

Single cell RNA sequencing data sets were collected using the Drop-seq (Macosko et al., 2015) pipeline. Cells were isolated and sorted as described in the flow cytometry methods from animals treated with decellularized extracellular matrix or PCL implants, control wounds, or naïve animals 1- or 6-weeks after surgery. Each sample was composed of three biological replicates pool after sorting to obtain sufficient cell numbers. The single cell suspension was manually counted and run through a microfluidic device which encapsulates the cells in aqueous droplets with barcoded beads and lysis buffer. After incubation, the emulsion is broken, beads collected, and cDNA synthesis performed with Maxima Reverse Transcriptase. After quality control and quantification with the Agilent BioAnalyzer high sensitivity DNA kit, whole transcriptome amplification was performed with the Kapa HiFi PCR kit. Finally, library preparation was performed with the Nextera XT kit. A detailed protocol is available from the McCarroll Lab website mccarrolllab.org/dropseq. Finally, libraries were pooled and sequenced with an Illumina NovaSeq to a target depth of 100,000 unique reads per cell.

### scRNAseq computational processing

Barcode identification, alignment, and counting were performed using the McCarroll lab’s DropSeq toolkit following the protocol and guidelines outlined in the Drop-Seq Cookbook. Read 1 was used to identify cell and molecular barcodes by read. Read 2, containing the transcript itself, was aligned to STAR using the GENCODE release M23 GRCm38.p6 genome and annotations. Unique molecular barcodes (UMI) were counted by cell barcode and cells separated from background using elbow plots of read count by barcode.

The Seurat (Hao et al., 2021) package was used for normalization, scaling, principal component analysis (PCA), clustering, and differential expression. Raw counts were first normalized to the total number of counts per cell barcode and then log scaled. Percentage of counts from mitochondrial genes and cell cycle scores were calculated with Seurat’s CellCycleScoring. These parameters were then regressed from the normalized counts and the resulting residuals z-scored, providing scaled inputs for PCA. 30 principal components were selected for downstream analysis using an elbow plot. The principal components were corrected for batch effect using Harmony (Korsunsky et al., 2019) with default settings. A shared-nearest neighbor graph and Louvain clustering was then performed on the corrected principal components. Cluster number selection was performed by identifying the local maximum silhouette score within a biologically significant range of cluster numbers. Dimensional reduction for visualization was performed using the PHATE (Moon et al., 2019) python package and the uwot implementation of UMAP (McInnes, Healy, & Melville, 2018) in R. Mann-Whitney U-test was used for differential expression in the single cell data set. Clusters were compared to all other cells in the data set to determine characteristic genes for cluster phenotypes.

Differentiation trajectories were determined using a combination of RNA velocity (La Manno et al., 2018), pseudotime (Street et al., 2018), and CytoTRACE (Gulati et al., 2020). Velocyto was used to calculate RNA velocities as well as visualize velocities on two-dimensional cell embeddings. CytoTRACE scores were calculated using default parameters. Slingshot was used to calculate differentiation trajectories using initial and terminal clusters as identified by RNA velocity and CytoTRACE (Gulati et al., 2020) scores.

Comparison between data sets was performed using singleCellNet (Tan & Cahan, 2019). singleCellNet generates similarity scores between two data sets using all clusters present in the reference data set. For cases where discrete groups were required for analyses, each cell in the target data set was scored according to the highest score from the reference data set. For cases of comparison across species, biomaRt (Smedley et al., 2009) with Ensembl annotations were used to convert murine genes to corresponding human genes or visa-versa.

Intercellular signaling analysis was performed with Domino (Cherry et al., 2021). First, transcription factor activation scores are estimated using SCENIC (Aibar et al., 2017). The resulting transcription factor activation scores are correlated with receptor expression and thresholded, identifying a number of receptors correlated with activation of target transcription factors across the entire data set. Finally, candidate ligands for target receptors are pulled from the CellphoneDB2 (Efremova, Vento-Tormo, Teichmann, & Vento-Tormo, 2020) database, generating a signaling network connecting ligands with candidate receptors and their putative transcriptional targets. The transcription factors within the network are evaluated for enrichment in target clusters with Mann-Whitney U test to generate subnetworks targeting specific clusters.

### Transfer learning for identification of SnCs from single cell data sets

Senescent signatures were calculated using the set of differentially expressed genes from the bulk RNA sequencing comparing p16^+^ and p16^-^ stromal cells. The set of differentially expressed genes (FDR < .01) and their accompanying direction of fold change (increased or decreased in SnCs) were used. To generate a score for each cell, the z-scored expression values were filtered for the genes of interest, converted to indicate concordance or discordance with the direction of change in the bulk sequencing data set, and averaged. A positive resulting score indicates a similar expression pattern to the bulk signature compared to the other cells in the single cell data set.

### Analysis of public single cell RNA sequencing data sets

All public single cell RNA sequencing data sets presented in this analysis provided cluster labels which were used as is with the exception of the basal cell carcinoma data set where were changed the myofibroblast label to endothelial cell based on expression of endothelial cell marker *VWF*. The data sets were also prefiltered and no further filtering was needed after visualization of total UMI count and feature count by cell. UMAP reductions were determined as described above with the exception of the removal of Harmony for batch effect correction.

---

**Supplementary Figure 1.**
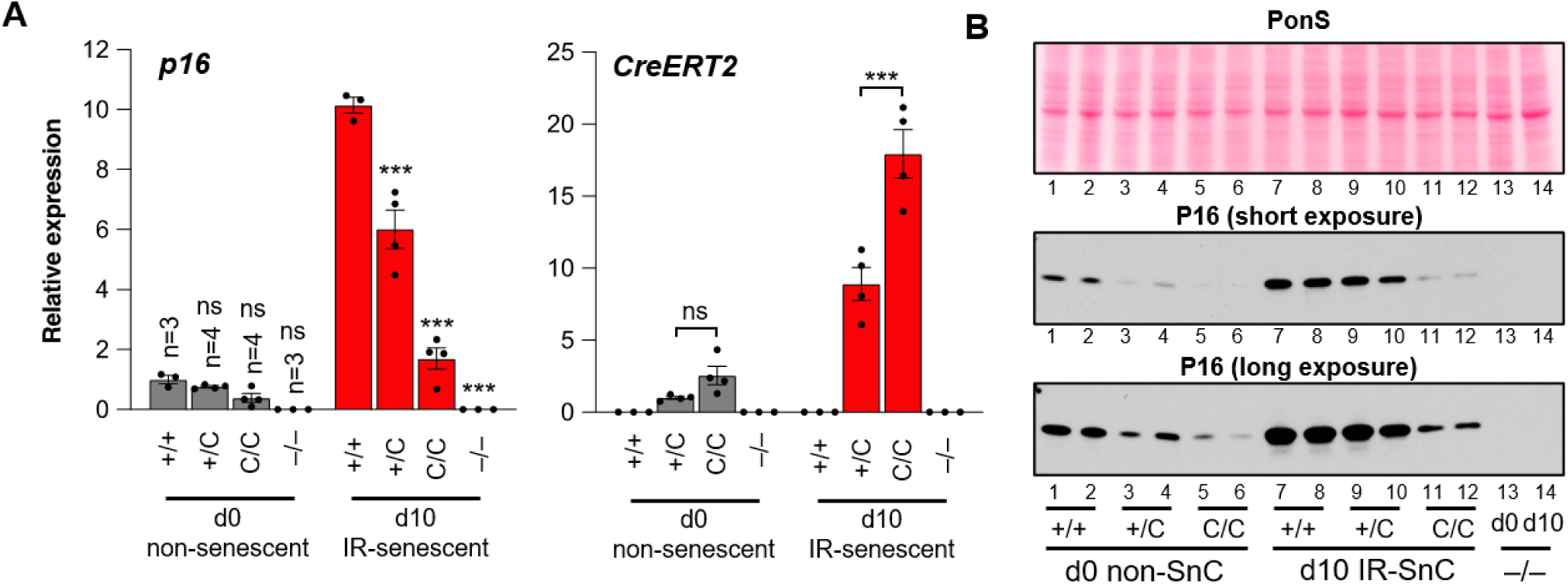
Validation of the *p16-EF/CreER^T2^;Ai14* reporter model. (A) Expression of *p16* and *CreERT2* as quantified by qPCR in mouse embryonic fibroblasts derived from animals with indicated genotypes for the *p16* allele. Alleles are either wild type p16 (+), CreER^T2^ knocked into the *p16*locus (C), or p16 knockout (–). Animals heterozygous for the wild-type and Cre construct are indicated as +/C. Cells were collected before irradiation induction of senescence (d0) or 10 days post irradiation. ns, not significant. (B) Western blot for p16 in mouse embryonic fibroblasts derived from animals with indicated genotypes for the p16 allele. Alleles are either wild type *p16* (+),CreER^T2^ knocked into the *p16* locus (C), or *p16* knockout (–). Cells were collected before irradiation induction of senescence (d0) or 10 days post irradiation. Ponceau S (PonS) served as loading control.

**Supplementary Figure 2.**
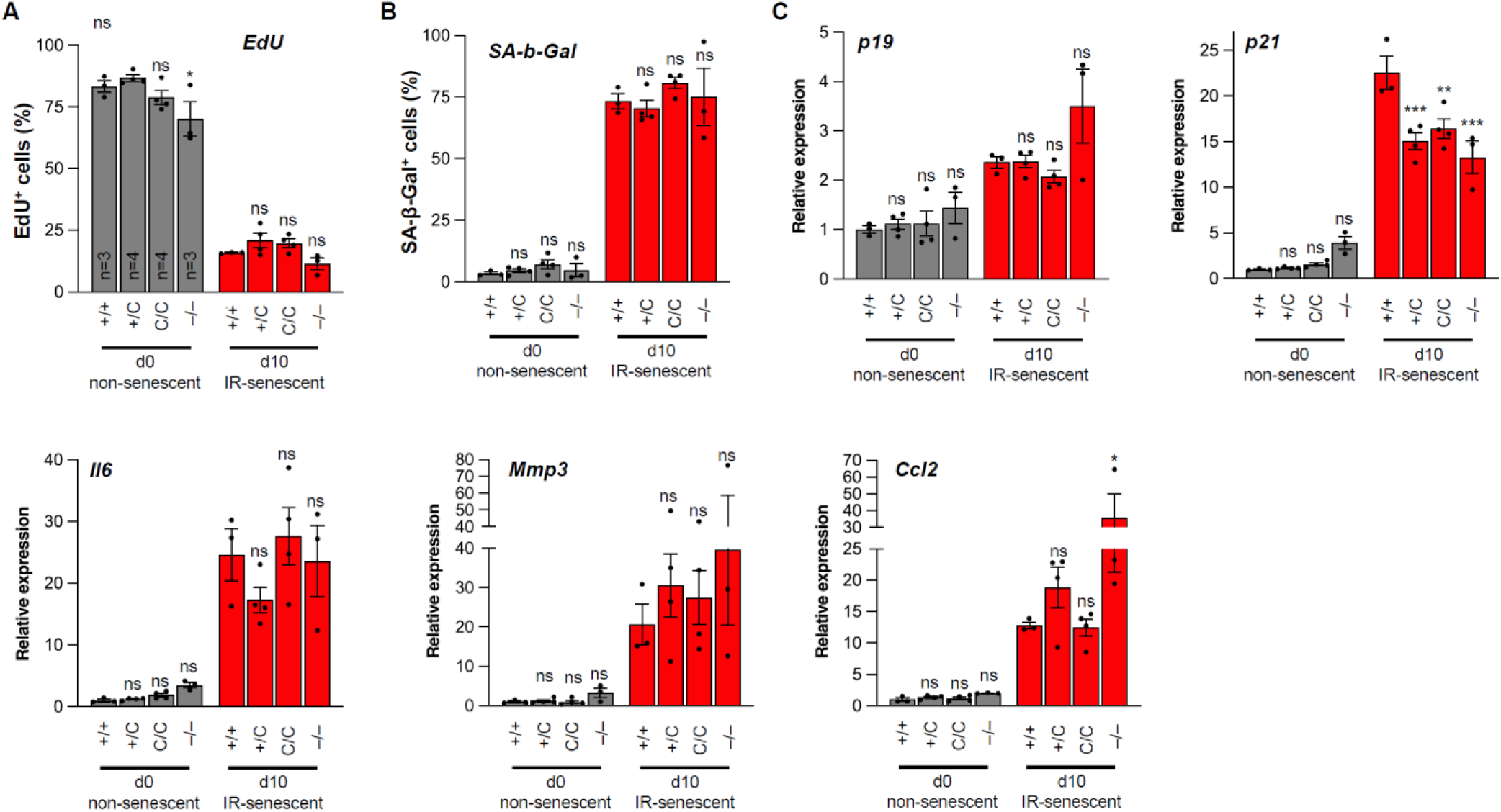
Phenotypic evaluation of irradiated mouse embryonic fibroblasts from reporter animals. (A) Quantification of cellular replication by EdU incorporation in mouse embryonic fibroblasts derived from animals with indicated genotypes for the *p16* allele. Alleles are either wild type *p16* (+),CreER^T2^ knocked into the *p16*locus (C), or *p16* knockout (–). Cells were allowed to incorporate EdU for 48 hours. Cells were collected before irradiation induction of senescence (d0) or 10 days post irradiation. (B) Same as in (A) but employing a senescence-associated β-galactosidase assay. (C) Same as in (A) but measuring mRNA expression of senescence associated genes as quantified by qPCR. ns, not significant. Statistical comparisons were made to the +/+ control group per timepoint.

**Supplementary Figure 3.**
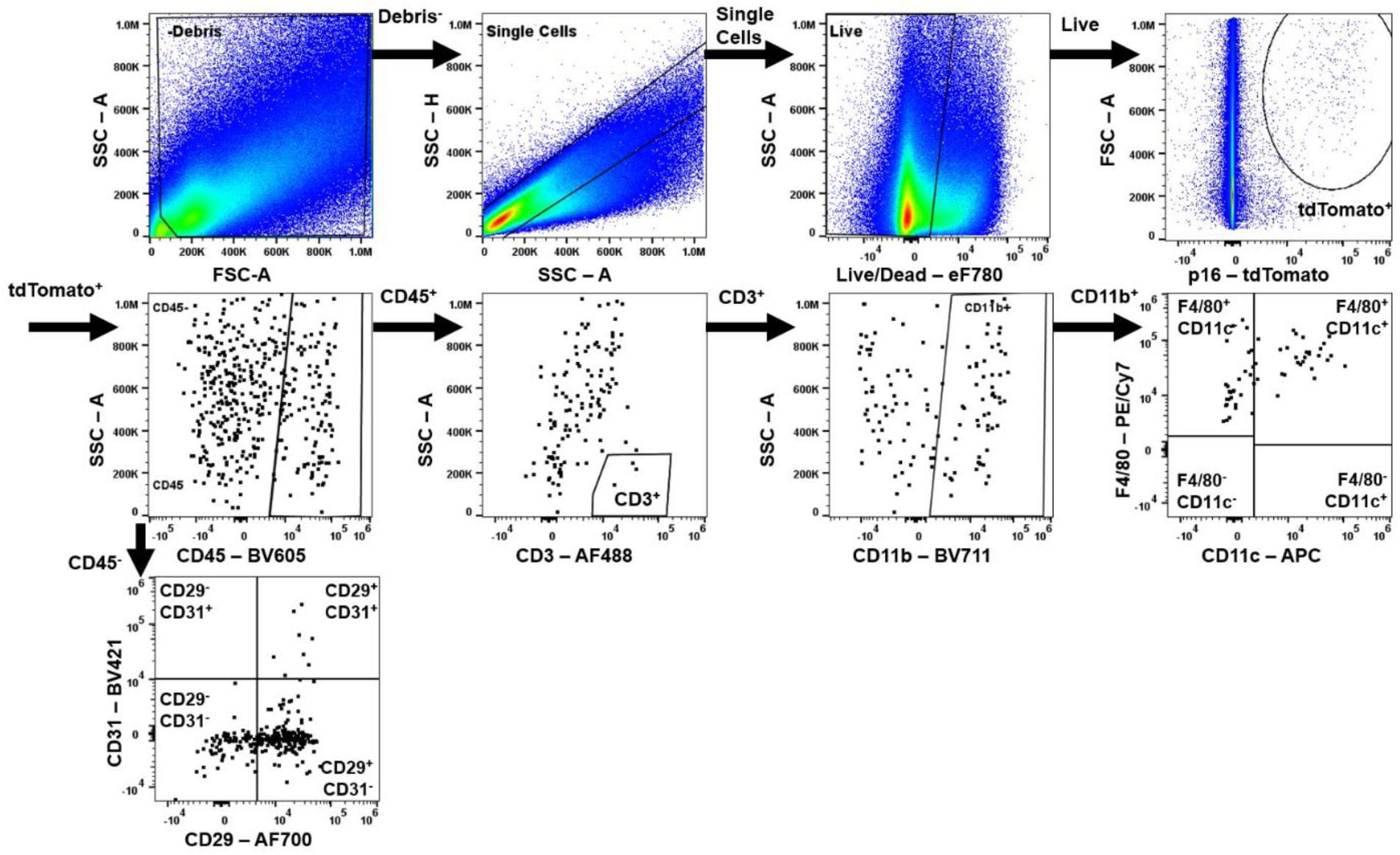
Flow cytometry gating for p16^+^ subsets. Example gating for VML injuries treated with PCL or Saline with or without tamoxifen (TAM) to induce tdTom expression . tdTom^+^ cells were selected after gating live cells. Endothelial (CD45^-^CD31^+^) and stromal cells (CD45^-^CD29^+^CD31^-^) were gated from the CD45^-^ population. Of the CD45^+^ populations, CD3^+^SSC^lo^ cells were gated as T-cells. Myeloid populations were then selected as CD45^+^CD3^-^CD11b^+^ and delineated into CD11c^+^ and CD11c^-^ macrophages with F4/80^+^ (CD45^+^CD3^-^CD11b^+^F4/80^+^).

**Supplementary Figure 4.**
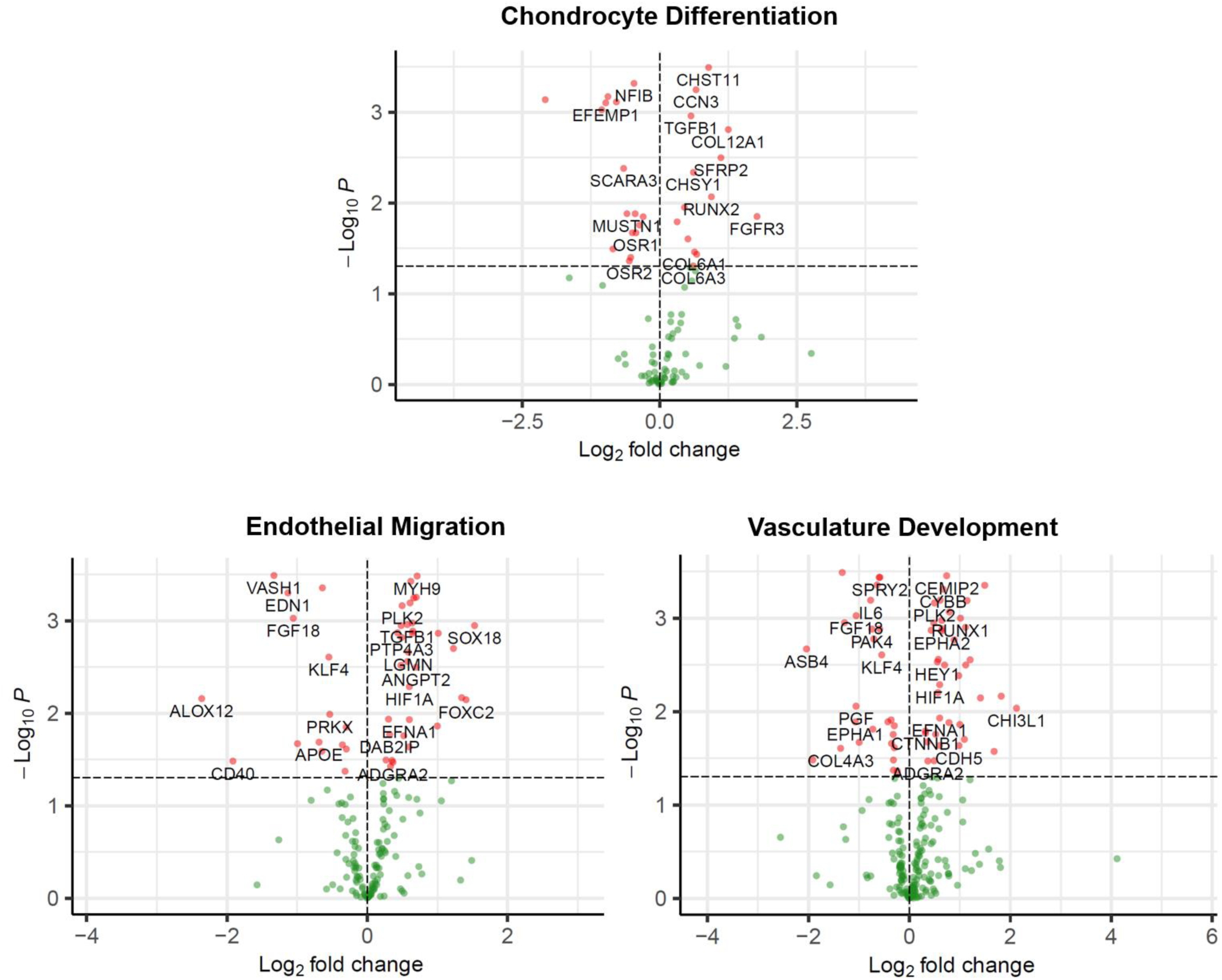
Gene expression patterns in sorted senescent stromal cells. Volcano plots of differentially expressed genes for gene ontology gene sets highlighted in Figure 2D. All genes in each gene set are shown.

**Supplementary Figure 5.**
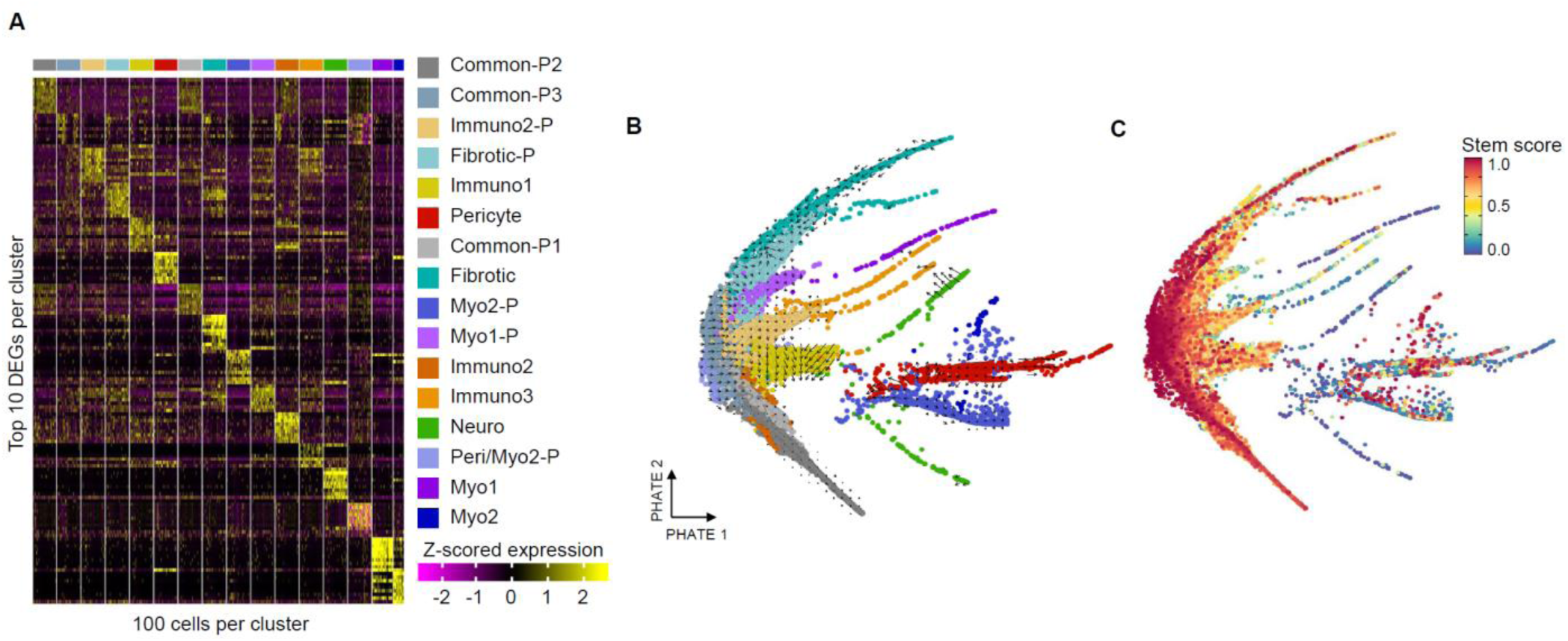
Characteristics of scRNAseq clusters and lineages. (A) Differentially expressed genes by cluster. Up to 10 positively differentially expressed genes were selected based on fold-change (rows) and 100 cells were sampled from each cluster. Z-scored gene expression values are shown for each cell by gene. (B) RNA velocity superimposed in PHATE embeddings. Cells are colored by cluster and positioned according to their PHATE embedding. RNA velocity is calculated by velocyto indicating the predicted movement of transcriptional characteristics within each cell. (C) CYTOtrace scores for stemness. Cells are shown on their PHATE embedding and colored by CYTOtrace score which predicts stemness of cells. High score indicates stem-like character.

**Supplementary Figure 6.**
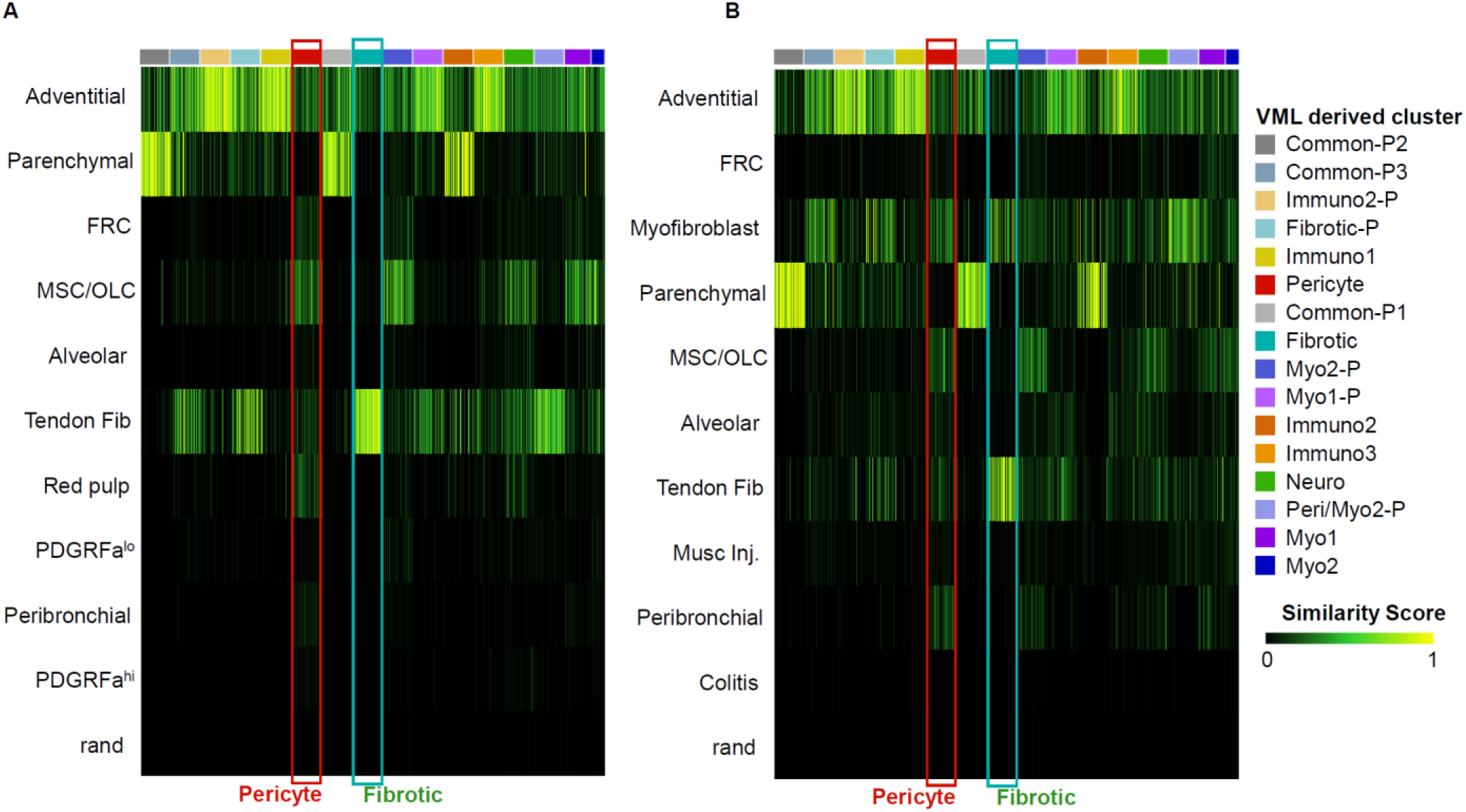
Comparison to public fibroblast single cell atlases. (A) Comparison of the VML stromal data set to a steady state fibroblast atlas described by Buechler et al. singleCellNet was used to map the clusters provided from the reference atlas to our stromal single cell data set. Heatmap rows indicate clusters from the reference atlas and columns indicate VML derived stromal clusters. Putative senescent populations from the PCL implant data set are indicated by red and teal boxes. Higher similarity scores indicate similar transcriptional characteristics between the reference clusters (rows) and the cells in the VML data set (column). (B) Comparison of the VML stromal data set to a perturbed state fibroblast atlas from Buechler et al. Processing and transfer scoring is identical to (A).

**Supplementary Figure 7.**
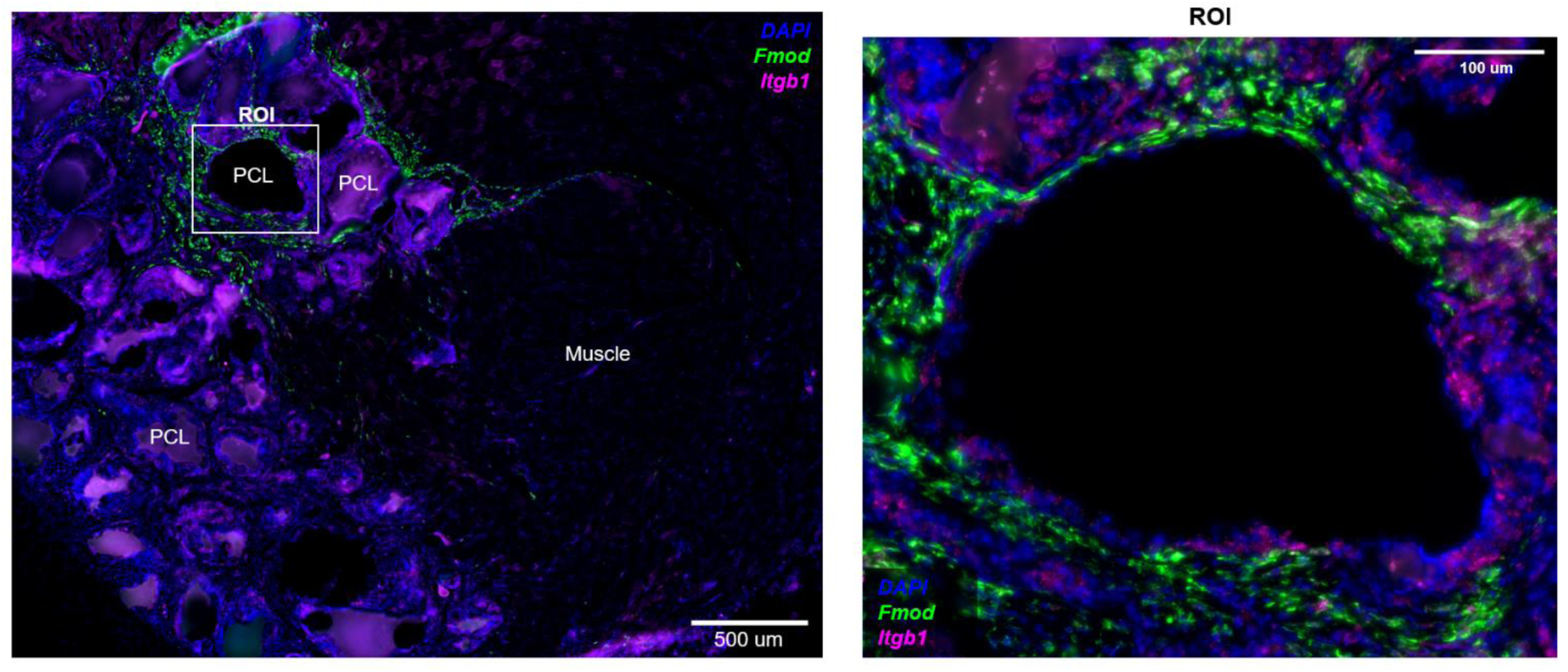
Fluorescent *in situ* hybridization for fibrotic marker *Fmod*. Fluorescent *in situ* hybridization for fibrotic marker *Fmod* in an animal treated with PCL implant 6 weeks post-surgery, scale bar = 500 μm. The general area of the uninjured muscle and the area of the PCL implants are labeled. A higher magnification image of the region positive for *Fmod* is shown, scale bar = 100 μm.

**Supplementary Figure 8.**
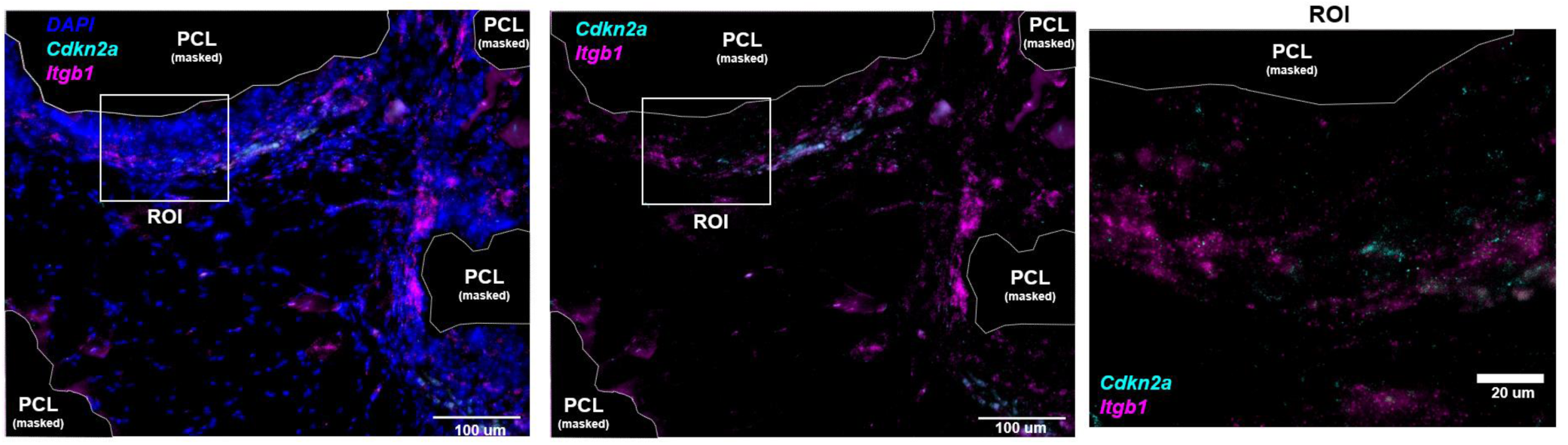
Fluorescent *in situ* hybridization for *Cdkn2a*. Fluorescent *in situ* hybridization for fibrotic marker *Cdkn2a* in an animal treated with PCL implant 6 weeks post-surgery, scale bar = 100 μm. A higher magnification image of the region positive for *Cdkn2a* is shown, scale bar = 20 μm.

**Supplementary Figure 9.**
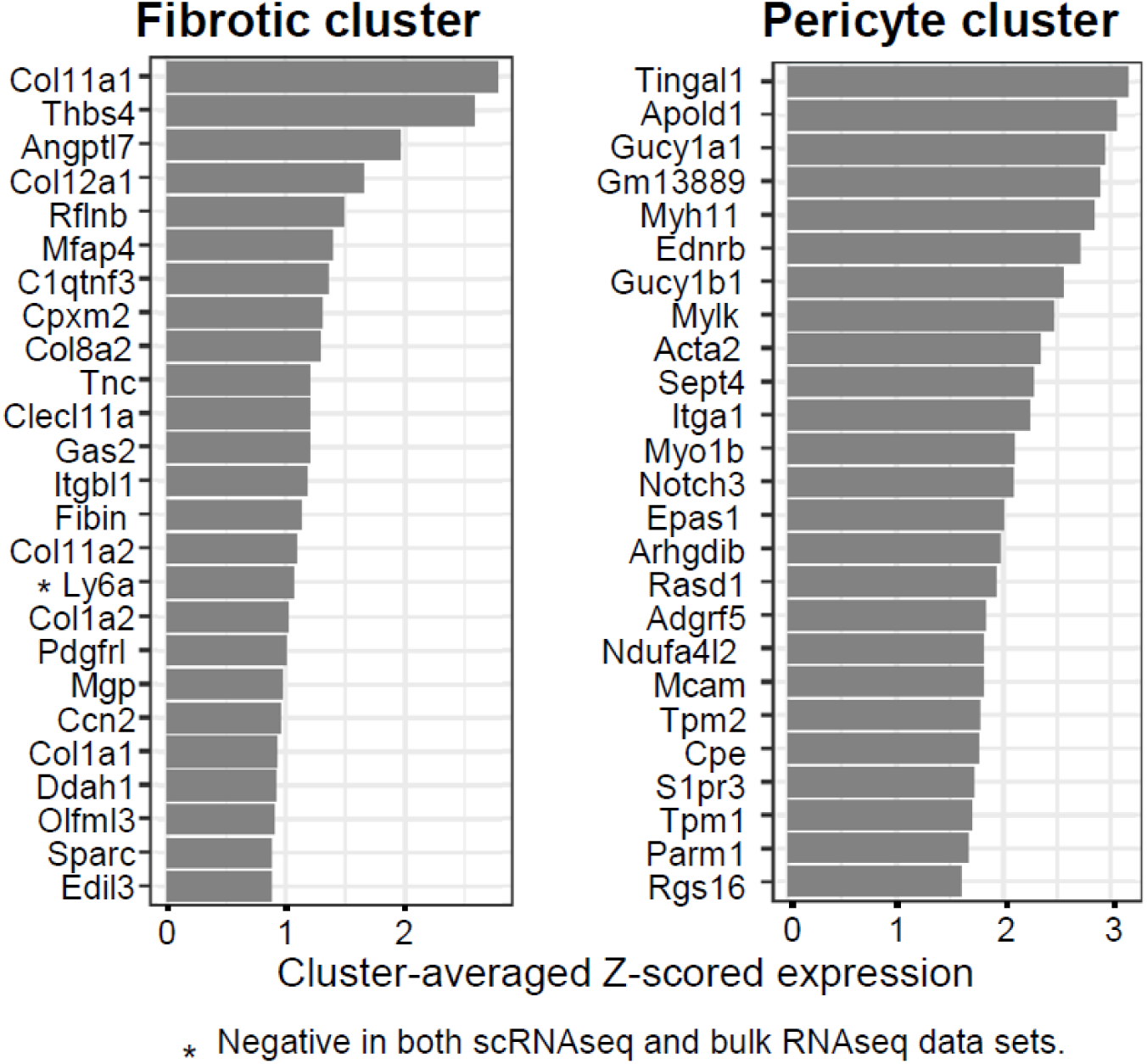
Top genes driving SenSig in the putative SnC clusters. The top 25 genes driving high SenSig in the fibrotic and pericyte clusters are shown. Scores shown are cluster-averaged z-scored gene expression. Only *Ly6a* was found to be negative in both the bulk and single cell data sets, all other genes were positive in both.

**Supplementary Figure 10.**
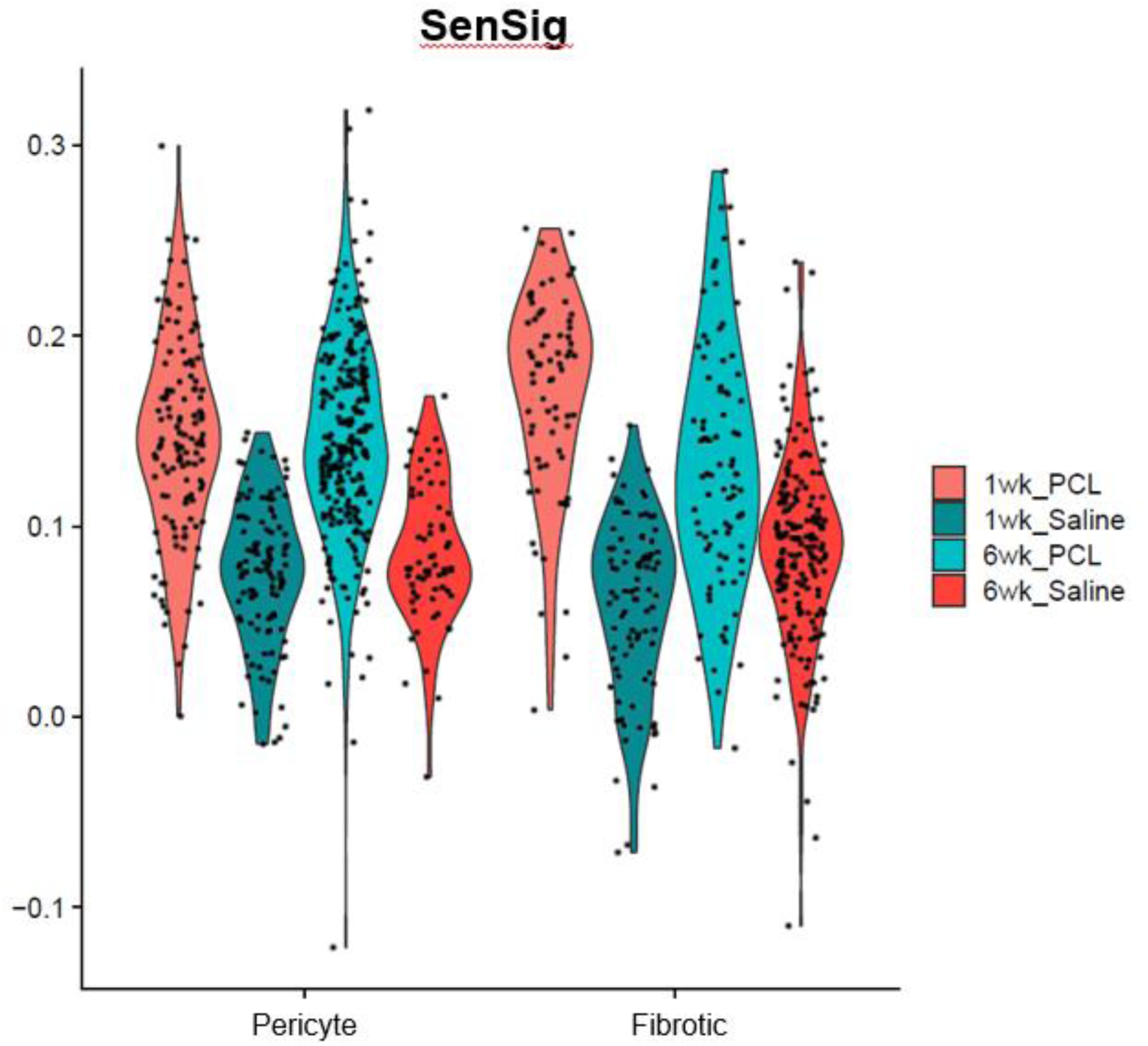
SenSig scores by sample within SenSig high clusters. Violin plots of SenSig for PCL and saline treated animals 1- and 6- weeks post injury split by cluster are shown for the two highest SenSig clusters in the stromal VML scRNAseq data set.

**Supplementary Figure 11.**
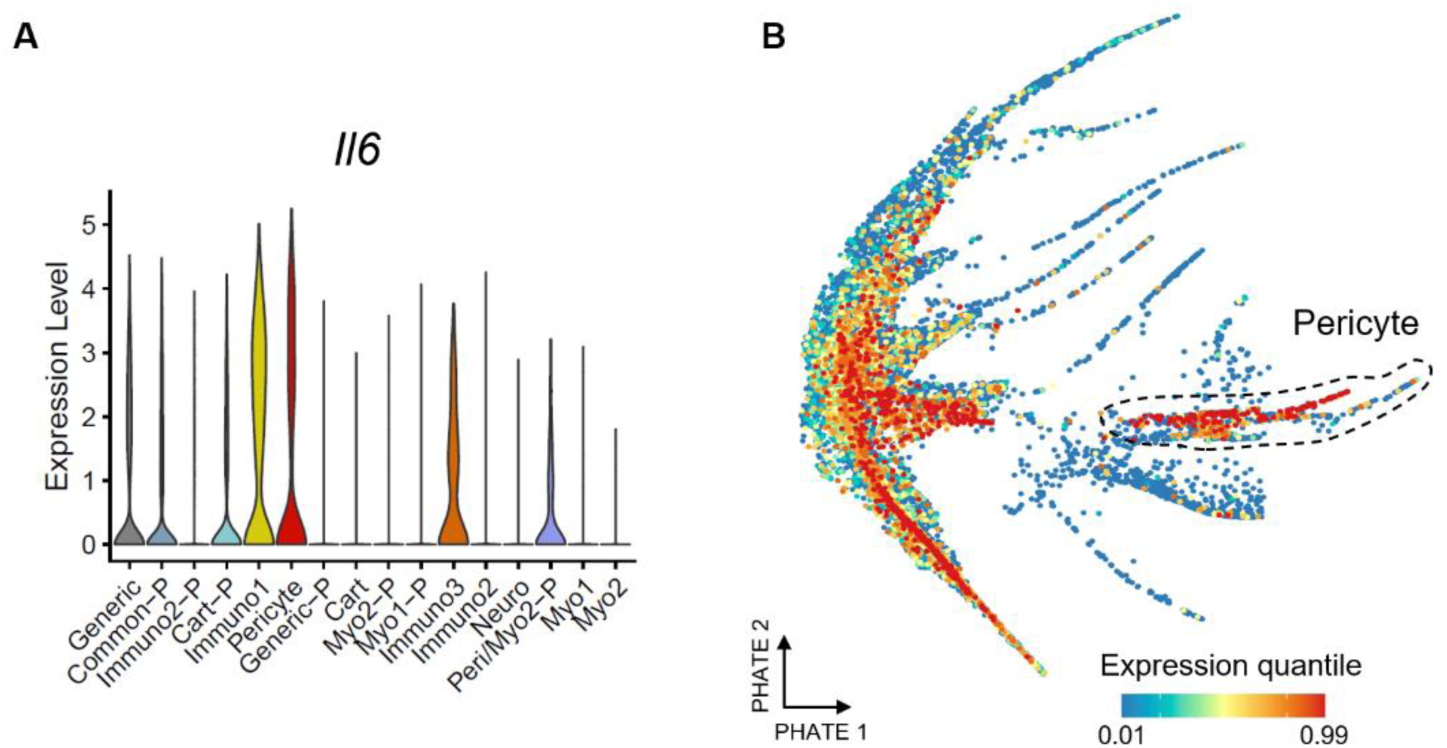
Expression of *Il6* in murine scRNAseq clusters. (A and B) Expression of *Il6* is shown by cluster in a violin plot (A) and feature plot with the pericyte cluster circled (B).

**Supplementary Figure 12.**
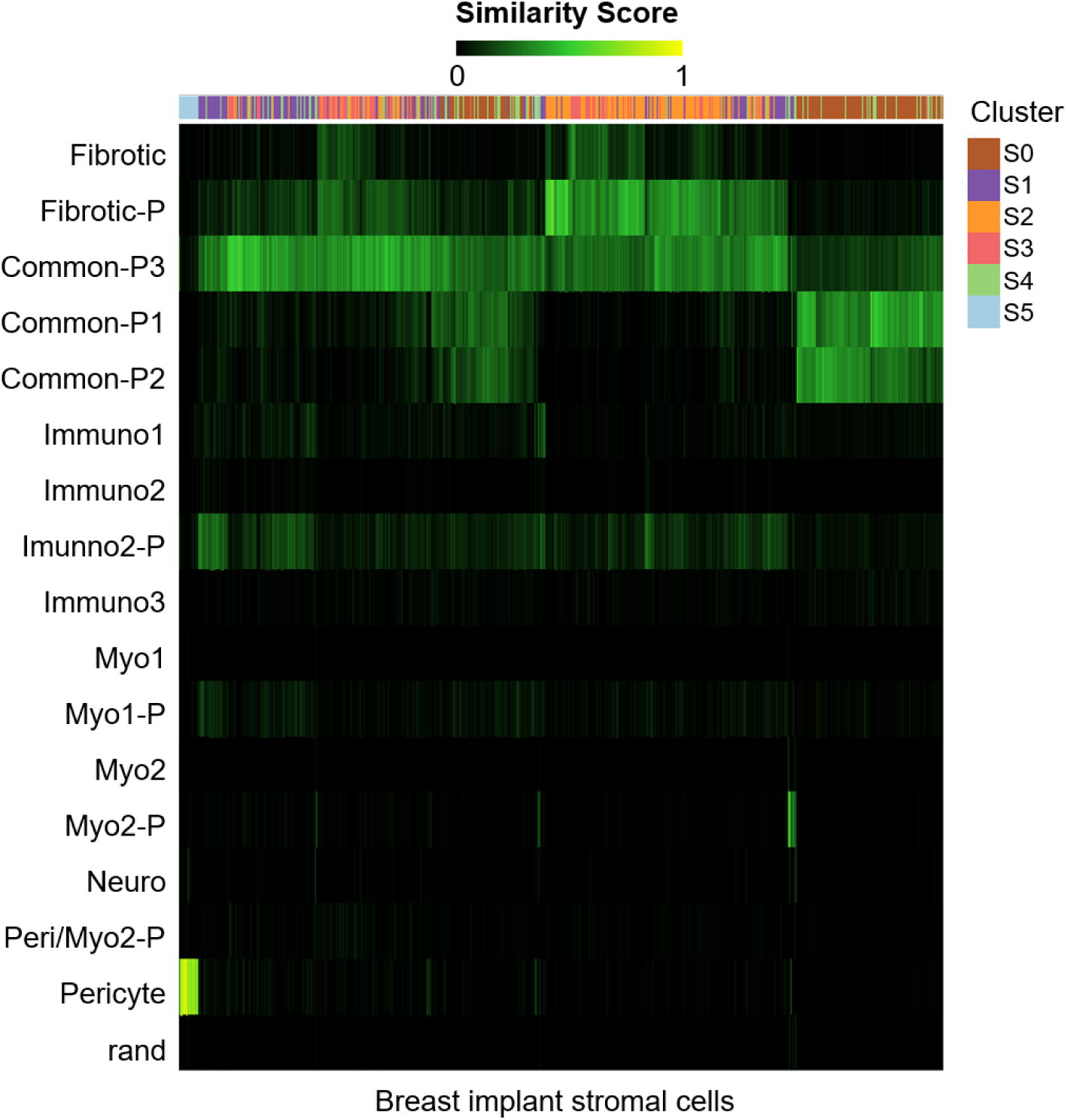
singleCellNet label transfer scores for the fibrotic breast implant stromal cells. Heatmap rows indicate clusters from the murine stromal scRNAseq data set presented in Figure 3. Columns indicate stromal cells computationally isolated from the fibrotic breast implant capsule scRNAseq data set in Figure 4. Higher similarity scores indicate similar transcriptional characteristics between the reference clusters (rows) and the cells in the human fibrotic breast implant data set (column).

**Supplementary Figure 13.**
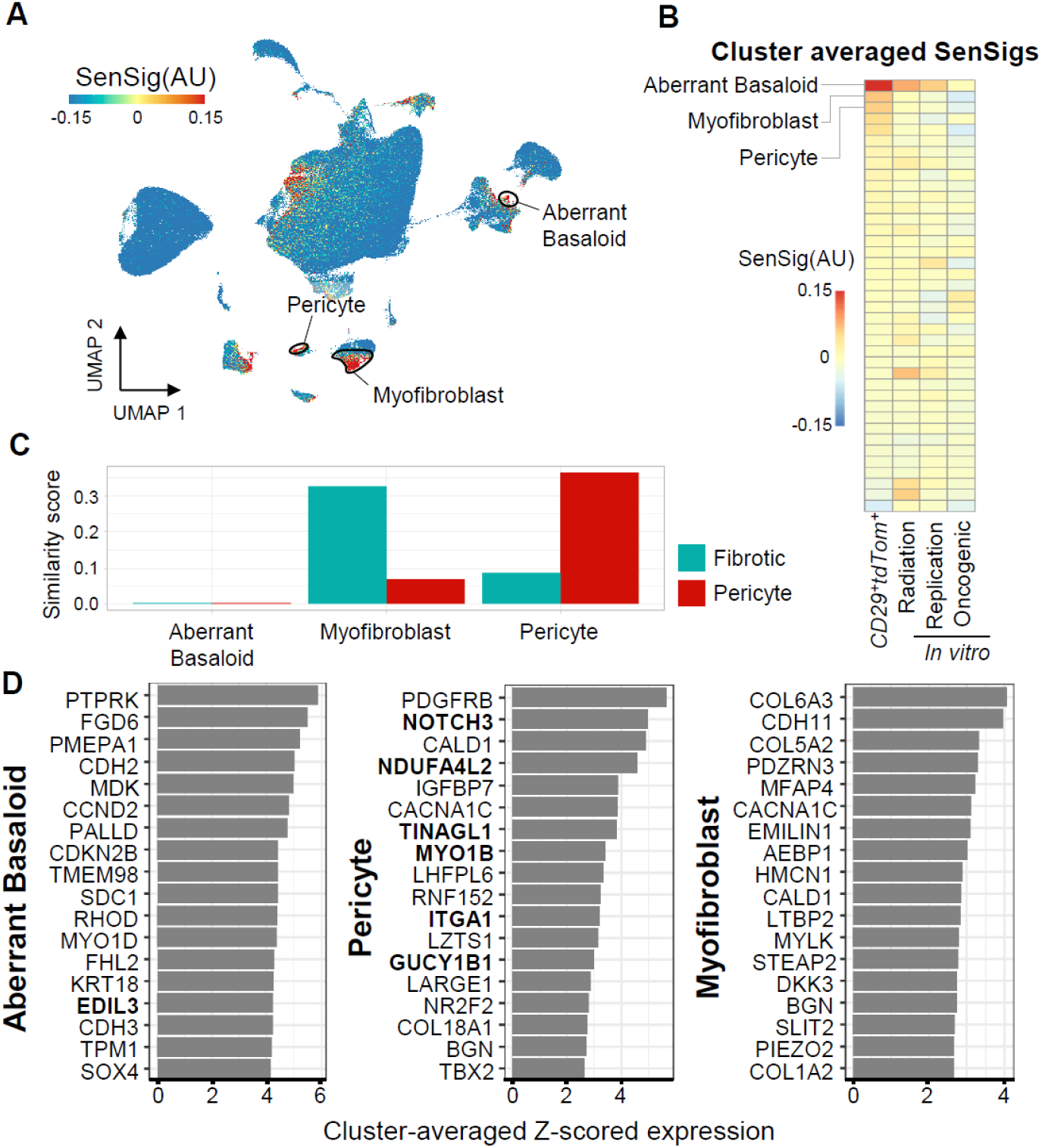
Transfer learning of senescence signature in an additional idiopathic pulmonary fibrosis data set. (A) Feature plots of SenSig calculated using our *in vivo* p16+ reporter mouse. (B) Cluster averages of SenSig using our senescence signature as well as three *in vitro* derived publicly available gene sets. Clustering included in the publicly available data were used and are described in detail in the source publications. (C) Similarity scores using our murine, VML derived scRNAseq clusters as references. Higher similarity score indicates similar gene expression patterns to murine scRNAseq stromal clusters. (D) Genes driving the largest increase in SenSig in the three human stromal clusters with highest average SenSig. Values shown are average z-scored expression across cells in the target cluster. Genes that were also in the top 25 genes driving senescent signature in the murine fibrotic or pericyte clusters are shown in bold.

**Supplementary Figure 14.**
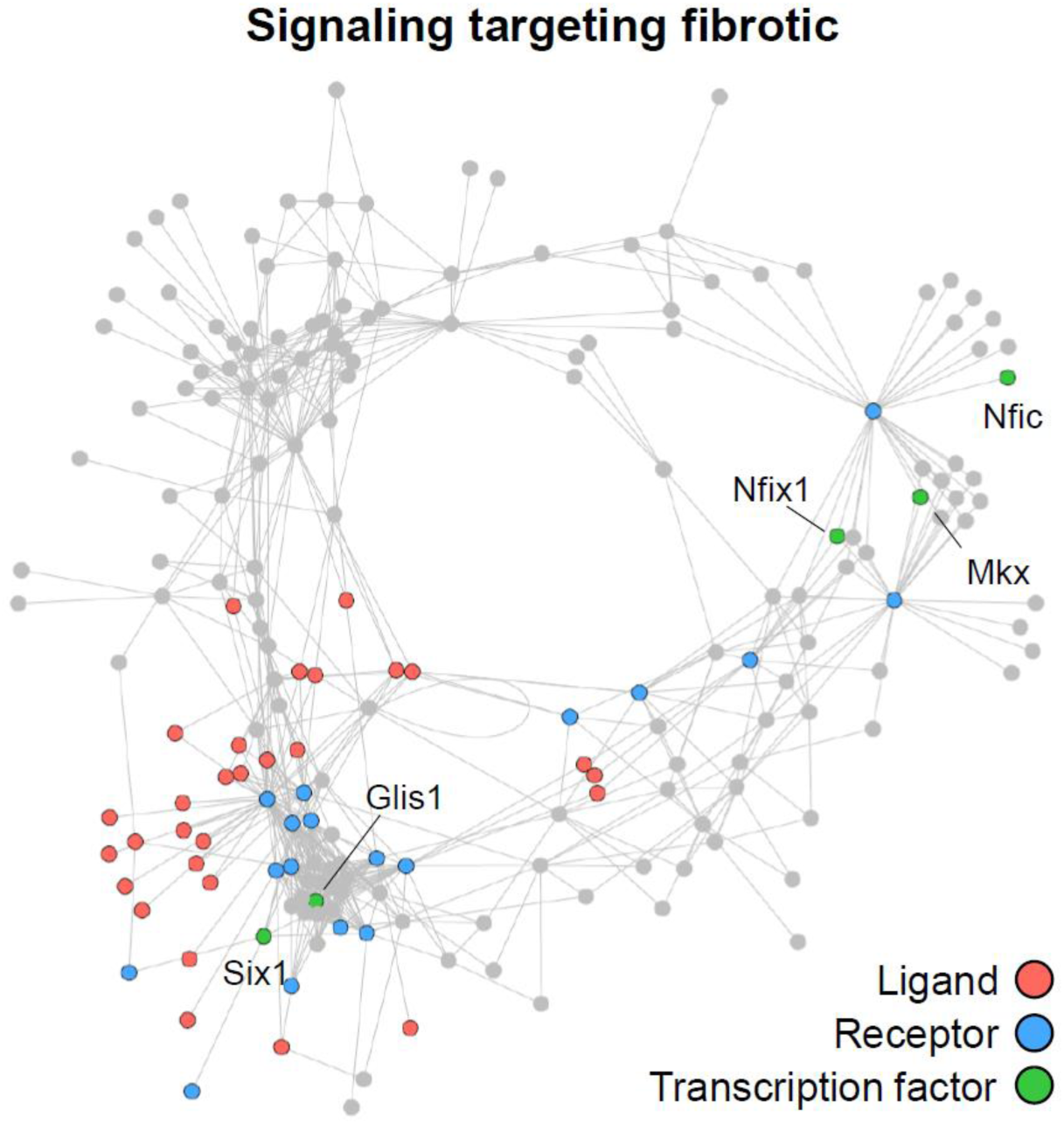
Intercellular signaling patterns targeting fibrotic cells. Predicted signaling patterns targeting fibrotic stromal cells. All signaling connections between ligands, receptors, and their predicted transcription factor targets are shown. Signaling members not related to fibrotic signaling are greyed out. Connections between ligands and receptors in the network are derived from a curated ligand-receptor pairing database. Connections between receptors and transcription factors indicate that expression of the receptor was correlated with activation of the transcription factor in the scRNAseq data set.

**Supplementary Figure 15.**
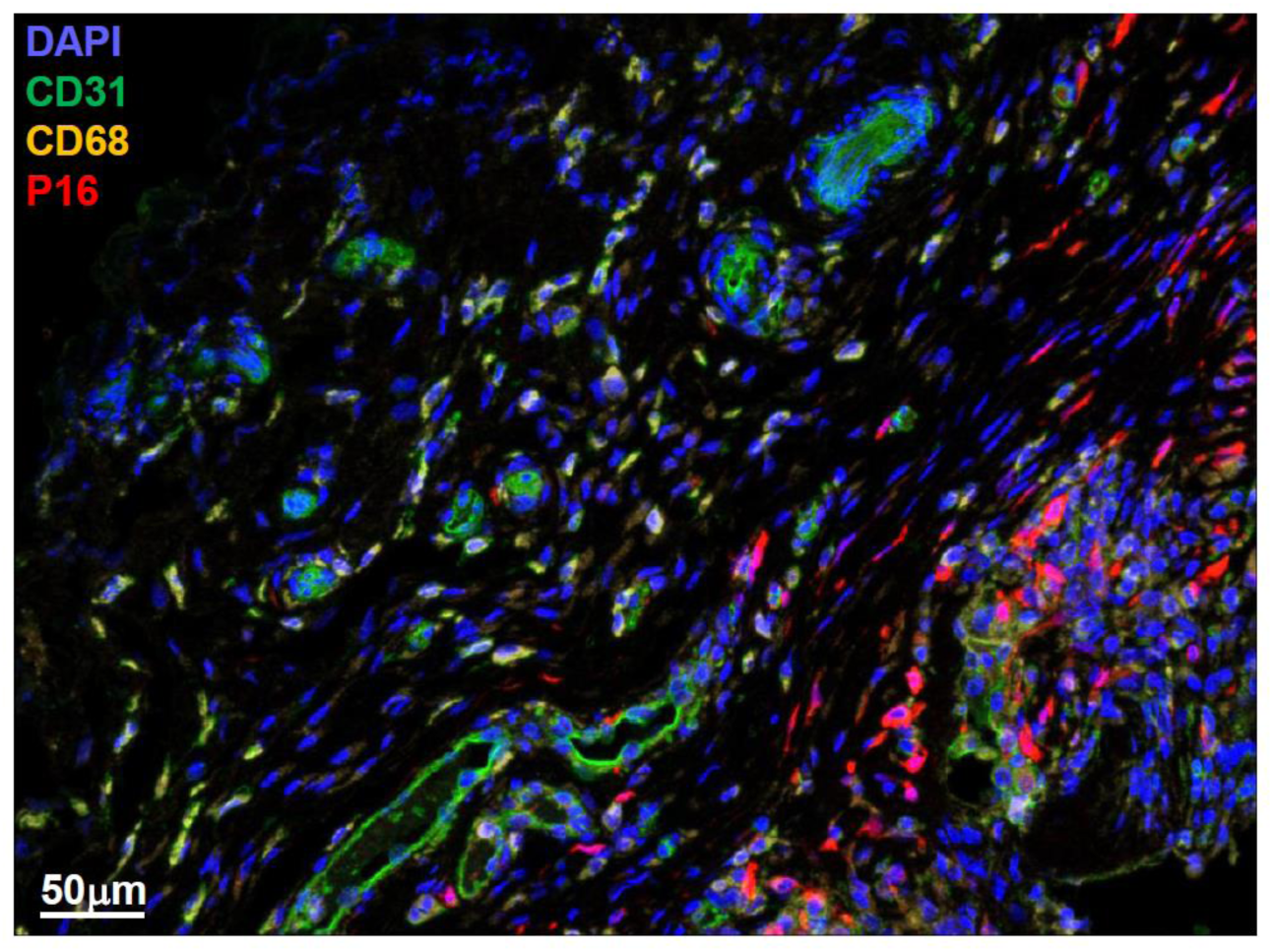
Immunofluorescent staining of myeloid and SnC populations in human fibrotic breast implant capsule. Immunofluorescent staining of P16, macrophage marker CD68, and endothelial cell marker CD31 in a human fibrotic breast implant capsule sample.

