## Supplementary material for "Transfer learning of an *in vivo-*derived senescence signature identifies conserved and tissue-specific senescence across species and diverse pathologies": High Resolution Figures

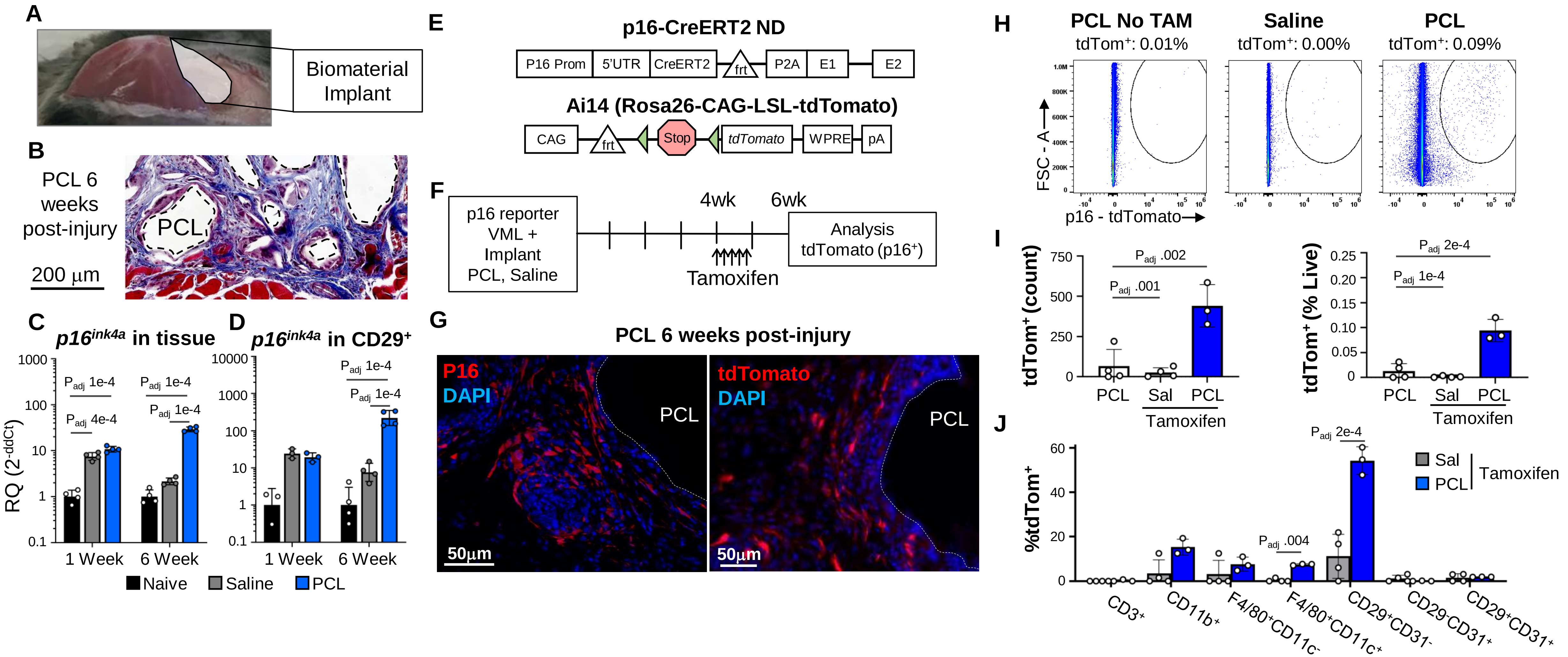
A
Biomaterial Implant
B
PCL 6 weeks post-injury
PCL
200 µm
C
p16ink4a in tissue
RQ (2-ddCt)
Padj 1e-4
Padj 4e-4
Padj 1e-4
Padj 1e-4
1 Week
6 Week
D
p16ink4a in CD29+
Padj 1e-4
Padj 1e-4
1 Week
6 Week
Naive
Saline
PCL
E
p16-CreERT2 ND
P16 Prom
5'UTR
CreERT2
frt
P2A
E1
E2
Ai14 (Rosa26-CAG-LSL-tdTomato)
CAG
frt
Stop
tdTomato
WPRE
pA
F
p16 reporter VML + Implant PCL, Saline
4wk
6wk
Tamoxifen
Analysis tdTomato (p16+)
G
PCL 6 weeks post-injury
P16
DAPI
PCL
50µm
tdTomato
DAPI
PCL
50µm
H
PCL No TAM
tdTom+: 0.01%
Saline
tdTom+: 0.00%
PCL
tdTom+: 0.09%
FSC - A
p16 - tdTomato
I
tdTom+ (count)
Padj .002
Padj .001
PCL
Sal
PCL
Tamoxifen
tdTom+ (% Live)
Padj 2e-4
Padj 1e-4
PCL
Sal
PCL
Tamoxifen
J
%tdTom+
Padj 2e-4
Padj .004
Sal
PCL
Tamoxifen
CD3+
CD11b+
F4/80+CD11c-
F4/80+CD11c+
CD29+CD31-
CD29-CD31+
CD29+CD31+

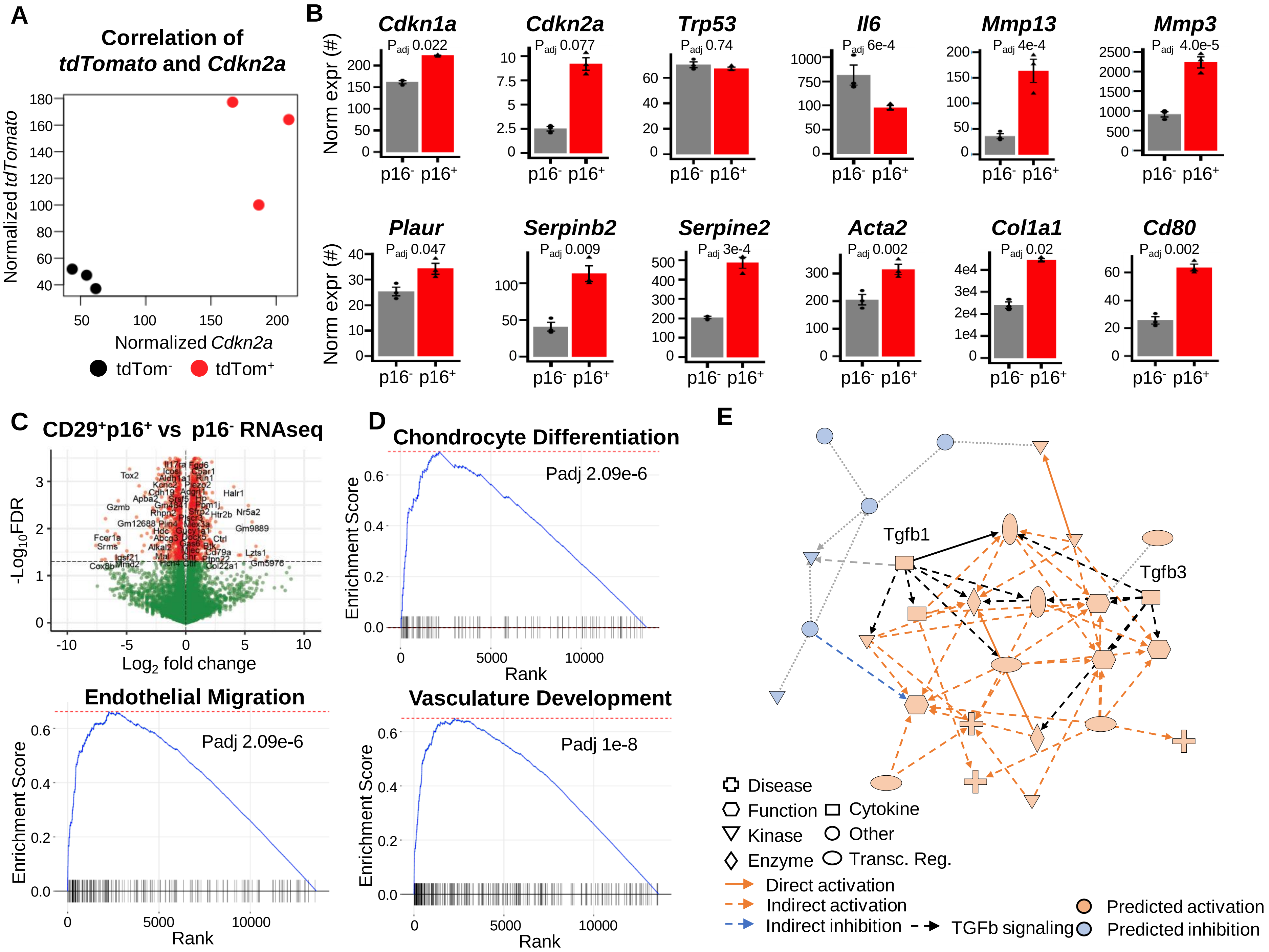
A
Correlation of tdTomato and Cdkn2a
Normalized tdTomato
Normalized Cdkn2a
tdTom-
tdTom+
B
Cdkn1a
Cdkn2a
Trp53
Il6
Mmp13
Mmp3
Plaur
Serpinb2
Serpine2
Acta2
Col1a1
Cd80
Norm expr (#)
p16- p16+
Padj 0.022
Padj 0.077
Padj 0.74
Padj 6e-4
Padj 4e-4
Padj 4.0e-5
Padj 0.047
Padj 0.009
Padj 3e-4
Padj 0.002
Padj 0.02
Padj 0.002
C
CD29+p16+ vs p16- RNAseq
-Log10FDR
Log2 fold change
D
Chondrocyte Differentiation
Padj 2.09e-6
Endothelial Migration
Padj 2.09e-6
Vasculature Development
Padj 1e-8
Enrichment Score
Rank
E
Tgfb1
Tgfb3
Disease
Function
Cytokine
Kinase
Other
Enzyme
Transc. Reg.
Direct activation
Indirect activation
Indirect inhibition
TGFb signaling
Predicted activation
Predicted inhibition

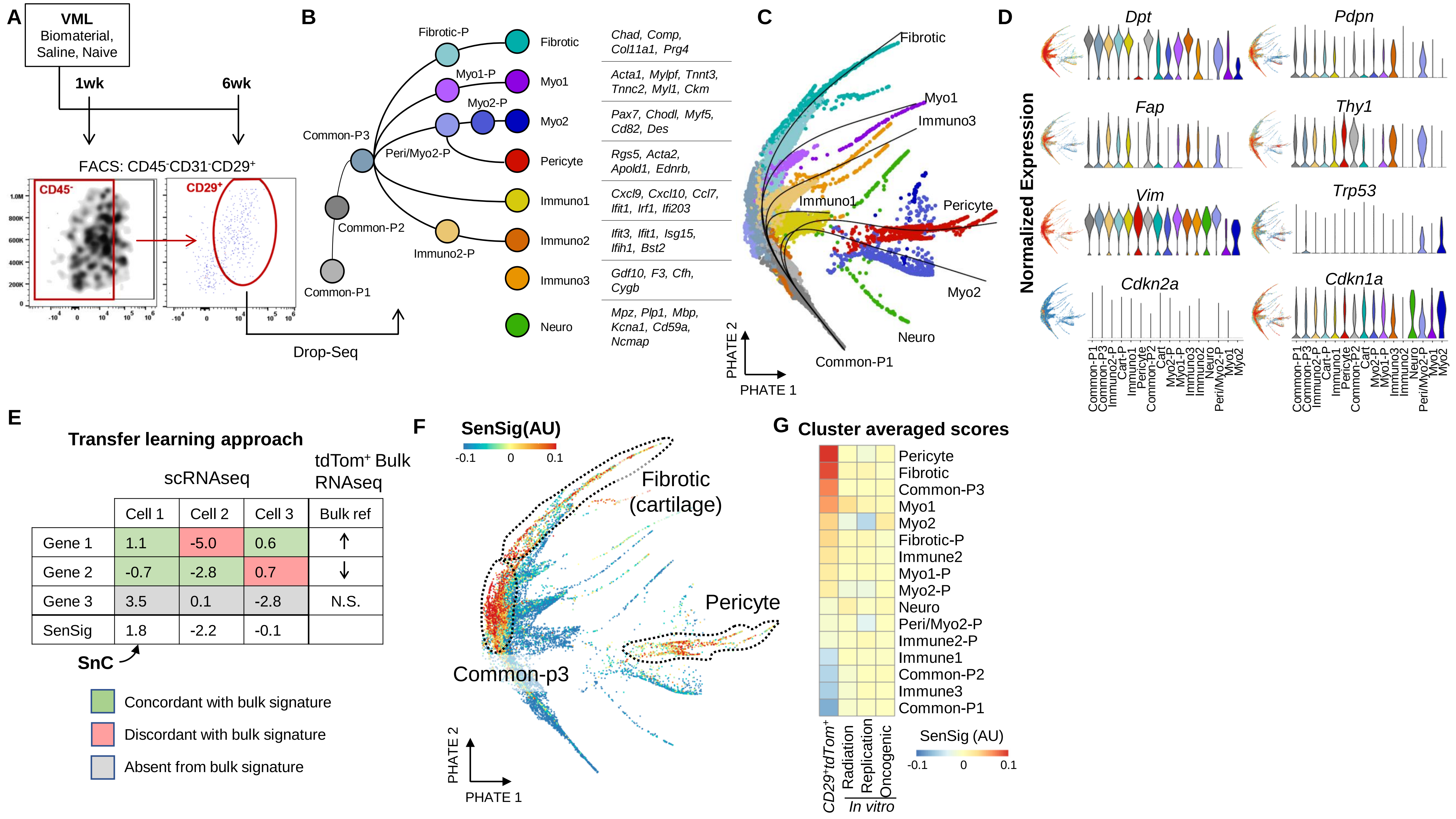
A
VML
Biomaterial,
Saline, Naive
1wk
6wk
FACS: CD45-CD31-CD29+
CD45-
CD29+
Drop-Seq
B
Fibrotic-P
Myo1-P
Myo2-P
Common-P3
Peri/Myo2-P
Common-P2
Immuno2-P
Common-P1
Fibrotic
Myo1
Myo2
Pericyte
Immuno1
Immuno2
Immuno3
Neuro
Chad, Comp, Col11a1, Prg4
Acta1, Mylpf, Tnnt3, Tnnc2, Myl1, Ckm
Pax7, Chodl, Myf5, Cd82, Des
Rgs5, Acta2, Apold1, Ednrb,
Cxcl9, Cxcl10, Ccl7, Ifit1, Irf1, Ifi203
Ifit3, Ifit1, Isg15, Ifih1, Bst2
Gdf10, F3, Cfh, Cygb
Mpz, Plp1, Mbp, Kcna1, Cd59a, Ncmap
C
Fibrotic
Myo1
Immuno3
Immuno1
Pericyte
Myo2
Neuro
Common-P1
PHATE 2
PHATE 1
D
Normalized Expression
Dpt
Pdpn
Fap
Thy1
Vim
Trp53
Cdkn2a
Cdkn1a
E
Transfer learning approach
scRNAseq
tdTom+ Bulk RNAseq
Cell 1 | Cell 2 | Cell 3 | Bulk ref
Gene 1 | 1.1 | -5.0 | 0.6 | ↑
Gene 2 | -0.7 | -2.8 | 0.7 | ↓
Gene 3 | 3.5 | 0.1 | -2.8 | N.S.
SenSig | 1.8 | -2.2 | -0.1
SnC
Concordant with bulk signature
Discordant with bulk signature
Absent from bulk signature
F
SenSig(AU)
-0.1 0 0.1
Fibrotic
(cartilage)
Pericyte
Common-p3
PHATE 2
PHATE 1
G
Cluster averaged scores
Pericyte
Fibrotic
Common-P3
Myo1
Myo2
Fibrotic-P
Immune2
Myo1-P
Myo2-P
Neuro
Peri/Myo2-P
Immune2-P
Immune1
Common-P2
Immune3
Common-P1
CD29+tdTom+
Radiation
Replication
Oncogenic
In vitro
SenSig (AU)
-0.1 0 0.1

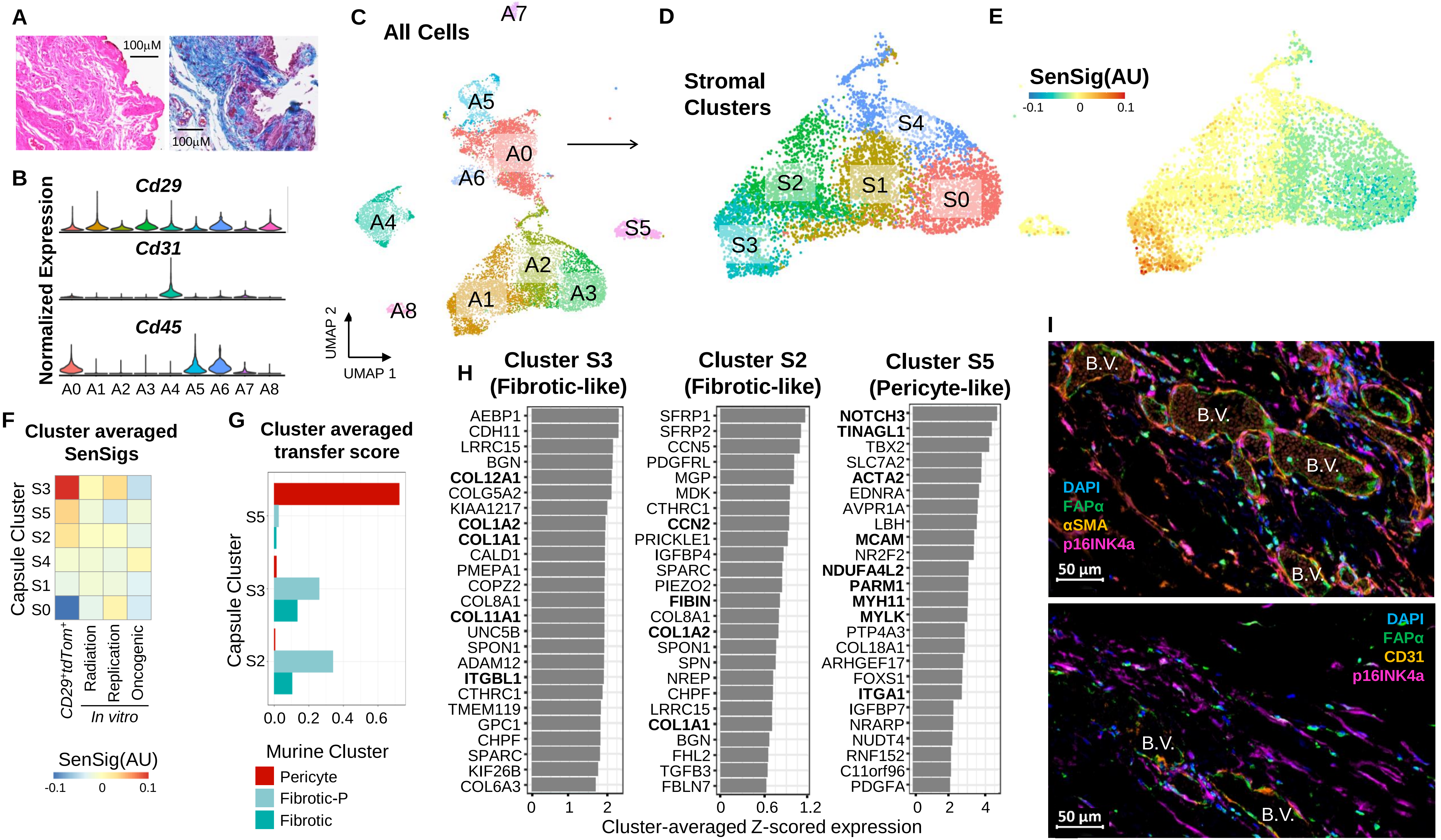
A
100μM
100μM
B
Normalized Expression
Cd29
Cd31
Cd45
A0 A1 A2 A3 A4 A5 A6 A7 A8
C
All Cells
A7
A5
A0
A6
A4
S5
A2
A1
A3
A8
UMAP 2
UMAP 1
D
Stromal Clusters
S4
S2
S1
S0
S3
E
SenSig(AU)
-0.1 0 0.1
F
Cluster averaged SenSigs
Capsule Cluster
S3
S5
S2
S4
S1
S0
CD29+tdTom+
Radiation
Replication
Oncogenic
In vitro
SenSig(AU)
-0.1 0 0.1
G
Cluster averaged transfer score
Capsule Cluster
S5
S3
S2
0.0 0.2 0.4 0.6
Murine Cluster
Pericyte
Fibrotic-P
Fibrotic
H
Cluster S3 (Fibrotic-like)
AEBP1
CDH11
LRRC15
BGN
COL12A1
COLG5A2
KIAA1217
COL1A2
COL1A1
CALD1
PMEPA1
COPZ2
COL8A1
COL11A1
UNC5B
SPON1
ADAM12
ITGBL1
CTHRC1
TMEM119
GPC1
CHPF
SPARC
KIF26B
COL6A3
0 1 2
Cluster S2 (Fibrotic-like)
SFRP1
SFRP2
CCN5
PDGFRL
MGP
MDK
CTHRC1
CCN2
PRICKLE1
IGFBP4
SPARC
PIEZO2
FIBIN
COL8A1
COL1A2
SPON1
SPN
NREP
CHPF
LRRC15
COL1A1
BGN
FHL2
TGFB3
FBLN7
0 0.6 1.2
Cluster S5 (Pericyte-like)
NOTCH3
TINAGL1
TBX2
SLC7A2
ACTA2
EDNRA
AVPR1A
LBH
MCAM
NR2F2
NDUFA4L2
PARM1
MYH11
MYLK
PTP4A3
COL18A1
ARHGEF17
FOXS1
ITGA1
IGFBP7
NRARP
NUDT4
RNF152
C11orf96
PDGFA
0 2 4
Cluster-averaged Z-scored expression
I
B.V.
B.V.
B.V.
B.V.
DAPI
FAPα
αSMA
p16INK4a
50 μm
DAPI
FAPα
CD31
p16INK4a
B.V.
B.V.
50 μm

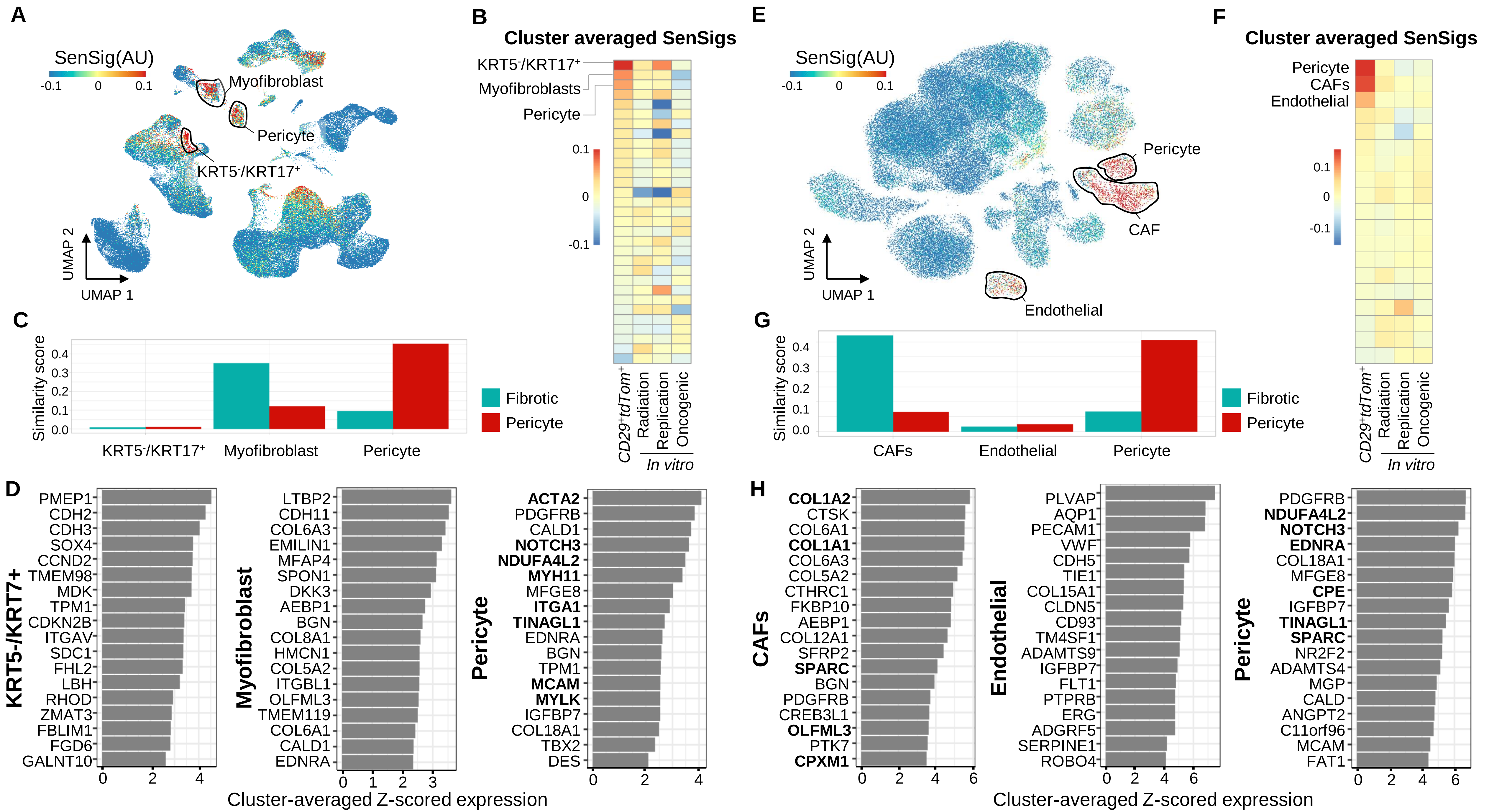
A
SenSig(AU)
-0.1 0 0.1
Myofibroblast
Pericyte
KRT5-/KRT17+
UMAP 2
UMAP 1
B
Cluster averaged SenSigs
KRT5-/KRT17+
Myofibroblasts
Pericyte
CD29+tdTom+
Radiation
Replication
Oncogenic
In vitro
C
Similarity score
KRT5-/KRT17+
Myofibroblast
Pericyte
Fibrotic
Pericyte
D
KRT5-/KRT7+
PMEP1 CDH2 CDH3 SOX4 CCND2 TMEM98 MDK TPM1 CDKN2B ITGAV SDC1 FHL2 LBH RHOD ZMAT3 FBLIM1 FGD6 GALNT10
Myofibroblast
LTBP2 CDH11 COL6A3 EMILIN1 MFAP4 SPON1 DKK3 AEBP1 BGN COL8A1 HMCN1 COL5A2 ITGBL1 OLFML3 TMEM119 COL6A1 CALD1 EDNRA
Pericyte
ACTA2 PDGFRB CALD1 NOTCH3 NDUFA4L2 MYH11 MFGE8 ITGA1 TINAGL1 EDNRA BGN TPM1 MCAM MYLK IGFBP7 COL18A1 TBX2 DES
Cluster-averaged Z-scored expression
E
SenSig(AU)
Pericyte
CAF
Endothelial
UMAP 2
UMAP 1
F
Cluster averaged SenSigs
Pericyte
CAFs
Endothelial
CD29+tdTom+
Radiation
Replication
Oncogenic
In vitro
G
Similarity score
CAFs
Endothelial
Pericyte
Fibrotic
Pericyte
H
CAFs
COL1A2 CTSK COL6A1 COL1A1 COL6A3 COL5A2 CTHRC1 FKBP10 AEBP1 COL12A1 SFRP2 SPARC BGN PDGFRB CREB3L1 OLFML3 PTK7 CPXM1
Endothelial
PLVAP AQP1 PECAM1 VWF CDH5 TIE1 COL15A1 CLDN5 CD93 TM4SF1 ADAMTS9 IGFBP7 FLT1 PTPRB ERG ADGRF5 SERPINE1 ROBO4
Pericyte
PDGFRB NDUFA4L2 NOTCH3 EDNRA COL18A1 MFGE8 CPE IGFBP7 TINAGL1 SPARC NR2F2 ADAMTS4 MGP CALD ANGPT2 C11orf96 MCAM FAT1
Cluster-averaged Z-scored expression

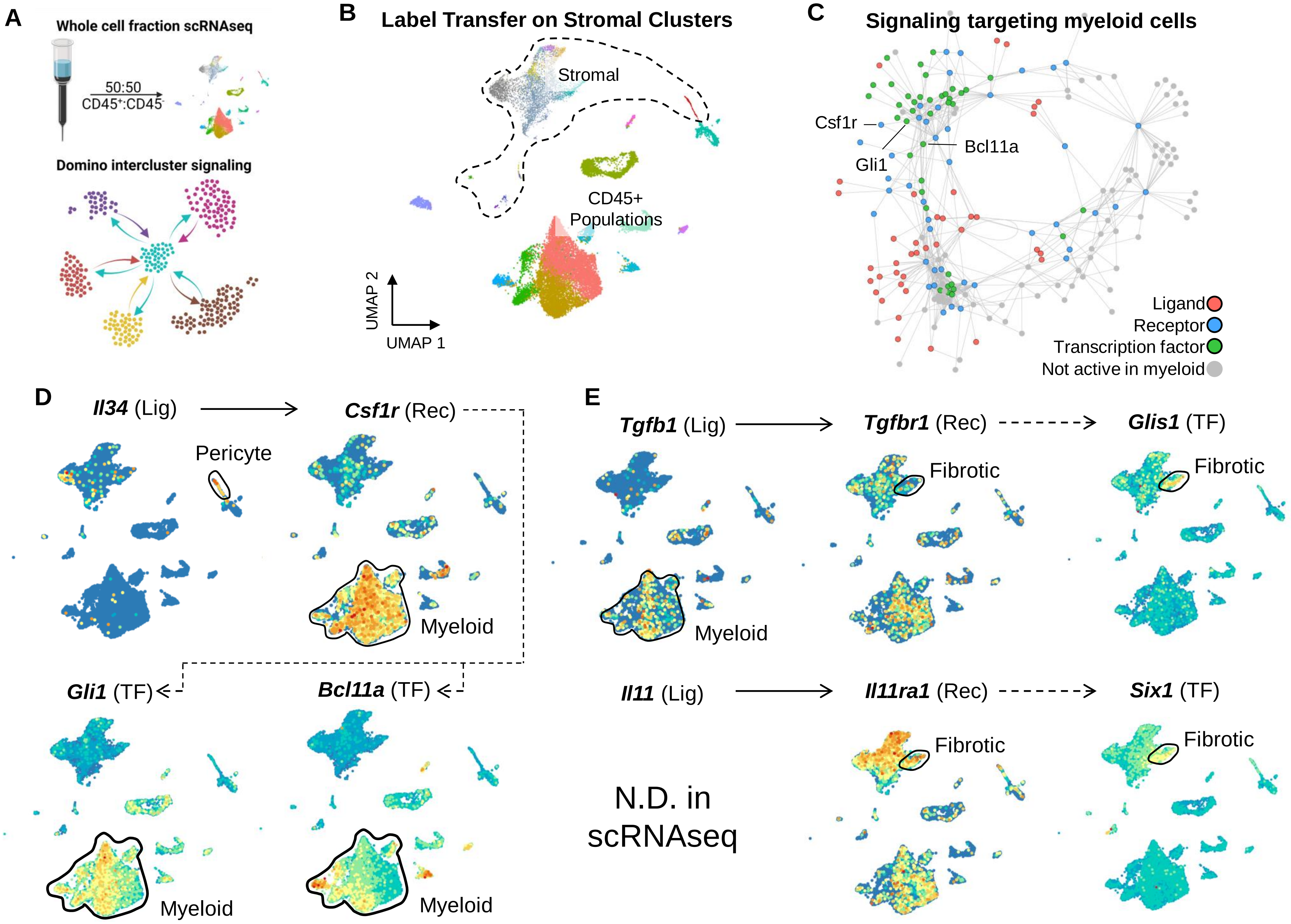
A
Whole cell fraction scRNAseq
50:50
CD45+:CD45-
Domino intercluster signaling
B
Label Transfer on Stromal Clusters
Stromal
CD45+ Populations
UMAP 2
UMAP 1
C
Signaling targeting myeloid cells
Csf1r
Gli1
Bcl11a
Ligand
Receptor
Transcription factor
Not active in myeloid
D
Il34 (Lig)
Csf1r (Rec)
Pericyte
Myeloid
Gli1 (TF)
Bcl11a (TF)
Myeloid
Myeloid
E
Tgfb1 (Lig)
Tgfbr1 (Rec)
Glis1 (TF)
Fibrotic
Fibrotic
Myeloid
Il11 (Lig)
Il11ra1 (Rec)
Six1 (TF)
Fibrotic
Fibrotic
N.D. in scRNAseq

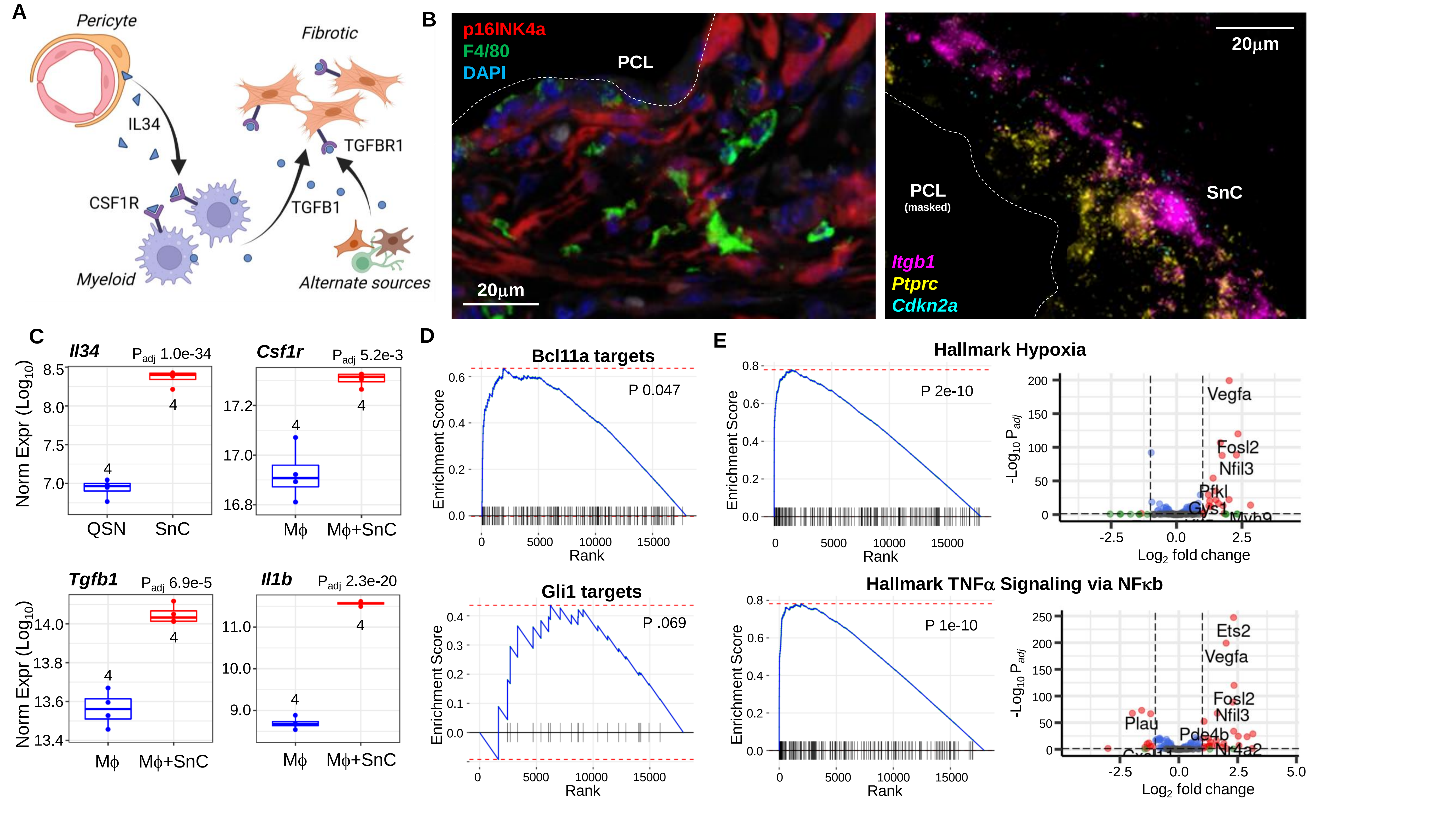
A
Pericyte
IL34
CSF1R
Myeloid
Fibrotic
TGFBR1
TGFB1
Alternate sources
B
p16INK4a
F4/80
DAPI
PCL
20μm
PCL
(masked)
SnC
Itgb1
Ptprc
Cdkn2a
20μm
C
Il34
Padj 1.0e-34
Csf1r
Padj 5.2e-3
Norm Expr (Log10)
QSN
SnC
Mϕ
Mϕ+SnC
Tgfb1
Padj 6.9e-5
Il1b
Padj 2.3e-20
D
Bcl11a targets
P 0.047
Enrichment Score
Rank
Gli1 targets
P .069
E
Hallmark Hypoxia
P 2e-10
Vegfa
Fosl2
Nfil3
Pfkl
Gys1
-Log10 Padj
Log2 fold change
Hallmark TNFα Signaling via NFκb
P 1e-10
Ets2
Vegfa
Fosl2
Nfil3
Plau
Pde4b

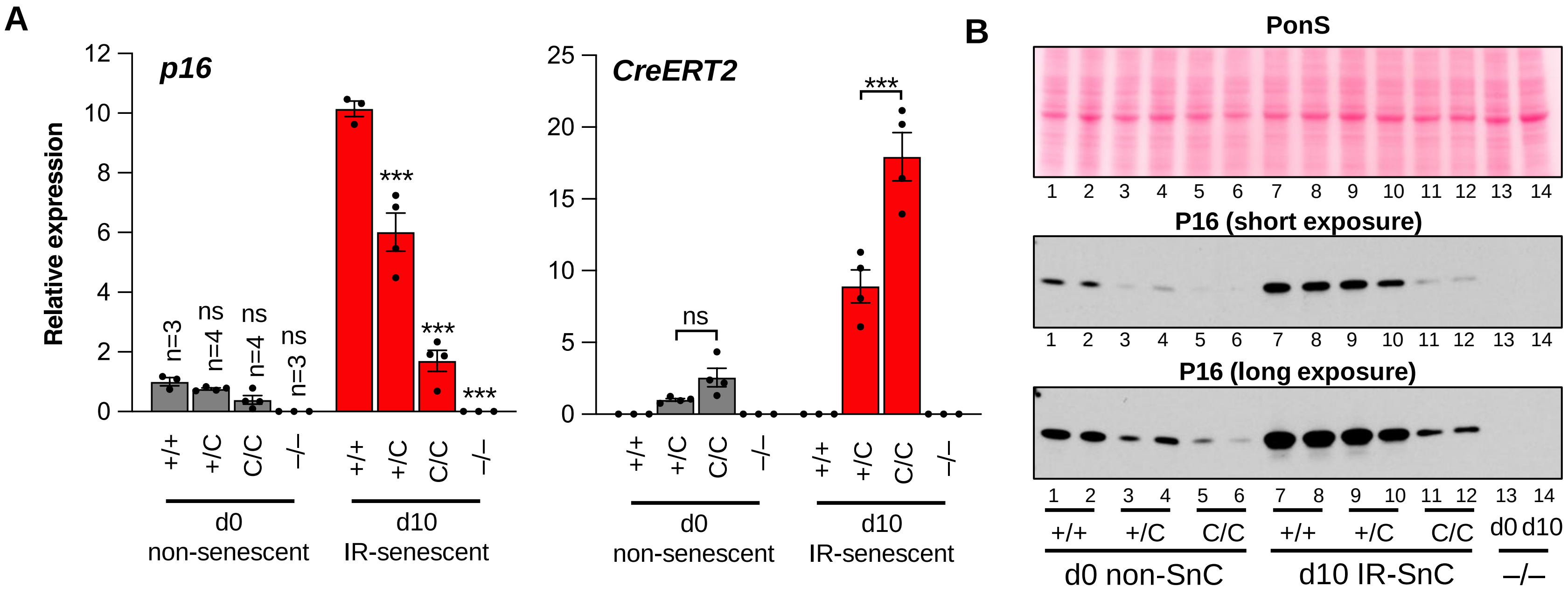
A
p16
Relative expression
n=3
n=4
n=4
n=3
ns
ns
ns
***
***
***
+/+
+/C
C/C
–/–
d0
non-senescent
d10
IR-senescent
CreERT2
ns
***
B
PonS
P16 (short exposure)
P16 (long exposure)
1 2 3 4 5 6 7 8 9 10 11 12 13 14
+/+
+/C
C/C
d0
d10
d0 non-SnC
d10 IR-SnC
–/–

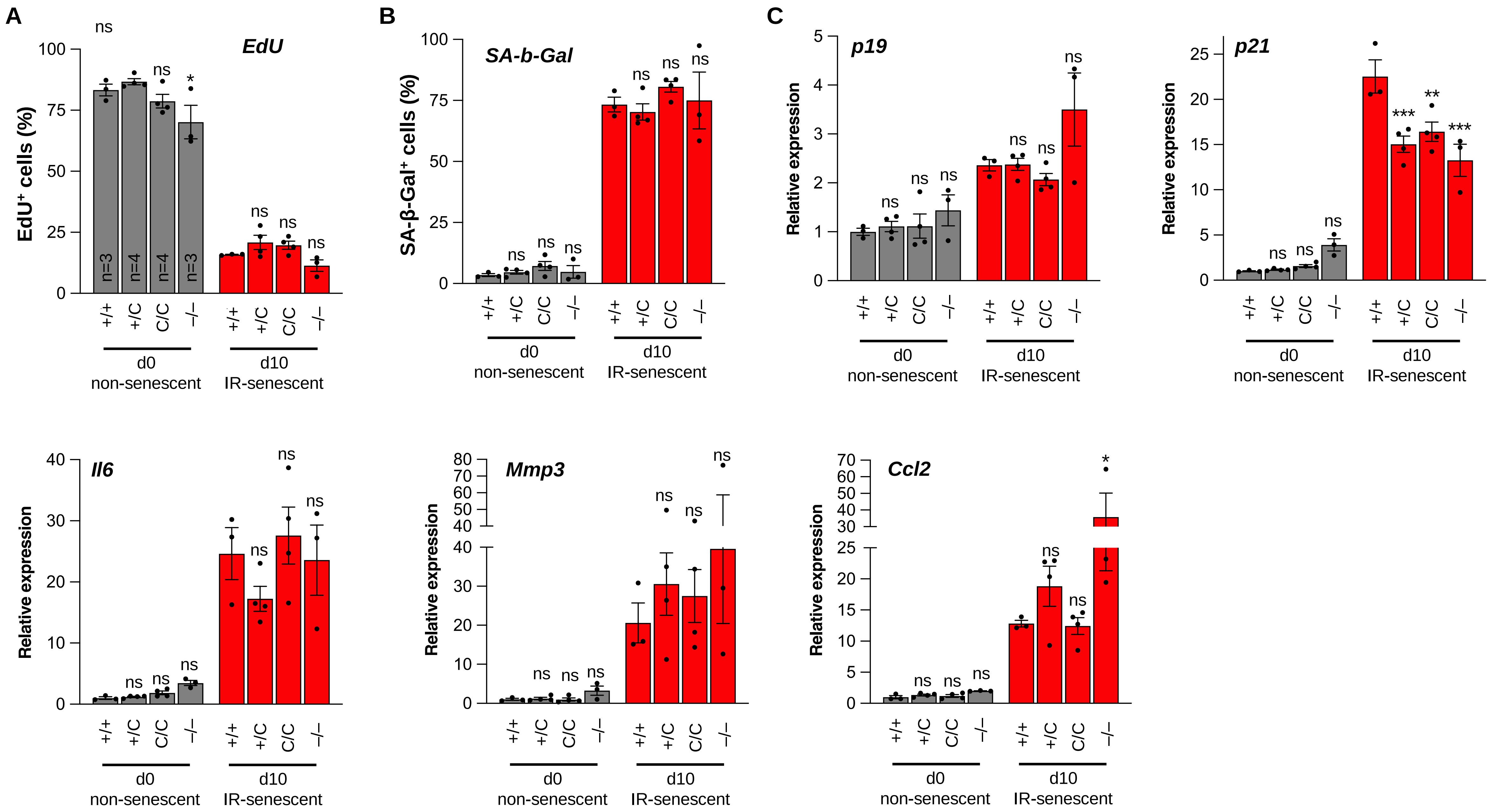

A
EdU
EdU+ cells (%)
n=3
n=4
n=4
n=3
ns
*
B
SA-b-Gal
SA-β-Gal+ cells (%)
C
p19
p21
Il6
Mmp3
Ccl2
Relative expression
+/+
+/C
C/C
–/–
d0
non-senescent
d10
IR-senescent
**
***

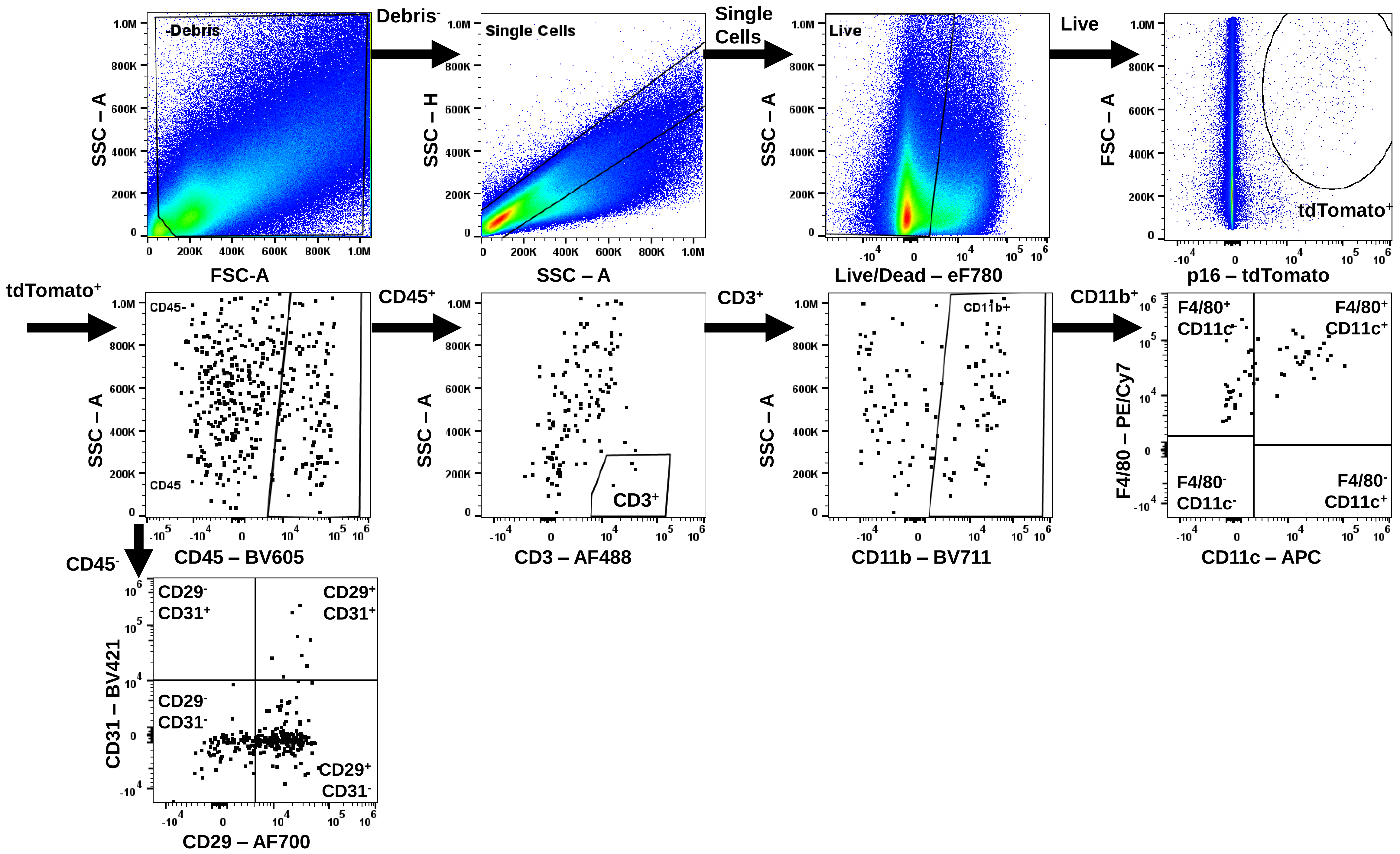

-Debris
Debris-
Single Cells
Single Cells
Live
Live
tdTomato+
tdTomato+
CD45-
CD45
CD45+
CD3+
CD3+
CD11b+
CD11b+
F4/80+ CD11c-
F4/80+ CD11c+
F4/80- CD11c-
F4/80- CD11c+
CD45-
CD29- CD31+
CD29+ CD31+
CD29- CD31-
CD29+ CD31-
FSC-A
SSC – A
SSC – H
Live/Dead – eF780
FSC – A
p16 – tdTomato
CD45 – BV605
CD3 – AF488
CD11b – BV711
F4/80 – PE/Cy7
CD11c – APC
CD31 – BV421
CD29 – AF700

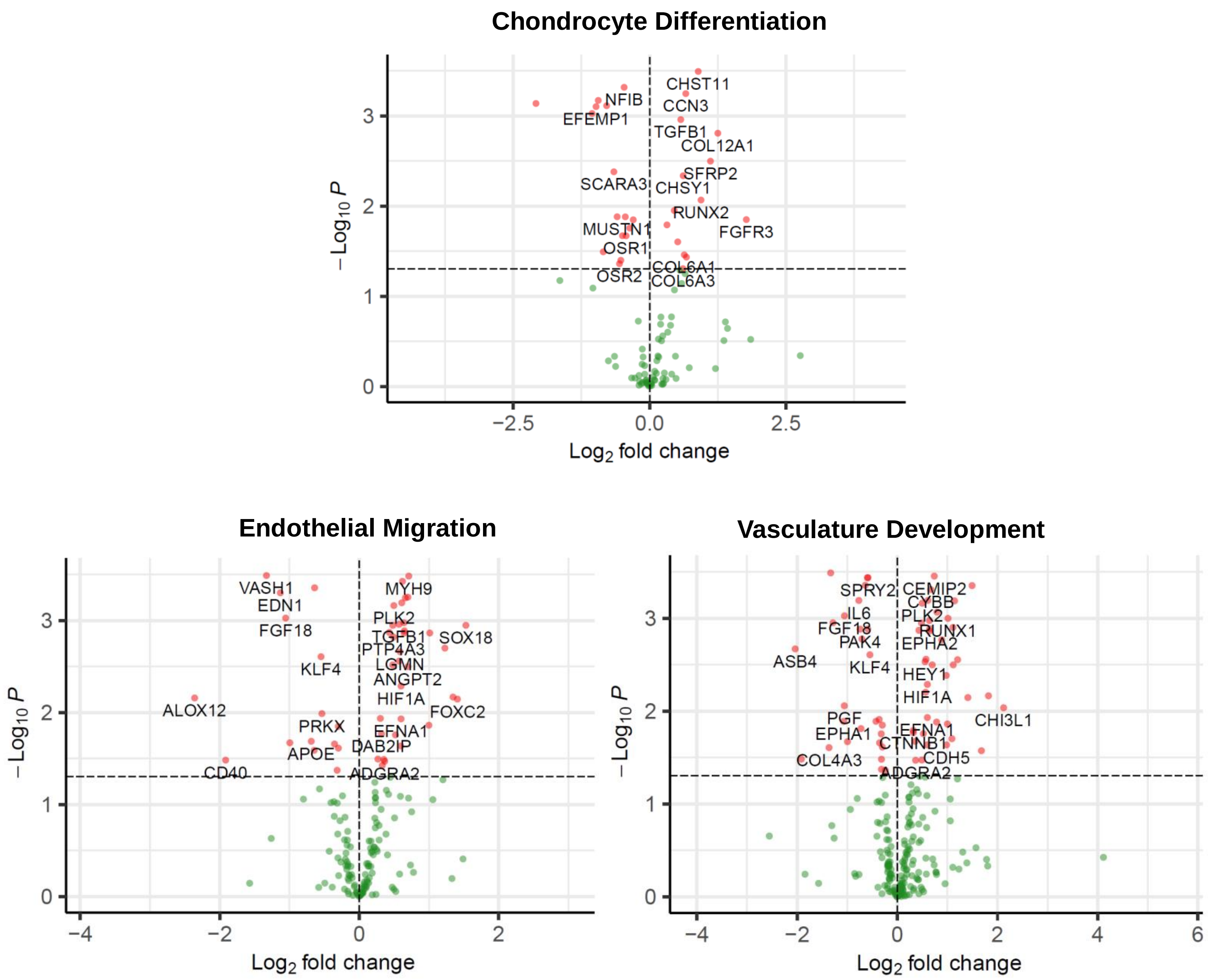
Chondrocyte Differentiation
CHST11
NFIB
CCN3
EFEMP1
TGFB1
COL12A1
SFRP2
SCARA3
CHSY1
RUNX2
FGFR3
MUSTN1
OSR1
COL6A1
OSR2
COL6A3
−2.5
0.0
2.5
Log2 fold change
−Log10 P
Endothelial Migration
VASH1
MYH9
EDN1
PLK2
FGF18
TGFB1
SOX18
PTP4A3
KLF4
LGMN
ANGPT2
HIF1A
ALOX12
FOXC2
PRKX
EFNA1
DAB2IP
APOE
CD40
ADGRA2
−4
−2
0
2
Log2 fold change
−Log10 P
Vasculature Development
SPRY2
CEMIP2
CYBB
IL6
PLK2
FGF18
RUNX1
PAK4
EPHA2
ASB4
KLF4
HEY1
HIF1A
CHI3L1
PGF
EFNA1
EPHA1
CTNNB1
CDH5
COL4A3
ADGRA2
−4
−2
0
2
4
6
Log2 fold change
−Log10 P

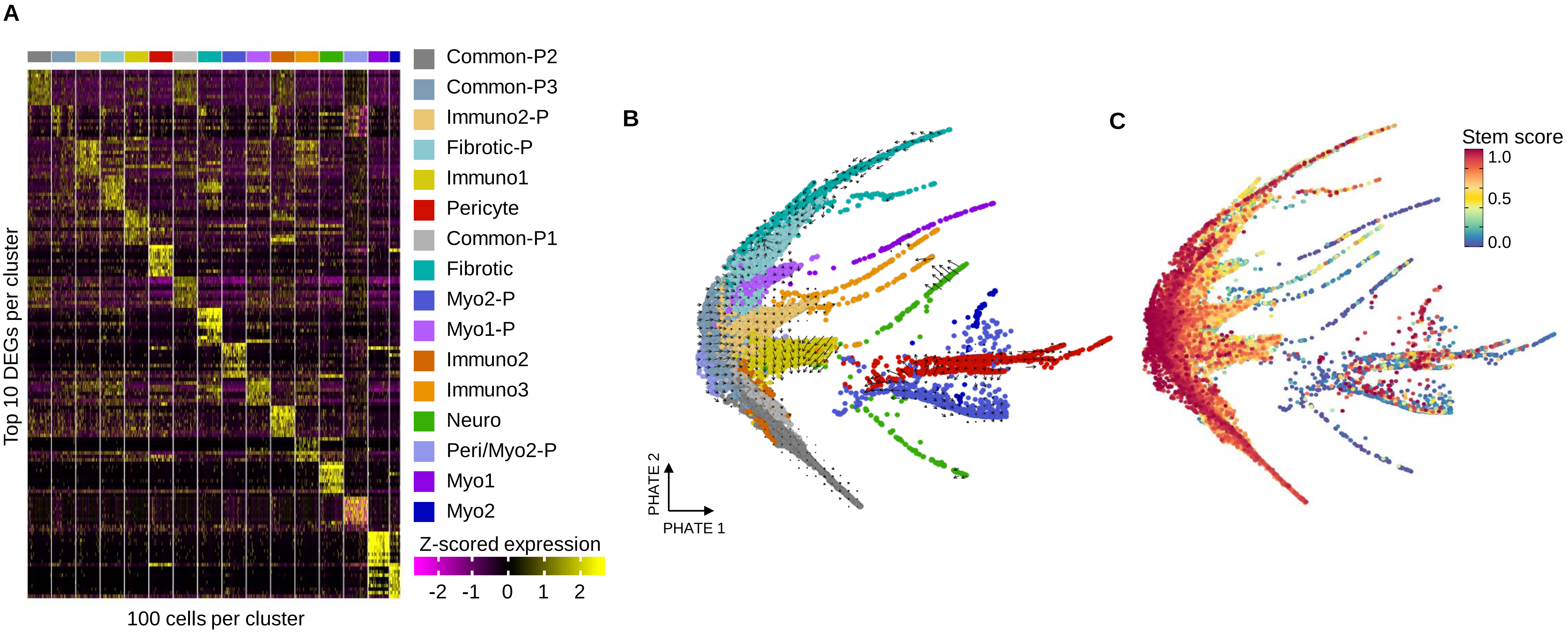

A
Common-P2
Common-P3
Immuno2-P
Fibrotic-P
Immuno1
Pericyte
Common-P1
Fibrotic
Myo2-P
Myo1-P
Immuno2
Immuno3
Neuro
Peri/Myo2-P
Myo1
Myo2
Z-scored expression
-2 -1 0 1 2
Top 10 DEGs per cluster
100 cells per cluster
B
PHATE 2
PHATE 1
C
Stem score
1.0
0.5
0.0

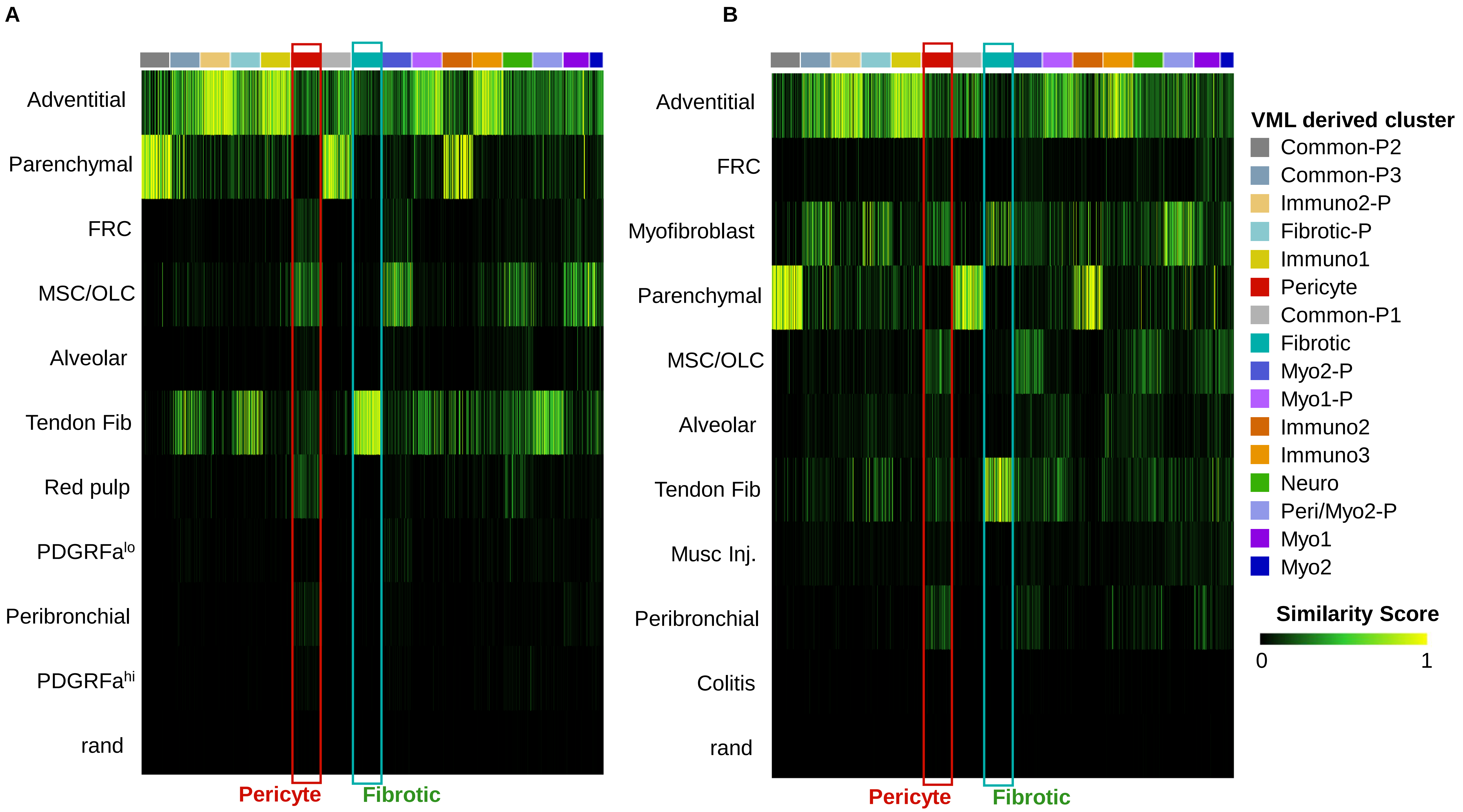
A
Adventitial
Parenchymal
FRC
MSC/OLC
Alveolar
Tendon Fib
Red pulp
PDGFRa^lo
Peribronchial
PDGFRa^hi
rand
Pericyte
Fibrotic
B
Adventitial
FRC
Myofibroblast
Parenchymal
MSC/OLC
Alveolar
Tendon Fib
Musc Inj.
Peribronchial
Colitis
rand
Pericyte
Fibrotic
VML derived cluster
Common-P2
Common-P3
Immuno2-P
Fibrotic-P
Immuno1
Pericyte
Common-P1
Fibrotic
Myo2-P
Myo1-P
Immuno2
Immuno3
Neuro
Peri/Myo2-P
Myo1
Myo2
Similarity Score
0
1

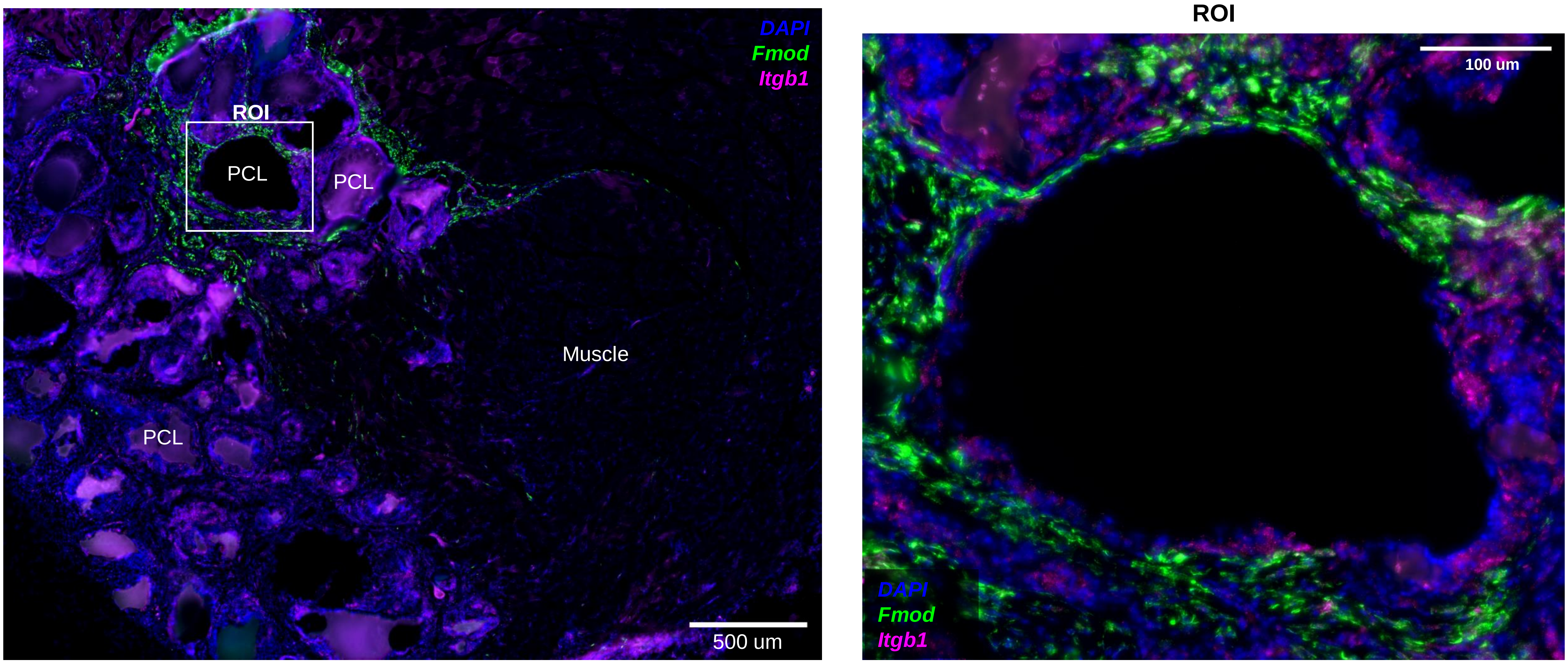
DAPI
Fmod
Itgb1
ROI
PCL
PCL
PCL
Muscle
500 um
ROI
100 um
DAPI
Fmod
Itgb1

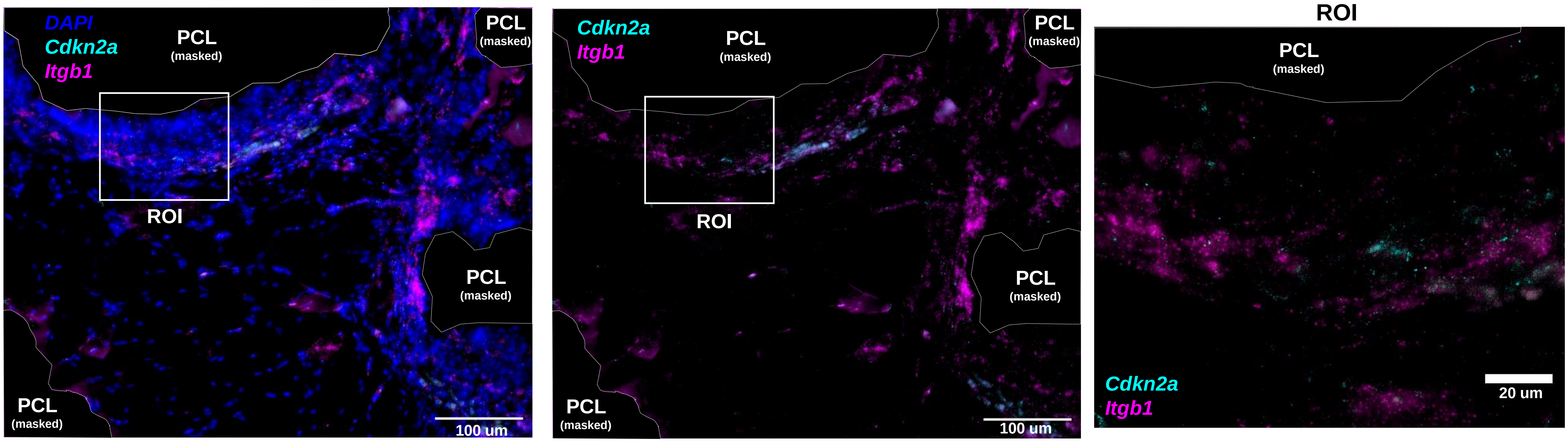
DAPI
Cdkn2a
Itgb1
PCL
(masked)
ROI
100 um
Cdkn2a
Itgb1
ROI
100 um
ROI
PCL
(masked)
Cdkn2a
Itgb1
20 um

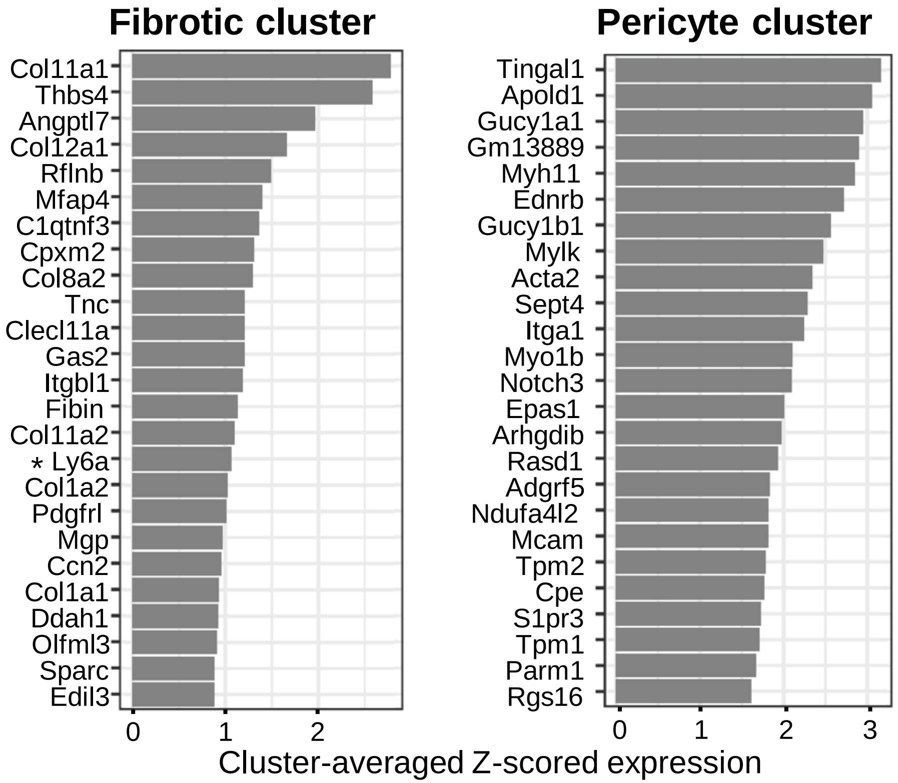


* Negative in both scRNAseq and bulk RNAseq data sets.

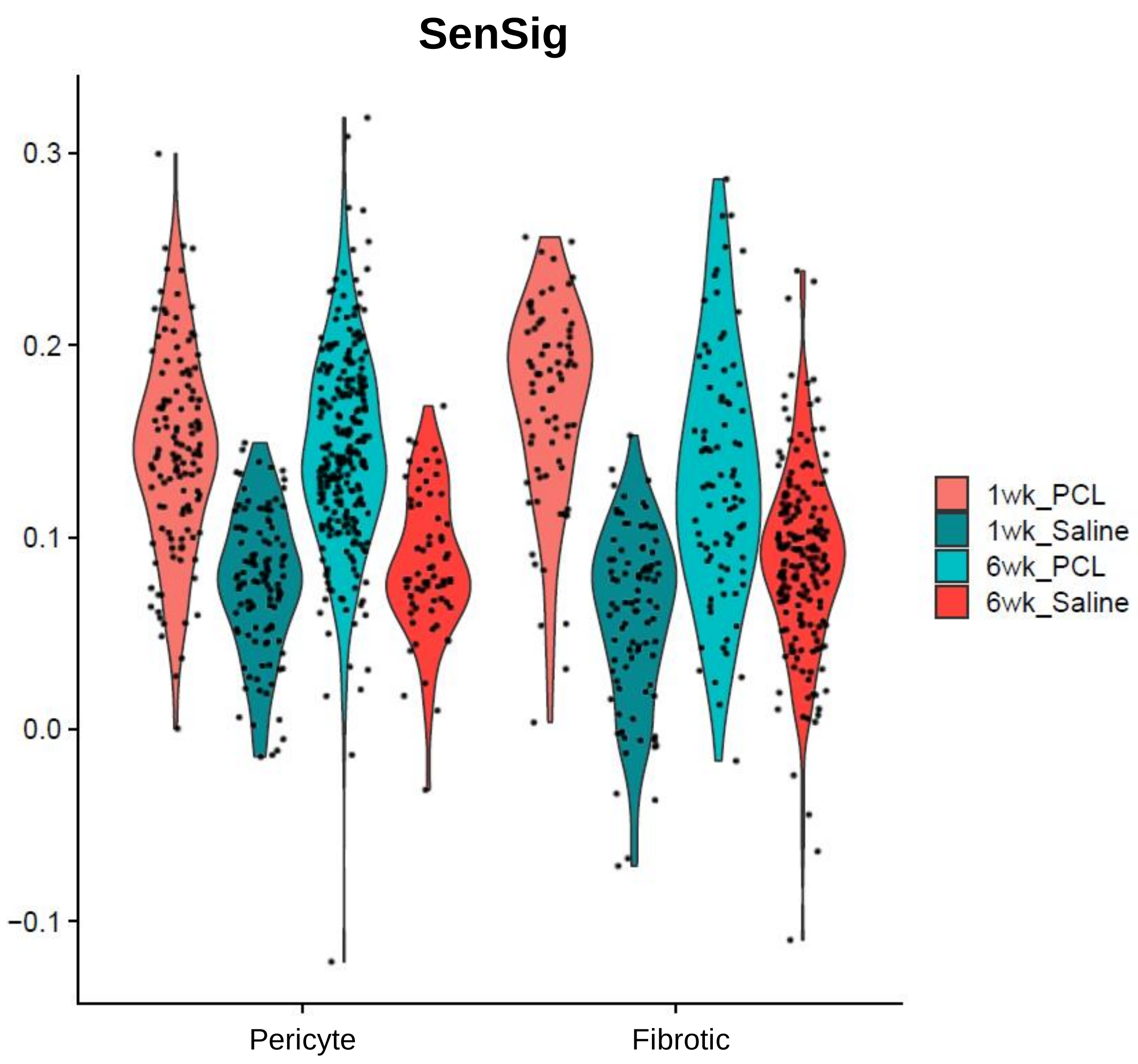
SenSig
0.3
0.2
0.1
0.0
−0.1
Pericyte
Fibrotic
1wk_PCL
1wk_Saline
6wk_PCL
6wk_Saline

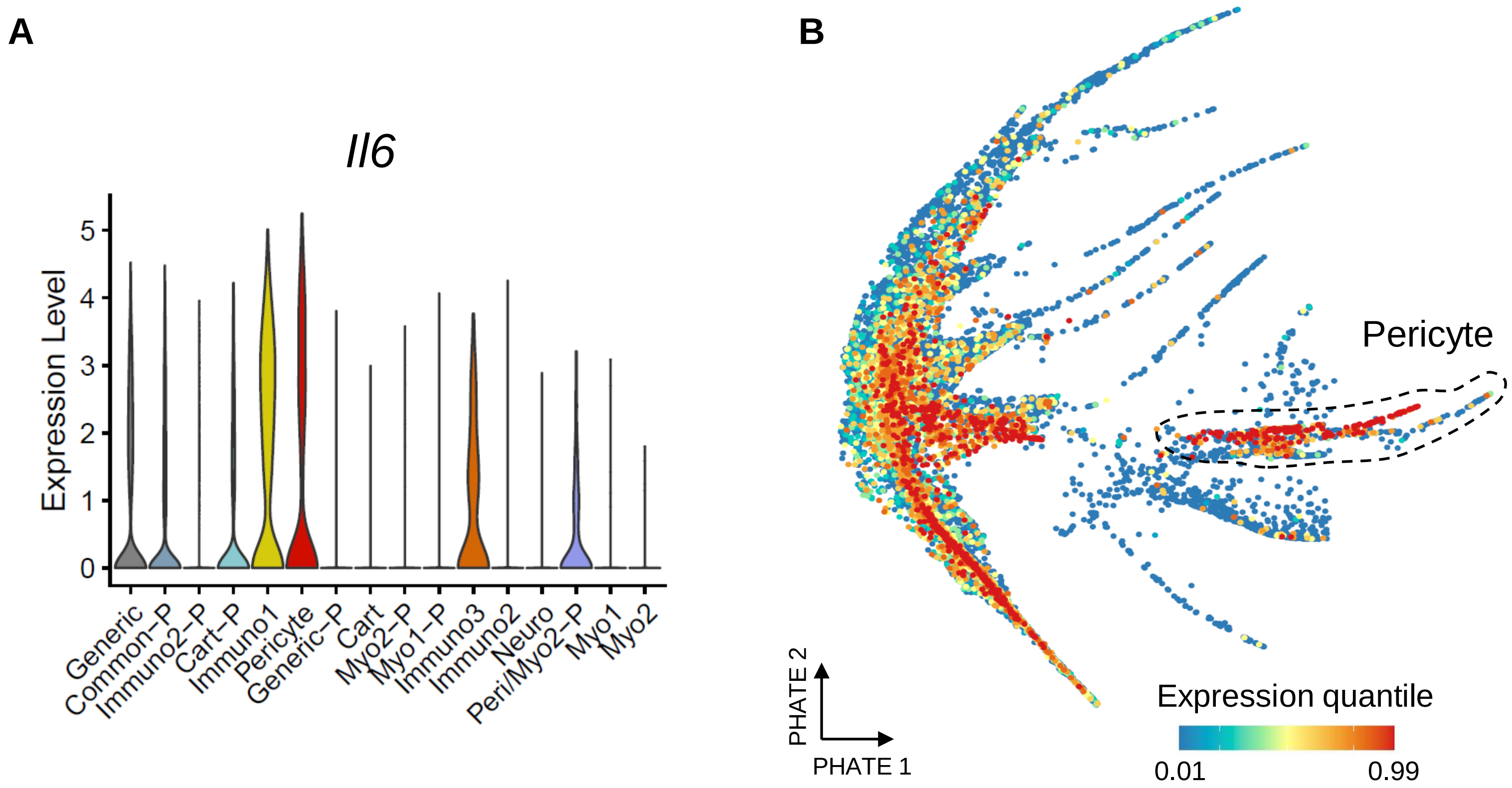
A
Il6
Expression Level
5
4
3
2
1
0
Generic
Common−P
Immuno2−P
Cart−P
Immuno1
Pericyte
Generic−P
Cart
Myo2−P
Myo1−P
Immuno3
Immuno2
Neuro
Peri/Myo2−P
Myo1
Myo2
B
Pericyte
PHATE 2
PHATE 1
Expression quantile
0.01
0.99

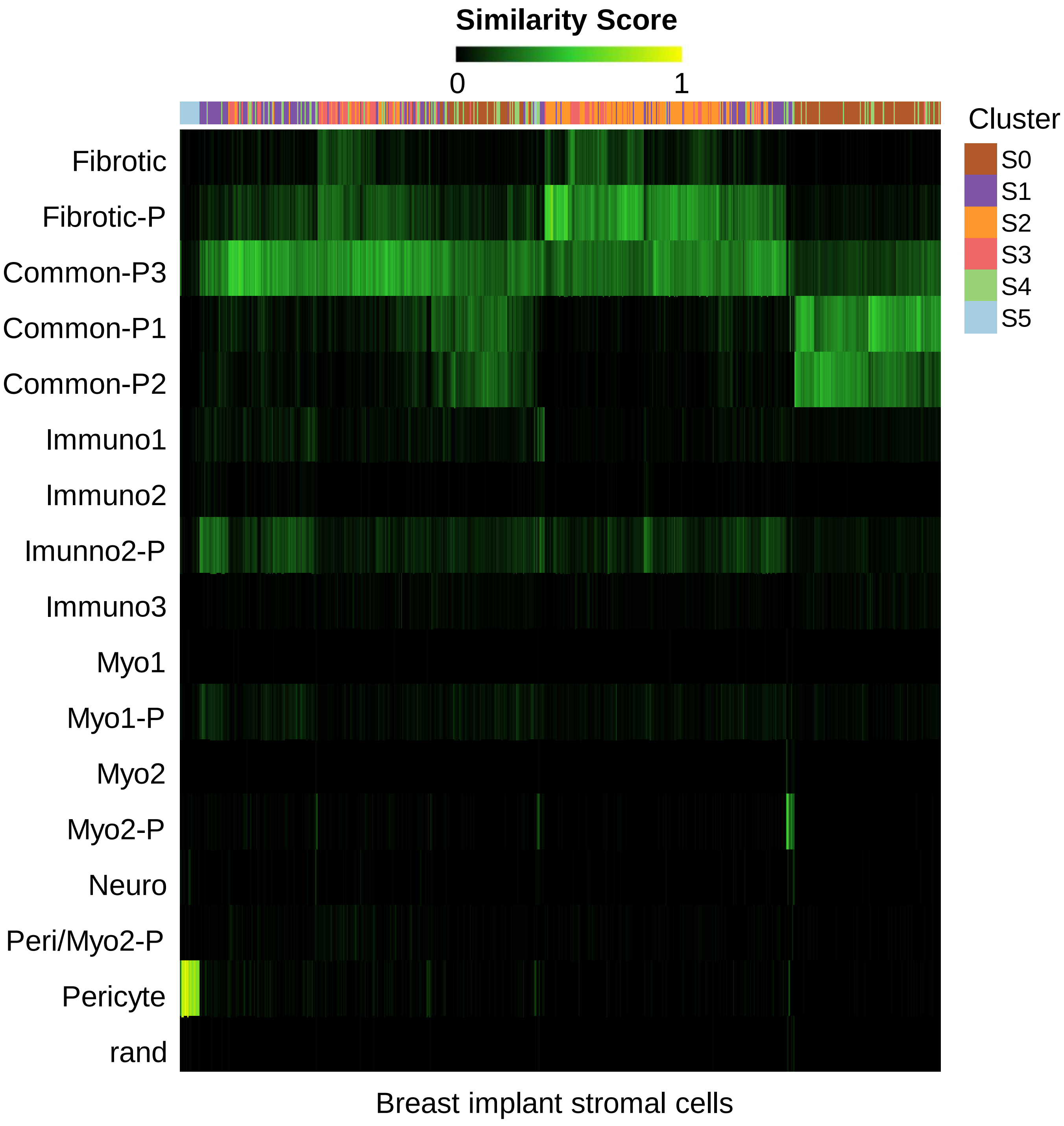
Similarity Score
0
1
Cluster
S0
S1
S2
S3
S4
S5
Fibrotic
Fibrotic-P
Common-P3
Common-P1
Common-P2
Immuno1
Immuno2
Imunno2-P
Immuno3
Myo1
Myo1-P
Myo2
Myo2-P
Neuro
Peri/Myo2-P
Pericyte
rand
Breast implant stromal cells

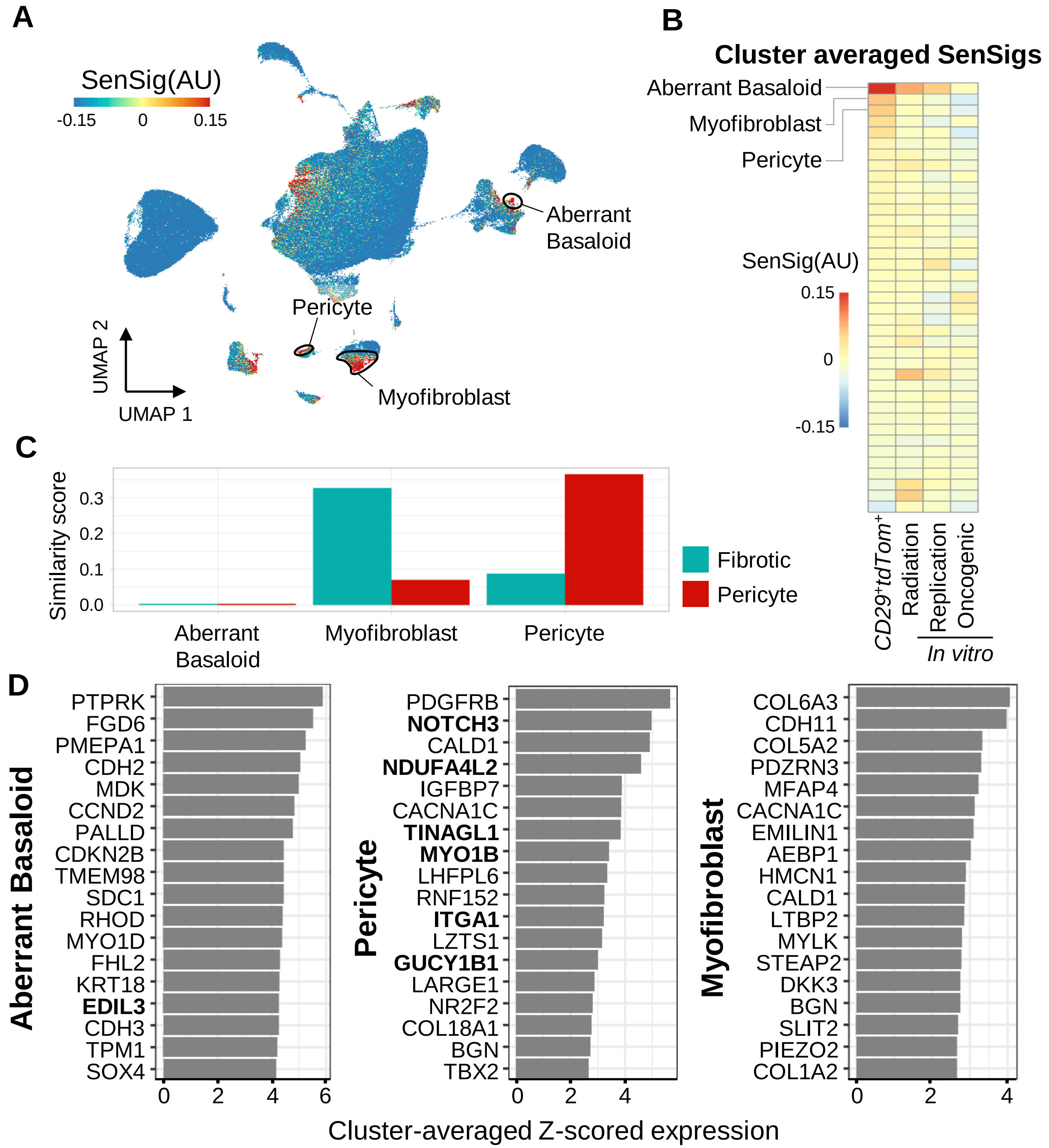
A
SenSig(AU)
-0.15
0
0.15
Aberrant Basaloid
Pericyte
Myofibroblast
UMAP 2
UMAP 1
B
Cluster averaged SenSigs
Aberrant Basaloid
Myofibroblast
Pericyte
SenSig(AU)
0.15
0
-0.15
CD29+tdTom+
Radiation
Replication
Oncogenic
In vitro
C
Similarity score
0.3
0.2
0.1
0.0
Aberrant Basaloid
Myofibroblast
Pericyte
Fibrotic
Pericyte
D
Aberrant Basaloid
PTPRK
FGD6
PMEPA1
CDH2
MDK
CCND2
PALLD
CDKN2B
TMEM98
SDC1
RHOD
MYO1D
FHL2
KRT18
EDIL3
CDH3
TPM1
SOX4
0
2
4
6
Pericyte
PDGFRB
NOTCH3
CALD1
NDUFA4L2
IGFBP7
CACNA1C
TINAGL1
MYO1B
LHFPL6
RNF152
ITGA1
LZTS1
GUCY1B1
LARGE1
NR2F2
COL18A1
BGN
TBX2
0
2
4
Myofibroblast
COL6A3
CDH11
COL5A2
PDZRN3
MFAP4
CACNA1C
EMILIN1
AEBP1
HMCN1
CALD1
LTBP2
MYLK
STEAP2
DKK3
BGN
SLIT2
PIEZO2
COL1A2
0
2
4
Cluster-averaged Z-scored expression

**Signaling targeting fibrotic SnC**

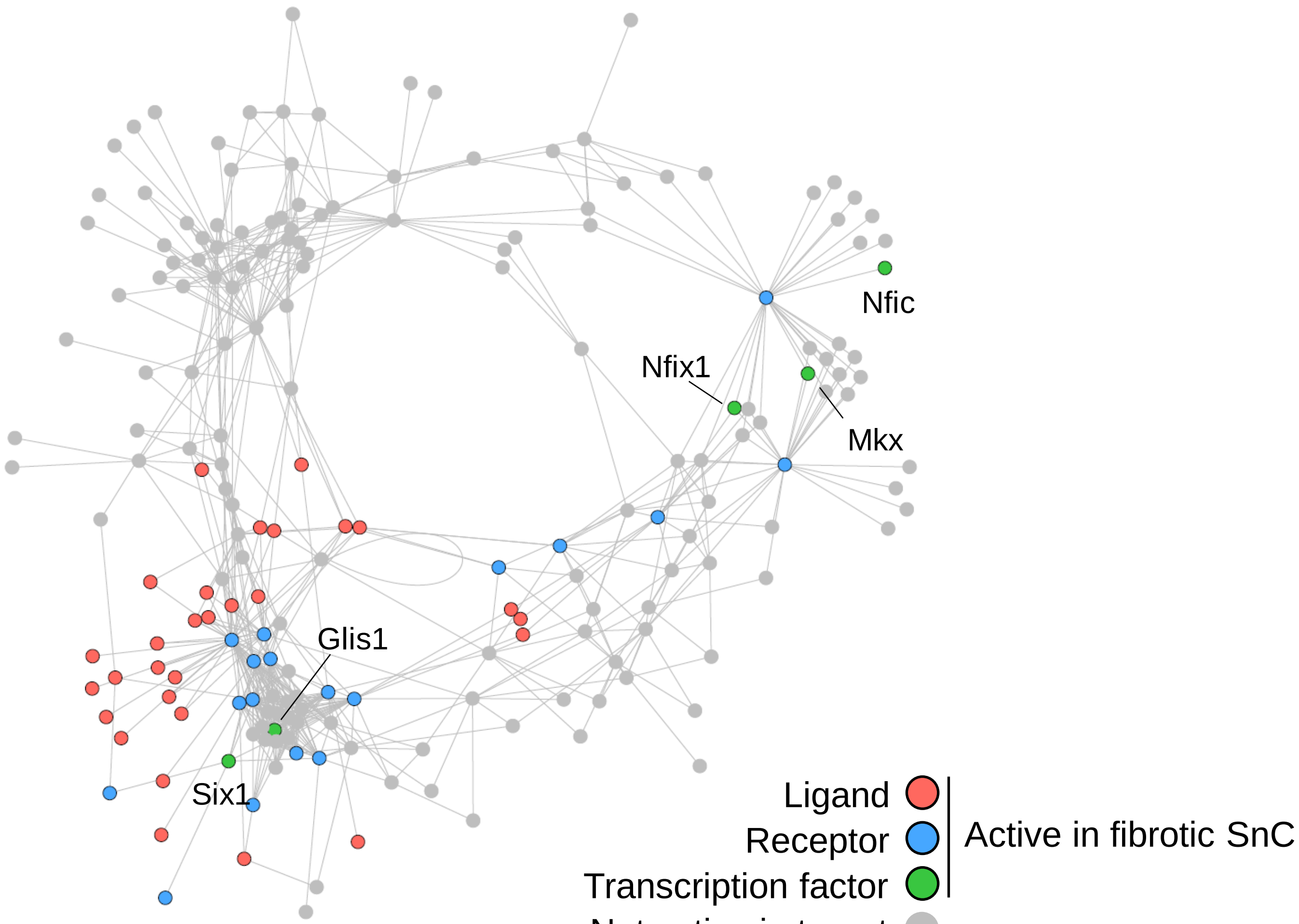

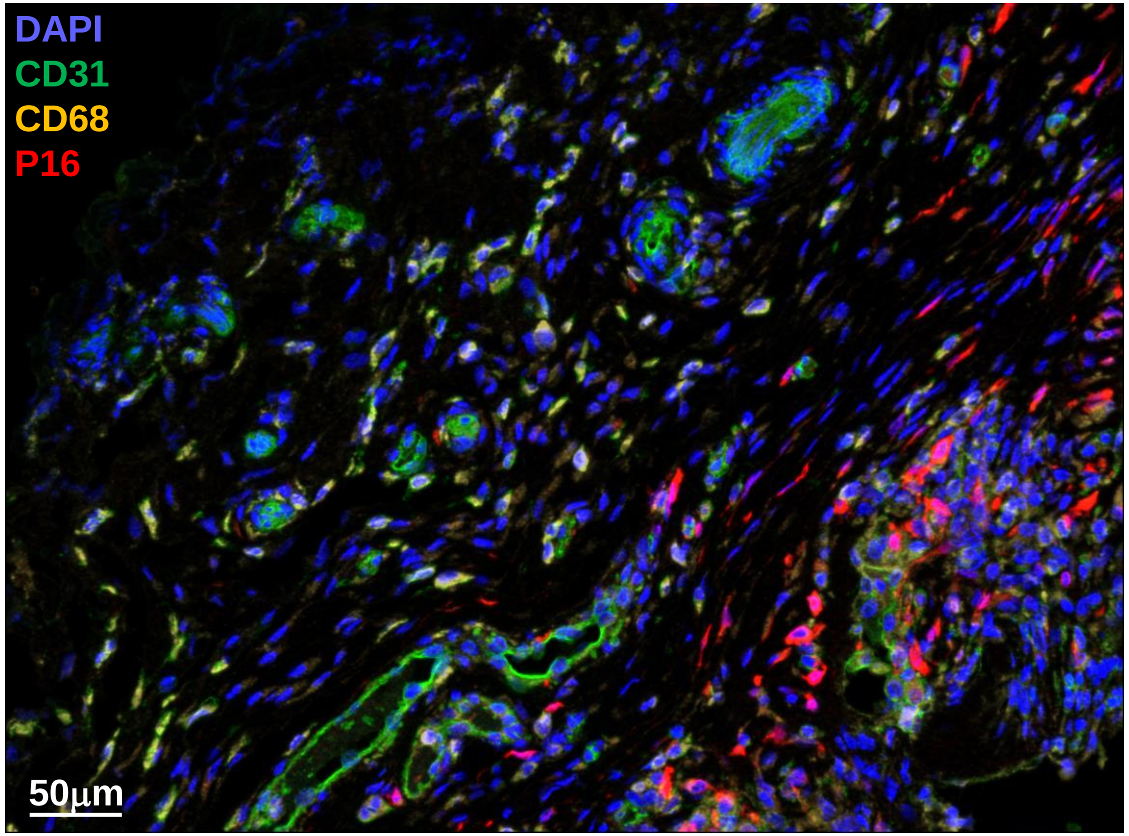
DAPI
CD31
CD68
P16
50µm
